# Regulation of sphingolipid synthesis by the C2H2 zinc finger transcription factor Com2 through ubiquitin-proteasome mediated degradation pathway

**DOI:** 10.64898/2026.02.20.707110

**Authors:** Kosei Matsumoto, Ayane Nagai, Nao Komatsu, Yuko Ishino, Rina Shirai, Toshiya Ueno, Mio Masaki, Ken-taro Sakata, Motohiro Tani, Tatsuya Maeda, Naotaka Tanaka, Mitsuaki Tabuchi

## Abstract

Membrane lipid synthesis is globally coordinated by a limited set of master transcription factors that regulate broad gene networks encoding lipid-metabolic enzymes and their regulators. Here, we identify the C2H2 zinc-finger transcription factor Com2 as a regulator of sphingolipid homeostasis in *Saccharomyces cerevisiae* that promotes transcription of downstream targets, including the protein kinase Ypk1, a key activator of sphingolipid synthesis. Com2 protein abundance increased upon treatment with myriocin, an inhibitor of sphingolipid synthesis, but rapidly decreased after addition of phytosphingosine (PHS), a precursor of complex sphingolipids; this decrease was blocked by proteasome inhibitors. These results suggest that Com2 is regulated in a sphingolipid-dependent manner through proteasome-mediated degradation. Moreover, a Com2 mutant in which lysine residues putatively involved in ubiquitination were replaced with arginine exhibited attenuated PHS-dependent degradation and elevated phosphorylation. Likewise, a mutant in which putative phosphorylation sites were replaced with alanine showed reduced PHS-dependent degradation. Together, these findings indicate that Com2 undergoes phosphorylation-dependent degradation via the ubiquitin–proteasome system in response to sphingolipid levels.

## Introduction

Cell membranes are composed of lipid bilayers, which consist of three major types of membrane lipids: glycerophospholipids, sphingolipids, and sterols (mainly cholesterol in mammals and ergosterol in yeasts) (van Meer et al., 2008). The synthesis of lipid bilayers during membrane formation involves the coordinated assembly of multiple lipid species, which requires the coordination of different levels, including lipid synthesis, uptake, metabolism, and subcellular distribution (Breslow and Weissman, 2010; Nohturfft and Zhang, 2009). This homeostasis is important for the maintenance of cellular and tissue functions, and its dysfunction in humans can lead to numerous diseases, such as cancer or type-2 diabetes (Harayama and Riezman, 2018). The principal instruments for the global control of lipid metabolism are regulated by master transcriptional regulators, which target genes encoding metabolic enzymes and/or their regulators working in specific metabolic pathways (Nohturfft and Zhang, 2009).

For example, in yeast, the transcription factors Ino2/Ino4 function in the expression of several glycerophospholipid synthases (Kwiatek et al., 2020). Under logarithmic growth or inositol-depleted conditions, the repressor Opi1 is bound to the endoplasmic reticulum (ER) membrane in a phosphatidic acid (PA)-dependent manner, and Ino2/Ino4 promote transcription. However, in steady-state or inositol-rich conditions, the decrease in PA on the ER membrane causes Opi1 to translocate to the nucleus, thereby inhibiting Ino2/Ino4-dependent transcription and repressing glycerophospholipid synthesis (Ambroziak and Henry, 1994; Loewen, 2003; Loewen et al., 2004; White et al., 1991). In the cholesterol metabolism of animal cells, the membrane-bound transcription factors, SREBPs, are synthesized as inactive precursors that are anchored in the membrane of the ER under sterol-rich conditions (Hua et al., 1996). SREBPs are translocated to the Golgi apparatus in a COPII vesicle-dependent manner (Nohturfft et al., 2000, 1999, 1998), where it is sequentially cleaved by Site-1 and Site-2 proteases, detached from the membrane (Rawson et al., 1997; Sakai et al., 1998, 1996; Wang et al., 1994), and translocated to the nucleus, thus promoting the expression of genes involved in cholesterol metabolism, such as HMG CoA reductase, the rate-limiting enzyme in cholesterol metabolism (Goldstein et al., 2006; Nohturfft and Zhang, 2009). Thus, in glycerophospholipid and sterol metabolism, it is well known that the synthesis of these lipids is coordinately regulated by master transcriptional regulators, in both yeast and mammals. However, master transcriptional regulators of sphingolipid metabolism remain elusive (Breslow and Weissman, 2010).

Sphingolipids and their metabolites play key cellular roles as both structural components of membranes and as signaling molecules that mediate responses to physiological cues (Breslow and Weissman, 2010). Recent studies using budding yeast, *Saccharomyces cerevisiae*, as a model system have improved our understanding of metabolic enzymes and how these enzymes are regulated to ensure sphingolipid homeostasis (Breslow et al., 2010; Breslow and Weissman, 2010). The regulation of sphingolipid metabolism by the target of rapamycin (TOR) complex 2 (TORC2)-Ypk1 pathway has been studied in detail in yeast (Berchtold et al., 2012; Eltschinger and Loewith, 2016; Niles et al., 2012; Tabuchi et al., 2006). As a regulatory mechanism of sphingolipid biosynthesis, TORC2, in concert with Slm1/2, which was identified as a synthetic lethal gene along with *MSS4* encoding a phosphatidylinositol 4-phosphate 5-kinase (Audhya et al., 2004; Tabuchi et al., 2006), phosphorylates Ypk1 upon depletion of complex sphingolipids at the plasma membrane (Berchtold et al., 2012; Niles et al., 2012). Phosphorylated Ypk1 further phosphorylates Orm1/2, which negatively regulates serine palmitoyl transferase (SPT) (Breslow et al., 2010; Gururaj et al., 2013; Roelants et al., 2011), which is the first step in sphingolipid metabolism. Thereafter, the phosphorylation of Orm1/2 through Ypk1 results in an increase in SPT activity. The TORC2-Ypk1 signal has also been reported to activate ceramide synthase by phosphorylating the N-terminal sites of the ceramide synthase catalytic subunits, Lag1/Lac1 (Muir et al., 2014). Together, TORC2-Ypk1-dependent signaling simultaneously upregulates the two most important steps in the sphingolipid biosynthetic pathway.

While investigating how reduced ceramide synthesis affects the TORC2-Ypk1 pathway using the ceramide synthase regulatory subunit mutant *lip1-1*, we found that *lip1-1* is hypersensitive to myriocin (Myr), a specific inhibitor of SPT (Figure 1A) (Ishino et al., 2022).

**Figure 1.**
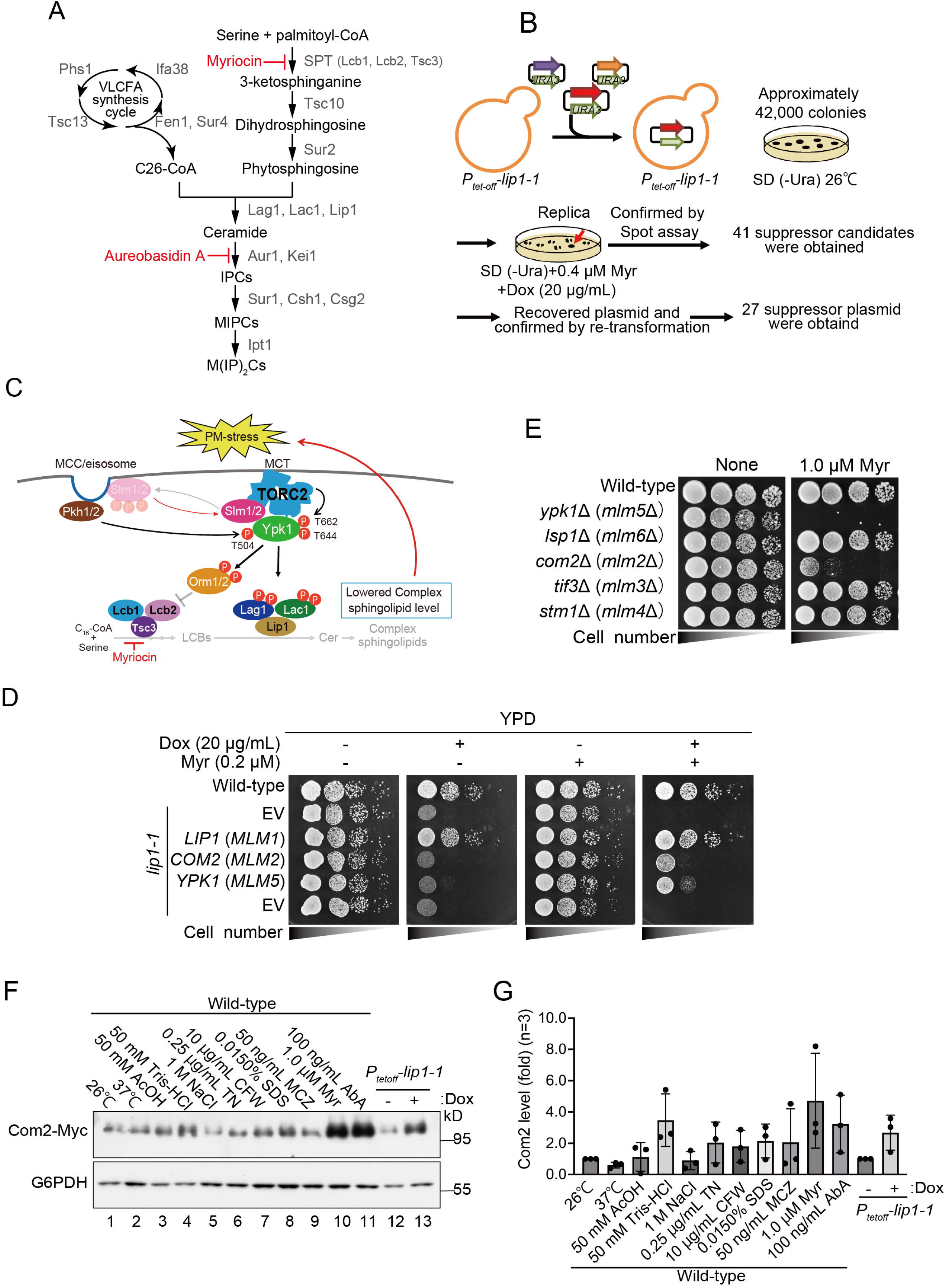
Screening for novel regulatory factors of sphingolipid metabolism. (**A**) *de novo* sphingolipid biosynthesis pathway in yeast, *Saccharomyces cerevisiae*. The pathway and proteins responsible for the synthesis of yeast sphingolipids are shown. IPC, inositol phosphorylceramide; MIPC, mannosylinositol phosphorylceramide; M(IP)_2_C, mannosyldiinositol phosphorylceramide. Myriocin (Myr) and aureobasidin A (AbA) inhibit the indicated steps of the sphingolipid biosynthesis pathway. (**B**) Schematic representation of screening for novel regulatory factors of sphingolipid metabolism by using the Myr-sensitive phenotype of *lip1-1* cells. (**C**) Schematic representation of the TORC2-Ypk1-dependent regulatory mechanism of sphingolipid biosynthesis. (**D**) Wild-type and *lip1-1* cells carrying an indicated plasmid were spotted at a 10-fold serial dilution on YPD supplemented with (0.2 µM) or without Myr, in the presence (20 µg/mL) or absence of doxycycline (Dox). (**E**) The diploid homozygous knockout cells of *MLM* genes (*ypk1*Δ, *lsp1*Δ, *com2*Δ, *tif3*Δ, and *stm1*Δ) were spotted at a 10-fold serial dilution on YPD supplemented with 1.0 µM Myr. (**F**, **G**) Wild-type (*COM2*-13Myc) and *P*_tetoff_*-lip1-1* cells were grown to mid-log in SD (+All) liquid medium in the presence (+) or absence (-) of Dox (20 µg/mL) and treated under the respective conditions for 3 hrs. Cells were then harvested and total cell lysates were prepared. The lysates were resolved by SDS-PAGE and immunoblotted with anti-Myc or anti-G6PDH antibodies, to detect Com2-13Myc or G6PDH (loading control), respectively. Western blot experiments were independently replicated three times. Values for Com2 expression level are expressed as mean ± SD (n≥3).

To identify novel regulators of sphingolipid metabolism, we performed a multicopy suppressor screen for genes that alleviate the Myr sensitivity of *lip1-1* cells. This approach led to the identification of Com2, a C2H2-type zinc finger transcription factor that had not previously been linked to sphingolipid metabolism. Further analyses revealed that Myr-induced sphingolipid depletion resulted in increased Com2 expression, accompanied by elevated Ypk1 expression and enhanced TORC2-dependent phosphorylation of Ypk1. In *com2*Δ cells, both the Myr-induced upregulation of Ypk1 and its activation by TORC2 were markedly reduced. Moreover, deletion of the Com2-binding site (CBS) in the *YPK1* promoter abolished these inductions, indicating that Ypk1 upregulation in response to sphingolipid depletion depends on Com2 and the CBS element in the *YPK1* promoter. We further found that Com2 is rapidly degraded via the ubiquitin-proteasome system (UPS) upon addition of exogenous sphingolipid precursor, demonstrating that Com2 abundance is primarily controlled through UPS-dependent proteolysis in a sphingolipid-dependent manner.

Collectively, these findings suggest that Com2 acts as a master transcription factor that senses intracellular sphingolipid levels, regulates CBS-dependent target genes such as *YPK1*, and is itself controlled by sphingolipid-dependent degradation via the ubiquitin–proteasome system.

## Results

### A screen for novel regulatory factors involved in sphingolipid metabolism

We previously generated and characterized a *P*_tet-off_*-lip1-1* (*lip1-1*) strain in which ceramide synthesis decreased in the presence of doxycycline (Dox) (Ishino et al., 2022). Dox-treated *lip1-1* cells are hypersensitive to myriocin (Myr), an inhibitor of serine palmitoyltransferase (SPT), the first step in sphingolipid biosynthesis (Figure 1A), presumably due to the synthetic effect of the Dox-dependent decrease in ceramide synthesis and inhibition of SPT. We hypothesized that overexpressing a protein involved in sphingolipid biosynthesis would confer Myr resistance to *lip1-1* cells. To identify novel regulatory factors, we transformed *lip1-1* cells with a multi-copy genomic library and selected transformants that grew on Myr-containing media (Figure 1B). Sequencing 27 Myr-resistant plasmids revealed 12 gene groups (*MLM* genes: *M*ulti-copy suppressor for *L*ip1-1 *M*yr-sensitive phenotype, Figure1—figure supplement 1, Table 1). Among these, the *YPK1* (*MLM5*), which encodes for a protein kinase downstream of the TORC2 signaling pathway, which is known to be involved in the regulation of sphingolipid metabolism (Figure 1C), was identified, thus validating that genes involved in the regulation of sphingolipid metabolism can be obtained employing this screening. Besides the *LIP1* (*MLM1*), the *COM2* (*MLM2*), which encodes a C2H2-type zinc finger transcription factor, was the most abundant (Figure 1D, Figure1—figure supplement 1, Table 1).

**Table 1.** Multi-copy suppressor for *lip1-1*-myriocin sensitivity (*MLM* genes)

| Suppressor | Number of clones isolated | Gene | Description* |
| --- | --- | --- | --- |
| *MLM1* | 8 | *LIP1* | Ceramide synthase subunit |
| *MLM2* | 7 | *COM2* | Transcriptional factor that binds IME1 Upstream Activation Signal (UAS)ru |
| *MLM3* | 2 | *TIF3* | Transcription initiation factor eIF-4B |
| *MLM4* | 2 | *YLR149C-A* | Dubious open reading frame |
|  |  | *STM1* | Protein required for optimal translation under nutrient stress |
| *MLM5* | 1 | *YPK1* | Ser/Thr protein kinase |
| *MLM6* | 1 | *LSP1* | Eisosome core component |
| *MLM7* | 1 | *BMH2* | 14-3-3 protein |
| *MLM8* | 1 | *RIM20* | Protein involved in proteolytic activation of Rim101p |
|  |  | *CAF20* | Phosphoprotein of the mRNA cap-binding complex |
|  |  | *YOR277C* | Dubious open reading frame; almost completely overlaps the verified gene CAF20 |
|  |  | *SNR31* | H/ACA box small nuclear RNA (snoRNA) |
|  |  | *SNR5* | H/ACA box small nuclear RNA (snoRNA) |
|  |  | *HEM4* | Uroporphyrinogen III synthase |
|  |  | *RFM1* | Component of the Sum1p-Rfm1p-Hst1p complex |
| *MLM9* | 1 | *MOT3* | Transcriptional repressor, activator |
|  |  | *TVP18* | Integral membrane protein |
|  |  | *ABF2* | Mitochondrial DNA-binding protein |
| *MLM10* | 1 | *SRO77* | Protein with roles in exocytosis and cation homeostasis |
|  |  | *PKC1* | Ser/Thr protein kinase |
| *MLM11* | 1 | *PIN4* | Protein involved in G2/M phase progression and response to DNA damage |
|  |  | *SEC17* | Alpha-SNAP cochaperone |
| *MLM12* | 1 | *SUT2* | Zn2Cys6 family transcription factor |

\**Saccharomyces* GENOME DATABASE: https://www.yeastgenome.org/

To clarify whether the obtained *MLM* genes are involved in the regulation of sphingolipid metabolism, we examined the Myr sensitivity of each gene-knockout homozygous diploid strain. Upon doing so, we found that *com2*Δ, as well as *ypk1*Δ, exhibited Myr sensitivity, while the other *MLM* gene-deficient strains did not (Figure 1E, Figure 1—figure supplement 2).

We further examined the phenotype of *com2*Δ in response to various drugs and stress conditions in addition to Myr. Our results showed that *com2*Δ exhibited higher sensitivity to Myr compared to the wild-type. However, it did not display sensitivity to aureobasidin A (AbA), an inhibitor of inositol phosphoceramide (IPC) synthesis, ergosterol synthesis inhibitors such as miconazole (MCZ) and terbinafine, or any other drugs and stress conditions tested (Figure1—figure supplement 3A-C). Unlike *ypk1*Δ, *com2*Δ does not exhibit sensitivity to AbA. This observation suggests that Com2 plays a more specialized role in regulating sphingolipid biosynthesis at an earlier, upstream step.

Next, we analyzed the expression levels of Com2 in wild-type cells under various drug and stress treatment conditions. We observed a remarkable upregulation of Com2 expression under conditions that reduced complex sphingolipid synthesis, such as in Myr-treated cells, AbA-treated cells or Dox-treated *lip1-1* cells (Figure 1F, G). In contrast, inhibition of sterol synthesis did not lead to a significant increase in Com2 expression (Figure 1—figure supplement 3D, E). These findings indicate that Com2 expression is specifically upregulated in response to a reduction in complex sphingolipid levels.

### Com2 functions upstream of TORC2-Ypk1 pathway

Overexpression of Com2 or Ypk1 suppressed the Myr sensitivity of *lip1-1* (Figure 1D) and deletion of any of these genes conferred Myr sensitivity (Figure 1E), suggesting that these two proteins function in the same pathway involved in the regulation of sphingolipid metabolism. Com2 is a C2H2-type zinc finger transcription factor consisting of 443 amino acid residues with two zinc fingers at its C-terminus. In addition to that, several putative phosphorylation sites for AGC kinases, including Ypk1, were found in its amino acid sequence (Figure 2A) (Muir et al., 2014). Based on these observations, we hypothesized that Com2 is involved in sphingolipid metabolism downstream of Ypk1 and that Ypk1 impacts the phosphorylation and function of Com2.

**Figure 2.**
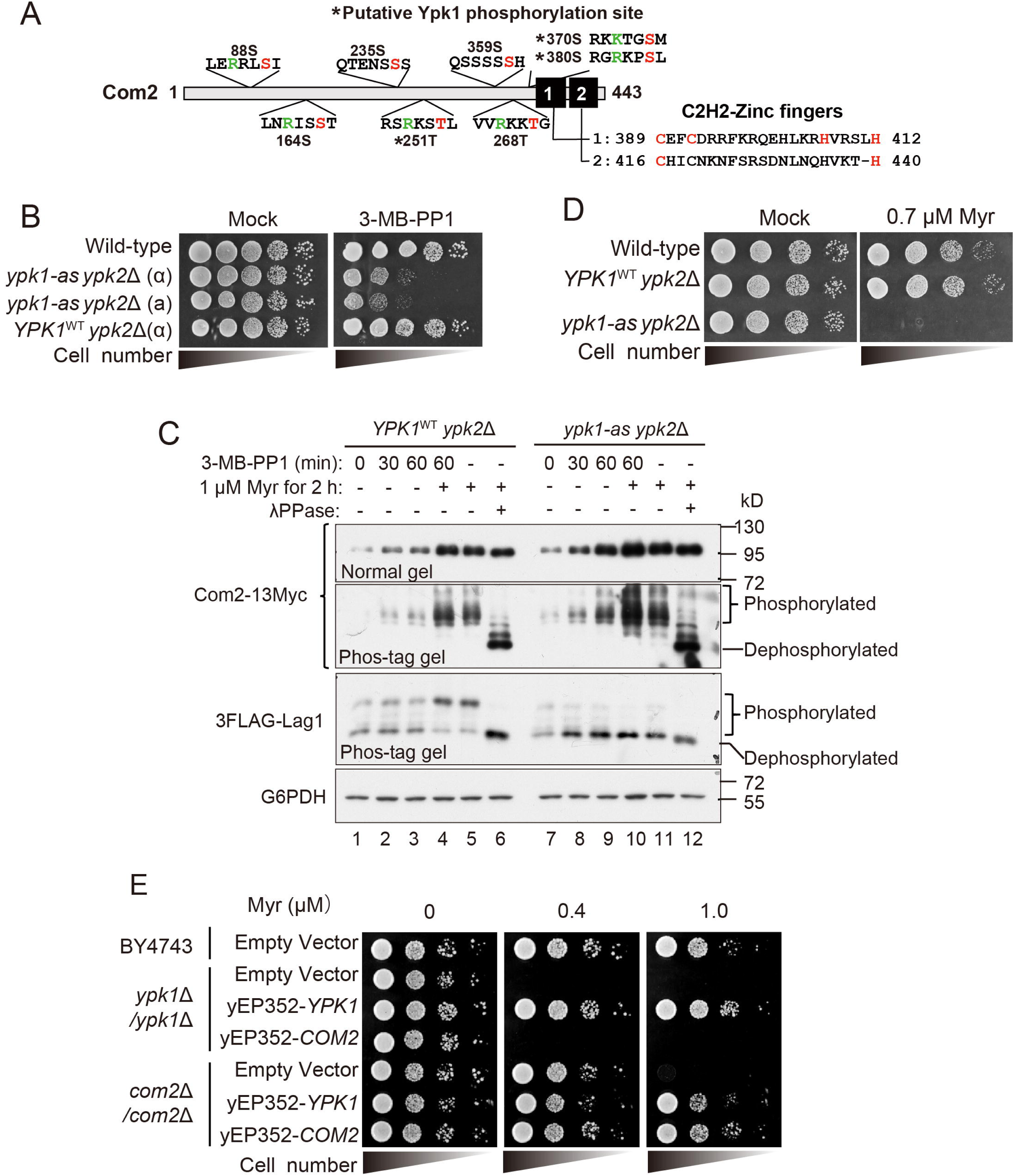
Com2 functions upstream of the TORC2-Ypk1 pathway. (**A**) Schematic representation of Com2, with the positions of the predicted AGC-kinase-dependent phosphorylation sites and two Zinc-finger domains highlighted. (**B**) Characterization of *ypk1-as ypk2*Δ mutant. Wild-type, the ATP-analogue sensitive allele *ypk1*^L424A^, *ypk1-as ypk2*Δ cells and the wild-type allele of Ypk1, *YPK1^WT^ ypk2*Δ cells were spotted on YPD containing DMSO (Mock) or indicated concentration of 3MB-PP1 and incubated for 3 days at 26 °C. *ypk1-as ypk2*Δ cells exhibit growth defect by limiting of Ypk1-activity by addition of 3MB-PP1. (**C**) Wild-type cells, *YPK1*^WT^ *ypk2*Δ cells and *ypk1-as ypk2*Δ cells are spotted on YPD and YPD containing 0.7 μM myriocin and incubated for 2 days at 26 °C. *ypk1-as ypk2*Δ cells exhibit Myr-sensitive without inhibition by addition of 3MB-PP1. (**D**) Wild-type *YPK1*^WT^ *ypk2*Δ cells and analogue-sensitive mutant *YPK1*^L424A^*ypk2*Δ cells carrying the plasmid expressing Com2-13Myc were treated with 3MB-PP1 for the indicated time-periods and then treated with 1.0 µM Myr for 2 hrs. The lysates were then resolved using phos-tag or normal SDS-PAGE and immunoblotted with anti-Myc, anti-FLAG, or anti-G6PDH antibodies, to detect Com2-13Myc, 3FLAG-Lag1, or G6PDH (loading control), respectively. (**E**) Wild-type (BY4743) and knockout homozygote diploid cells of *ypk1*Δ or *com2*Δ carrying an indicated multi-copy plasmid were spotted at a 10-fold serial dilution on YPD supplemented with the indicated concentrations of Myr.

To test this hypothesis, we investigated the effect of limiting Ypk1 activity on the phosphorylation state of Com2. To this end, we employed *ypk1-as ypk2*Δ cells, which express an ATP analog-sensitive (AS) allele, Ypk1(L424A), and titrated down its activity by addition of an efficacious inhibitor, 1-(tert-butyl)-3-(3-methylbenzyl)-1H-pyrazole[3,4-d]pyrimidin-4-amoine (3-MB-PP1), which has no effect on the wild-type cells (Figure 2B) (Muir et al., 2014). As previously reported (Ishino et al., 2022; Muir et al., 2014), Myr treatment stimulated the phosphorylation of the 3FLAG-tagged Lag1 in wild-type cells, as verified using Phos-tag SDS-PAGE (Figure 2C, lanes 1-5 in the 3FLAG-Lag1 blot), but this effect was abolished in *ypk1-as ypk2*Δ cells (Figure 2C, lanes 7-11 in the 3FLAG-Lag1 blot), confirming the inhibition of Ypk1 activity. It is noted that the *ypk1-as ypk2*Δ mutant did not show Myr-dependent phosphorylation of Lag1 even in the absence of 3-MB-PP1 (Figure 2C, lane 11 in 3FLAG-Lag1 blot) and was also sensitive to Myr (Figure 2D), suggesting that this mutant appears to be a hypomorphic allele and treatment of 3-MB-PP1 completely blocks Ypk1 activity. In contrast to phosphorylation of Lag1, a 13Myc epitope-tagged Com2 was highly phosphorylated in both the wild-type and *ypk1-as ypk2*Δ cells (Figure 2C, lanes 1-5 and lanes 7-11 in the Com2-13Myc blot of Phos-tag gel), indicating that the phosphorylation state of Com2 is not dependent on Ypk1 activity. Notably, as shown in Figure 1F, G, Myr treatment increased Com2 expression in wild-type cells, and this effect was more pronounced in *ypk1-as ypk2*Δ cells treated with 3-MB-PP1.

Next, we performed an epistatic analysis between *COM2* and *YPK1,* confirming that overexpressing each gene suppressed Myr sensitivity in the corresponding deletion strain. Overexpression of *COM2* did not suppress the Myr sensitivity of *ypk1*Δ cells, whereas overexpression of *YPK1* suppressed the Myr sensitivity of *com2*Δ cells (Figure 2E), indicating that Com2 functions upstream of Ypk1.

Taken together, these results suggested that Com2 functions upstream of the TORC2-Ypk1 pathway rather than downstream of it.

### Com2 expression increases in response to a decrease in sphingolipids

Epistasis analysis suggested that *COM2* functions upstream of *YPK1* (Figure 2E). We noticed a putative Com2-binding site (CBS; 5′-CCCTAT-3′) exists in the *YPK1* promoter (Siggers et al., 2014). Based on these observations, we hypothesized that Com2 regulates *YPK1* expression in a sphingolipid-dependent manner (Figure 3A).

**Figure 3.**
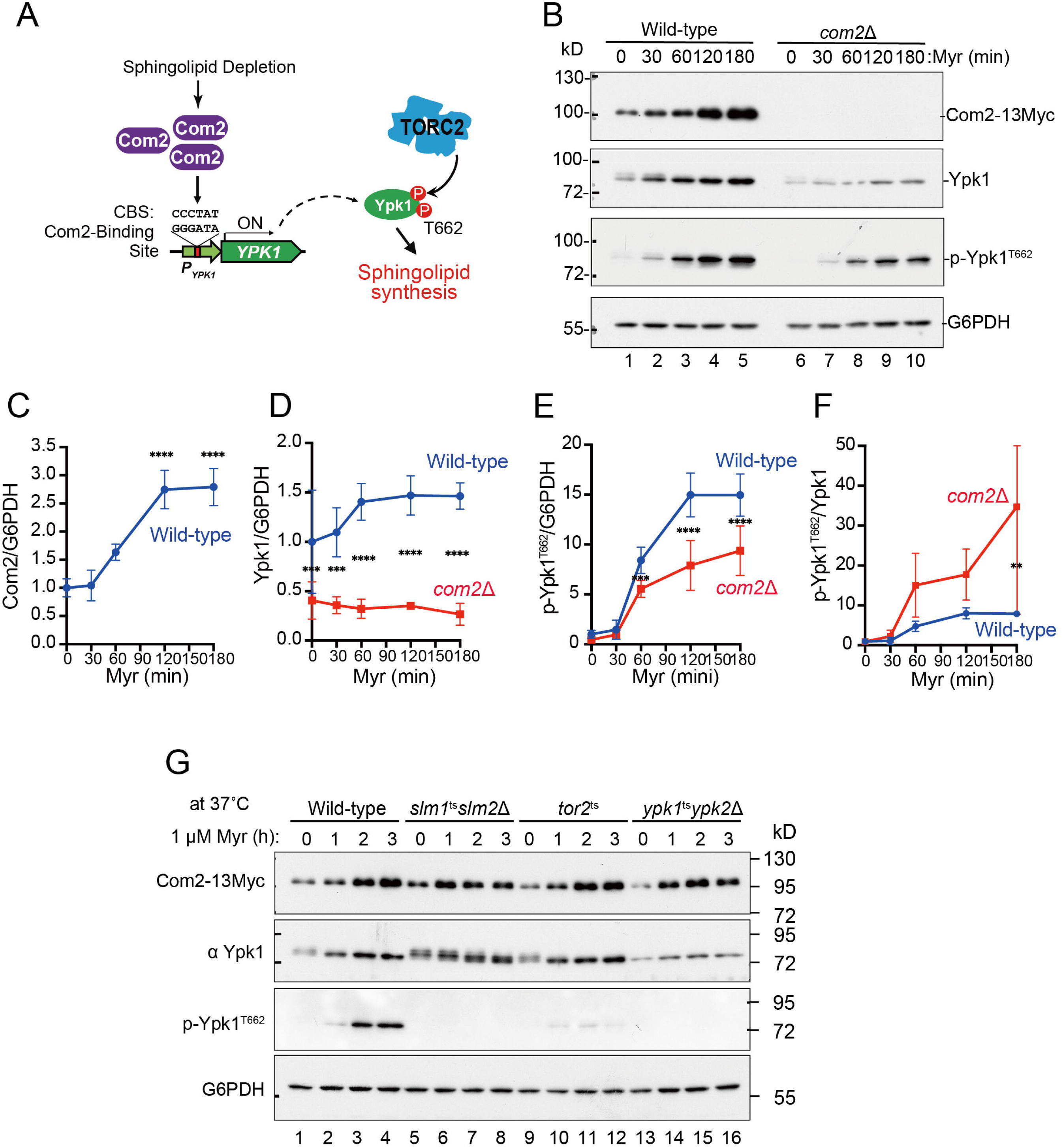
Ypk1 is expressed in a Com2-dependent manner. (**A**) Proposed model for the mechanism by which sphingolipid depletion increases Com2 expression, induces *YPK1* transcription, and activates the TORC2-Ypk1 pathway to promote sphingolipid biosynthesis. (**B**) Wild-type and *com2*Δ cells were grown to mid-log phase in SD liquid medium and then treated with 1.0 µM Myr. The cells were then harvested, and total cell lysates were prepared from them. The lysates were resolved using SDS-PAGE and immunoblotted with anti-Myc, anti-Ypk1, anti-phospho-Ypk1^T662^, or anti-G6PDH antibodies, to detect Com2-13Myc, Ypk1, TORC2-dependent phosphorylation site (hydrophobic motif) of Ypk1 (p-Ypk1^T662^) or G6PDH (loading control), respectively. (**C**) The time-dependent increase in Com2 expression following Myr treatment was quantified in wild-type cells. Experiments were performed three times independently. Com2 levels were normalized to the internal control G6PDH, with the value at 0 min of Myr treatment set to 1. Data are presented as the mean ± SD from three independent experiments. Asterisks indicate statistically significant differences compared with the 0 min time point as determined by one-way ANOVA followed by Tukey’s multiple comparisons test (\*\*\*\**p*<0.0001). (**D, E**) To examine time-dependent changes in Ypk1 or phosphorylated Ypk1 at T662 (p-Ypk1^T662^) in wild-type and *com2*Δ cells, the levels of Ypk1 or p-Ypk1^T662^ at each time point after Myr treatment were normalized to the loading control G6PDH and then further normalized to the value at 0 min before Myr treatment in wild-type cells (set to 1). Data are presented as the mean ± SD from three independent experiments. Asterisks indicate statistically significant differences as determined by two-way ANOVA followed by Sidak’s multiple comparisons test (\*\**p*<0.01, \*\*\**p*<0.001, \*\*\*\**p*<0.0001). (**F**) To examine time-dependent changes in phosphorylated Ypk1 at T662 (p-Ypk1^T662^) in wild-type and *com2*Δ cells, the levels of p-Ypk1^T662^ at each time point after Myr treatment were normalized to total Ypk1 and then further normalized to the value at 0 min before Myr treatment in wild-type cells (set to 1). Data are presented as the mean ± SD from three independent experiments. Asterisks indicate statistically significant differences as determined by two-way ANOVA followed by Sidak’s multiple comparisons test (\*\**p*=0.0092). (**G**) Wild-type, *slm1*^ts^*slm2*Δ, *tor2*^ts^, and *ypk1*^ts^*ypk2*Δ cells were grown to mid-log phase in SD liquid medium at 26°C and then shifted to 37°C for 30 min and treated with 1.0 μM Myr. The cells were then harvested at the indicated time-point and total cell lysates were prepared from them. The lysates were resolved using SDS-PAGE and immunoblotted with anti-Ypk1, anti-Myc, anti-G6PDH antibodies, or anti-phospho-Ypk1^T662^ antibodies, to detect Ypk1, Com2-13Myc, G6PDH (loading control), or TORC2-dependent phosphorylation of Ypk1 (p-Ypk1^T662^), respectively.

To test this hypothesis, we performed a time-course analysis of Com2 and Ypk1 during Myr treatment in wild-type and *com2*Δ cells and assessed protein levels by immunoblotting. In wild-type cells, Myr-treatment caused a time-dependent increase in Com2, accompanied by an increase in total Ypk1 as well as enhanced TORC2-dependent phosphorylation of the Ypk1 hydrophobic motif (p-Ypk1^T662^) (Figure 3B, lanes 1-5, Figure 3C-E). In contrast, in *com2*Δ cells, both the upregulation of Ypk1 and the Myr-dependent increase in p-Ypk1^T662^ were strongly attenuated (Figure 3B, lanes 6–10, Figure 3D, E), indicating that these responses depend on Myr-induced Com2 expression. Notably, TORC2 activity was not reduced in *com2*Δ cells (if anything, it appeared elevated) (Figure 3F), suggesting that the diminished p-Ypk1^T662^ signal primarily reflects decreased Ypk1 abundance rather than defective TORC2 activation.

TORC2 senses plasma membrane stress caused by a reduction in complex sphingolipids through the eisosome-associated adaptor proteins, Slm1/2, thereby activating Ypk1 to stimulate sphingolipid synthesis (Figure 1C) (Berchtold et al., 2012). Next, we investigated the contribution of the Slm-TORC2-Ypk1 pathway to the Myr-induced expression of Com2. We observed Myr-induced expression of Com2 even after blocking of the Slm-TORC2-Ypk1 pathway at non-permissive temperatures using each temperature-sensitive mutant (Figure 3G), indicating that the induction of Com2 expression upon decreased complex sphingolipids is due to a sensing pathway that is independent from the Slm-TORC2-Ypk1 pathway.

Taken together, we demonstrated that Com2 is upregulated in response to a decrease in sphingolipid synthesis, accompanied by upregulation of Ypk1 expression and its TORC2-dependent phosphorylation (Figure 3A).

### Overexpression of Com2 causes increased expression of Ypk1 via a putative CBS in the *YPK1* promoter

To further substantiate that Com2 acts upstream of Ypk1, we replaced the endogenous *COM2* promoter with a tetracycline-repressible tet-off promoter by homologous recombination, generating a *P*_tet-off_*-GFP-COM2* strain in which Com2 expression can be controlled by Dox (Figure 4A). Using this strain, we assessed Myr sensitivity under conditions of GFP-Com2 overexpression or repression. Cells overexpressing GFP-Com2 (-Dox) were more resistant to Myr than wild-type cells (Figure 4B, upper panel), whereas repression of GFP-Com2 (+Dox) rendered cells as Myr-sensitive as *com2*Δ cells (Figure 4B, lower panel).

**Figure 4.**
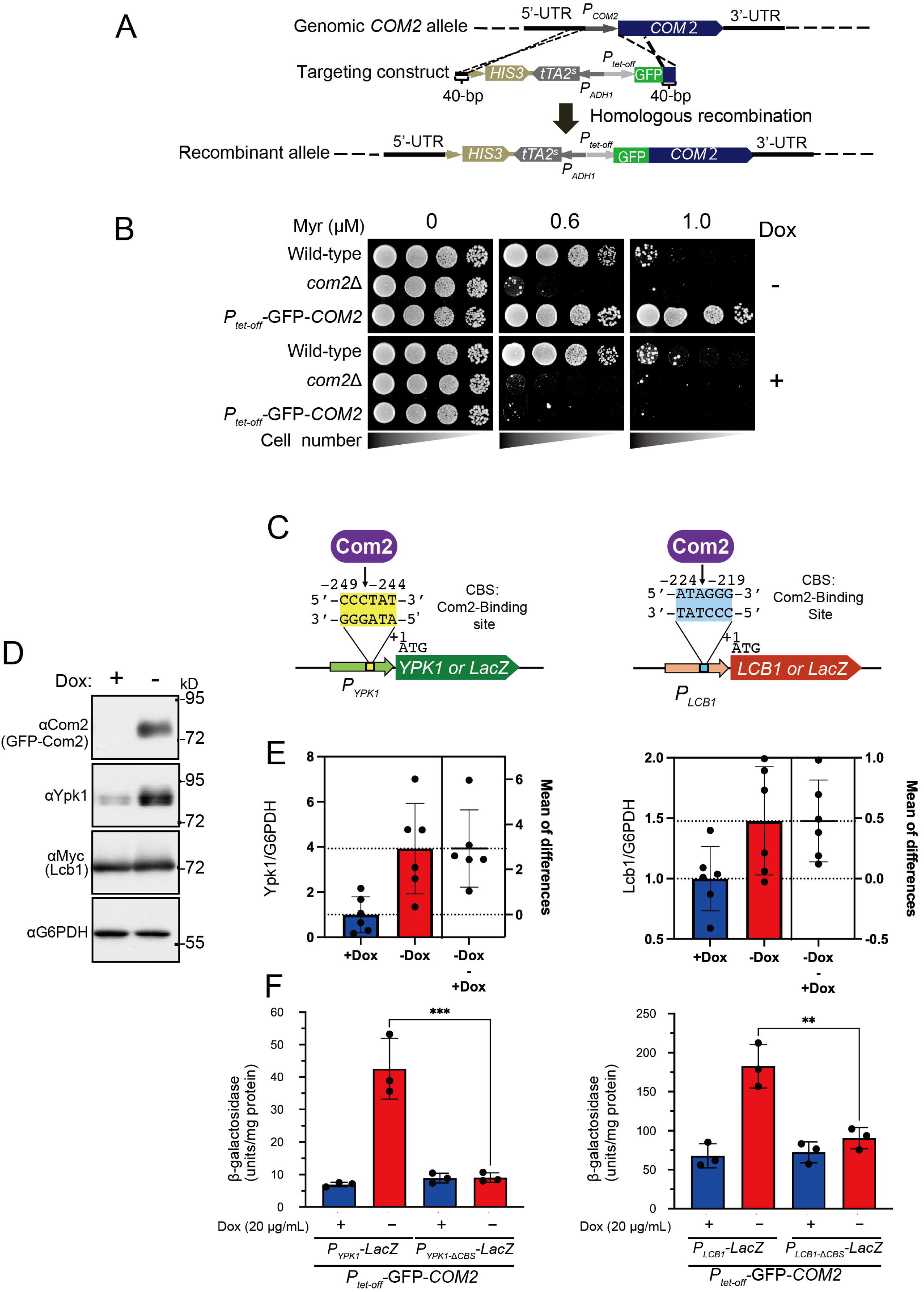
Overexpression of Com2 causes increased expression of Ypk1. (**A**) The chromosomal *COM2* promoter (*P_COM2_*) was replaced by a Tet-off promoter and GFP was fused to the N-terminus of Com2, to generate the *P*_tet-off_-GFP-*COM2* strain. (**B**) Wild-type, *com2*Δ, and *P*_tet-off_-GFP-*COM2* cells were spotted at a 10-fold serial dilution on YPD supplemented with 0.6 or 1.0 µM Myr, in the presence (20 µg/mL) (+) or absence (-) of Dox. (**C**) *YPK1* and *LCB1* possess a putative Com2-binding site (CBS) in the promoter region. *LacZ* reporter plasmids containing the wild-type or Com2-binding site (CBS)-deleted promoter regions of *YPK1* or *LCB1* were used to assess Com2-dependent transcriptional regulation. (**D**) *P*_tet-off_*-GFP-COM2* cells were grown to mid-log phase in SD liquid medium in the presence (+) or absence (−) of doxycycline (Dox) and then treated with 1.0 µM Myr for 2 hrs. Cell lysates were subjected to SDS-PAGE followed by immunoblotting with anti-Com2, anti-Ypk1, anti-Myc, or anti-G6PDH antibodies to detect GFP-Com2, Ypk1, Lcb1-13Myc, or G6PDH (loading control), respectively. A representative immunoblot is shown. The full set of immunoblots from six independent experiments is provided in Figure 4—figure supplement 2. (**E**) Quantification of Ypk1 and Lcb1 protein levels shown in Figure 4—figure supplement 2. Band intensities were quantified using ImageJ. Data are presented as the mean ± SD from six independent experiments. Statistical significance was assessed using paired t-test (Ypk1, *p*=0.0071; Lcb1, *p*=0.0151). The right panel indicates paired differences with mean difference and 95% confidence intervals. (**F**) *P*_tet-off_-GFP-*COM2* cells carrying p*P_YPK1_-LacZ*, p*P_YPK1CBS_*_Δ_*-LacZ* or p*P_LCB1_-LacZ*, p*P_LCB1CBS_*_Δ_*-LacZ* plasmid were grown to mid-log phase in SD liquid medium, in the presence (+) or absence (-) of Dox. Lysates were obtained from the indicated samples and assayed for β-galactosidase activity. Data are presented as the mean ± SD from three independent experiments. \*\*\**p*<0.001, as assessed using one-way ANOVA, with Tukey multiple comparison tests.

A putative Com2-binding site (CBS) was identified in the *YPK1* promoter (Figure 4C). In addition, a genome-wide search of the *S. cerevisiae* genome using the CBS consensus motif revealed a matching sequence in the promoter of *LCB1*, which encodes a catalytic subunit of SPT (Figure 4C; Figure 4—figure supplement 1). To test whether Com2 directly regulates *YPK1* and *LCB1* expression, we examined the effects of GFP-Com2 overexpression on their expression levels. Induction of GFP-Com2 overexpression (-Dox) markedly increased Ypk1 abundance, and also led to a significant, albeit more modest, increase in Lcb1 abundance (Figure 4D, E, Figure 4—figure supplement 2).

We next constructed *lacZ* reporters driven by the *YPK1* or *LCB1* promoter (Figure 4C) and assessed promoter activity by measuring β-galactosidase activity. Compared with conditions in which GFP-Com2 was repressed (+Dox), GFP-Com2 induction (-Dox) increased β-galactosidase activity ∼6-fold for the *YPK1* promoter and ∼2.5-fold for the *LCB1* promoter (Figure 4F). Notably, this Com2-dependent activation was abolished by deleting the CBS from each promoter, indicating that Com2 directly stimulates *YPK1* and *LCB1* transcription via the CBS motif. Collectively, these findings support a model in which Com2 contributes to sphingolipid metabolism by transcriptionally upregulating *YPK1* and *LCB1*.

### Com2 promotes sphingolipid synthesis, partly through the expression of Ypk1

To test whether Com2 directly regulates *YPK1* transcription and how this regulation impacts sphingolipid metabolism, we used CRISPR-Cas9 genome editing to delete the putative Com2-binding site (CBS) from the endogenous *YPK1* promoter, generating *P_YPK1_-*Δ*CBS* cells (Figure 5A).

**Figure 5.**
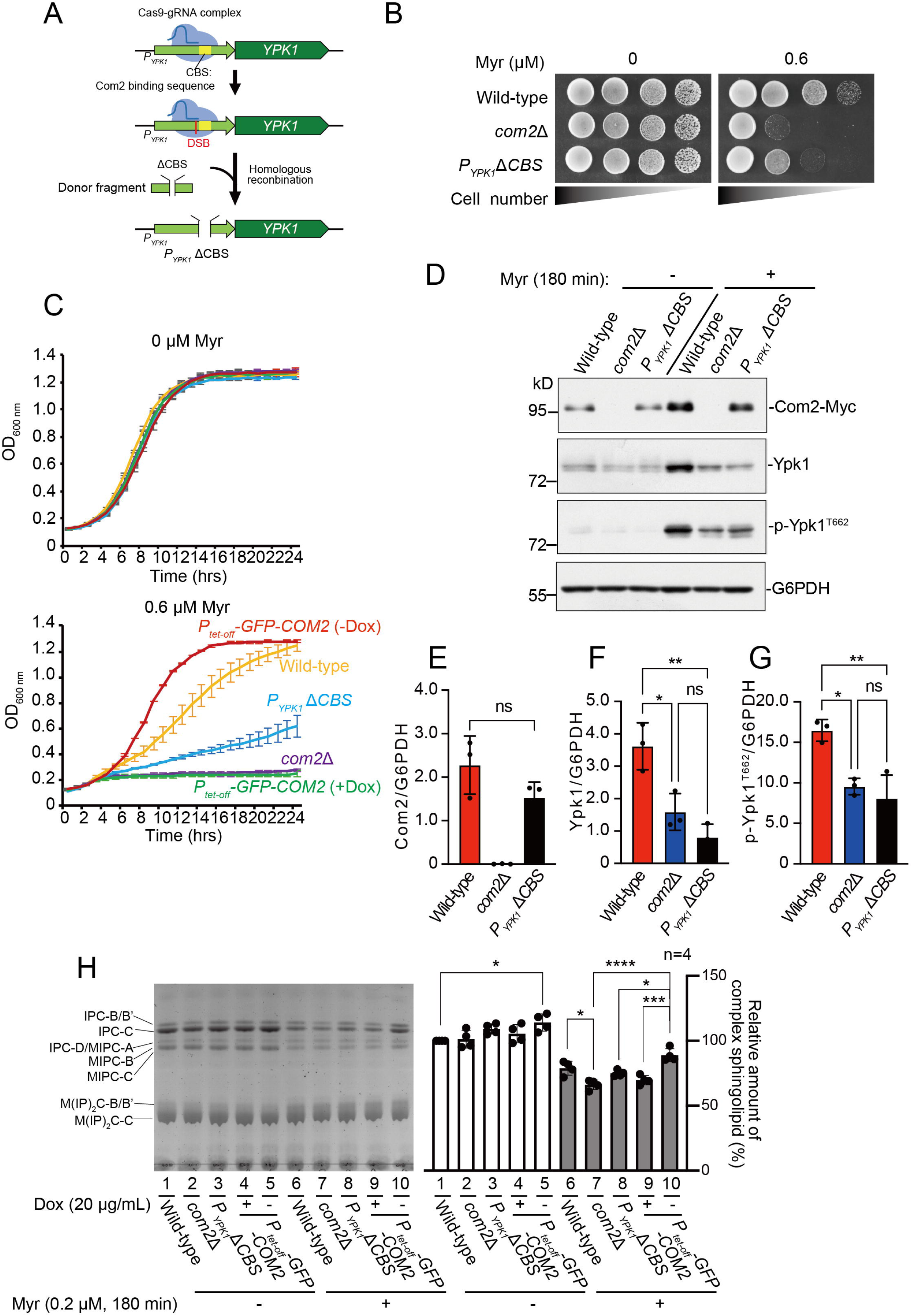
The Com2-binding site in the *YPK1* promoter is required for both proper expression and TORC2-dependent phosphorylation of Ypk1. (**A**) A schematic representation of the generation of the *P_YPK1_*-Com2 binding site–deleted (*P_YPK1_*_-_Δ*CBS*) strain using CRISPR-Cas9-mediated genome editing. The Cas9-gRNA complex introduces a double-strand break near the putative Com2 binding site (CBS) in the chromosomal *YPK1* promoter (*P_YPK1_*). A donor DNA fragment lacking the CBS (*P_YPK1_*ΔCBS) is simultaneously introduced and integrated into the chromosome by homologous recombination, resulting in deletion of the CBS from the endogenous *YPK1* promoter. (**B**) Wild-type, *com2*Δ, and *P_YPK1_-CBS*Δ cells were spotted at a 10-fold serial dilution on YPD supplemented with or without 0.6 µM Myr. (**C**) Wild-type, *com2*Δ, *P_YPK1_*Δ*CBS*, and *P*_tet-off_-GFP-*COM2* cells were grown to the exponential phase, diluted to OD_600_=0.1, and grown in YPD liquid medium treated with or without 0.6 µM Myr, in the absence (-) or presence (+) of Dox (20 µg/mL), in a microtiter plate. (**D, E, F, G**) The indicated cells were grown to mid-log phase in SD liquid medium and treated with 1.0 µM Myr at 26°C for 3 hrs. Cells were then harvested and whole cell lysates were prepared. Whole cell lysates were separated by SDS-PAGE and immunoblotted with anti-Myc, anti-Ypk1, anti-phospho-Ypk1^T662^ or anti-G6PDH antibodies to detect Com2-13Myc, Ypk1, p-Ypk1^T662^ or G6PDH (loading control), respectively. The amounts of Com2-13Myc, Ypk1, and p-Ypk1^T662^ in wild-type cells (containing Myr) were normalized by the amount of G6PDH. Data represents the mean ± SD of three independent experiments. Statistical significance was determined by one-way ANOVA followed by Tukey’s multiple comparison test (\**p*<0.05, \*\**p*<0.01, ns: no significant difference). (**H**) TLC analysis of sphingolipids extracted from wild-type, *com2*Δ, *P_YPK1_*-*CBS*Δ, and *P*_tet-off_-GFP-*COM2* cells. Cells were grown to mid-log phase in YPD liquid medium, in the presence (+) or absence (-) of Dox (20 µg/mL) and treated with or without 0.2 µM Myr for 3 hrs. Complex sphingolipids were analyzed using TLC (left). The level of total complex sphingolipids in the wild-type cells was taken as 100% and each value is displayed as a graph (right). The data have been represented as mean ± SD of four independent experiments. Statistical significance was determined by one-way ANOVA followed by Tukey’s multiple comparison test (\**p*<0.05, \*\*\**p* =0.0002, and \*\*\*\**p*<0.0001).

We first assessed the Myr sensitivity of *P_YPK1_-*Δ*CBS* cells. *P_YPK1_-*Δ*CBS* cells were more sensitive to Myr than wild-type cells, but less sensitive than *com2*Δ cells or doxycycline-treated *P*_tet-off_*-GFP-COM2* cells (+Dox) (Figure 5B, C).

We next examined Ypk1 expression in *P_YPK1_-*Δ*CBS* cells by western blotting. Under Myr-treated conditions, Com2-13Myc levels were comparable between wild-type and *P_YPK1_-*Δ*CBS* cells, whereas Ypk1 levels were reduced in *P_YPK1_-*Δ*CBS* cells relative to wild-type and were similar to those in *com2*Δ cells (Figure 5D–F). Consistently, TORC2-dependent phosphorylation at the hydrophobic motif (p-Ypk1^T662^) was decreased in *P_YPK1_-*Δ*CBS* cells compared with wild-type and was comparable to that in *com2*Δ cells (Figure 5D, G).

We then performed TLC analysis to examine how Com2-dependent regulation influences sphingolipid synthesis. Consistent with the Myr resistant phenotype, overexpression of GFP-Com2 (-Dox) significantly increased sphingolipid levels under both untreated (Figure 5H, lane 1 vs. lane 5) and Myr-treated conditions (Figure 5H, lane 6 vs. lane 10). In contrast, under Myr treatment, sphingolipid levels were substantially reduced in *com2*Δ cells and in GFP-Com2-repressed (+Dox) cells compared with wild-type (Figure 5H, lane 6 vs. lane 7 or 9). In Myr-treated *P_YPK1_-*Δ*CBS* cells, sphingolipid levels were significantly lower than in GFP-Com2-overexpressing (-Dox) cells (Figure 5H, lane 8 vs. lane 10), but only slightly reduced compared with wild-type (Figure 5H, lane 6 vs. lane 8), consistent with the intermediate Myr sensitivity of *P_YPK1_-*Δ*CBS* cells (Figure 5B).

Notably, the double mutant *P_YPK1_-*Δ*CBS* and *P_LCB1_-*Δ*CBS* (in which the CBS sequence was deleted from the *LCB1* promoter) did not exhibit Myr sensitivity comparable to that of *com2*Δ (Figure 5—figure supplement A-C).

Taken together, these results suggest that Ypk1 represent a major Com2 target under sphingolipid-depleted conditions, although additional Com2-dependent targets likely contribute to sphingolipid homeostasis.

### Mapping *COM2* regulatory regions required for sphingolipid-dependent Com2 expression

To identify the regions required for Com2 expression control, we constructed a series of plasmids carrying stepwise deletions in either the ∼500 bp 5′ upstream region (putative *COM2* promoter) or the *COM2* ORF (Figure 6A). These plasmids were introduced into *com2*Δ cells, and we compared Myr sensitivity and Com2 protein abundance in the presence or absence of Myr (Figure 6). Whereas *com2*Δ cells were Myr-sensitive, introduction of most plasmids restored Myr resistance to varying degrees (Figure 6B). In contrast, the Δ2-190 mutant and truncation mutants lacking the DNA region corresponding to the Zn-finger domain (1-379, 1-230, 1-143, 1-87) failed to complement Myr sensitivity.

**Figure 6.**
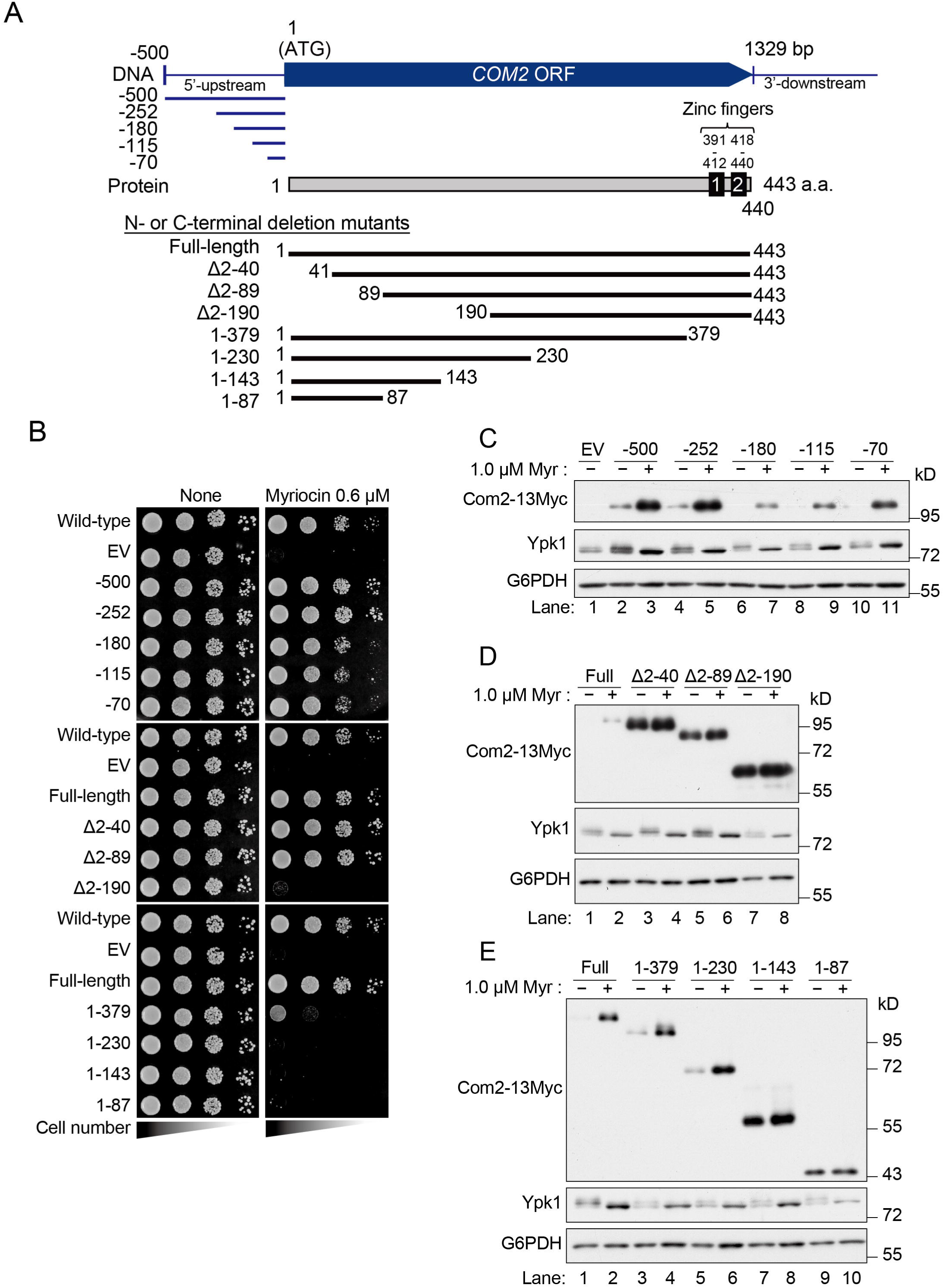
Com2 expression is regulated by sequences within the ORF region. (**A**) Schematic representation of Com2 expression plasmids carrying stepwise deletions in the 5′ upstream promoter region or truncations of the Com2 open reading frame (ORF) from either the N-terminal or C-terminal side. (**B**) The promoter- or ORF-deletion mutants of Com2 shown in (A) were transformed into *com2*Δ cells, and functional complementation of the Myr sensitivity of the *com2*Δ strain was assessed by spot assays using 10-fold serial dilutions on YPD medium or YPD medium supplemented with 0.6 µM Myr. (**C**) Expression of Com2-13Myc variants carrying stepwise deletions in the 5′ upstream promoter region of *COM2* was examined in untreated cells or cells treated with 1.0 µM Myr for 3 hrs. (**D**) Expression of Com2-13Myc variants carrying N-terminal truncations of the ORF was analyzed under the same conditions as in (C). (**E**) Expression of Com2-13Myc variants carrying C-terminal truncations of the ORF was analyzed under the same conditions as in (C). For (C-E), *com2*Δ cells expressing 13Myc-tagged Com2 wild-type or deletion variants were grown to mid-log phase in SD (-Ura) liquid medium at 26°C, treated with 1.0 µM Myr for 3 hrs, and lysed. Proteins were separated by SDS-PAGE and immunoblotted with anti-Myc, anti-Ypk1, or anti-G6PDH antibodies to detect Com2-13Myc, Ypk1, or G6PDH (loading control), respectively.

We next examined the expression of these mutants by immunoblotting. For the promoter-deletion series, removal of ≥180 bp from the 5′ upstream region caused a marked reduction in basal Com2 abundance. Nevertheless, all promoter-deletion constructs still showed a clear increase in Com2 levels upon Myr treatment (Figure 6C). Consistently, in a strain in which the endogenous *COM2* promoter was replaced with a tet-off promoter (*P*_tet-off_*-13Myc-COM2*), Com2 levels increased over time after Myr addition both under “induced (-Dox)” and “repressed (+Dox)” conditions (Figure 6—supplement 1). These results suggest that Myr-dependent upregulation of Com2 is not primarily driven by the native promoter.

In contrast, stepwise deletions within the *COM2* ORF revealed a prominent effect on Com2 abundance. Deletion of ≥40 amino acids from the N-terminus caused a strong accumulation of Com2 protein irrespective of Myr treatment (Figure 6D). A C-terminal truncation removing 213 residues (1-230) still exhibited Myr-dependent induction, whereas more extensive C-terminal truncations (1-143 and 1-87) accumulated to high levels regardless of Myr (Figure 6E). Together, these findings indicate that Com2 expression control is largely determined by sequences within the ORF, rather than by the promoter.

### Com2 is rapidly degraded by the ubiquitin-proteasome system in response to increased sphingolipid levels

Because Com2 abundance increases upon Myr-induced sphingolipid depletion and this response is independent of the promoter, we next asked whether increasing intracellular sphingolipids would conversely downregulate Com2. Three hours after Myr treatment, we added phytosphingosine (PHS), a ceramide precursor, to bypass the blocked step and restore sphingolipid synthesis (Figure 7A). Com2 levels, which were elevated by Myr, dropped rapidly within 1 hr of PHS addition (Figure 7B, C). In contrast, Ypk1 levels also increased upon Myr treatment but did not show a significant decrease after PHS addition (Figure 7B, D). Notably, TORC2-dependent phosphorylation of Ypk1 (p-Ypk1^T662^) increased over time during Myr treatment and declined promptly after PHS addition (Figure 7B, E). These results suggest that Com2 abundance is dynamically tuned to intracellular sphingolipid levels.

**Figure 7.**
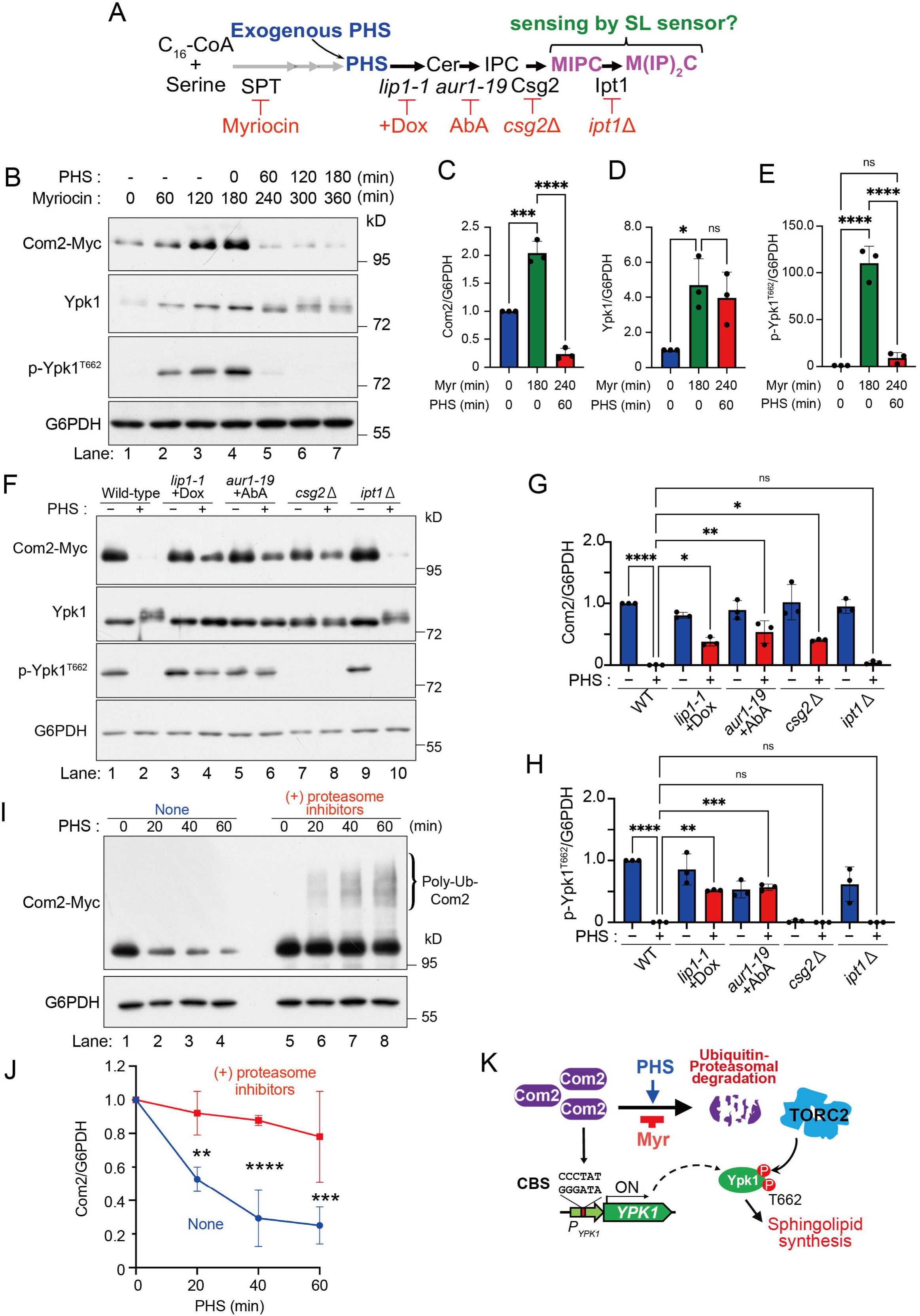
Com2 is regulated by ubiquitin-proteasome system-mediated degradation in a manner dependent on intracellular sphingolipid levels. (**A**) Sphingolipid biosynthetic pathway and perturbations at each step (Myr, PHS bypass, *lip1-1* +Dox, *aur1-19* +AbA, *csg2*Δ, *ipt1*Δ). (**B**) Immunoblot of Com2 and Ypk1 in wild-type cells treated with 1.0 µM Myr for 3 hrs, followed by 10 µM PHS for 3 hrs. G6PDH, loading control. (**C-E**) Quantification of (B): Com2 (C), Ypk1 (D), and p-Ypk1^T662^ (E). One-way ANOVA with Tukey’s test. (**F**) PHS-induced Com2 degradation in wild-type and sphingolipid-pathway-perturbed cells. Log-phase cultures were pretreated with 1 µM Myr for 3 hrs, then PHS for 3 hrs. *lip1-1* was cultured with Dox (20 µg/mL) from the preculture; *aur1-19* was pretreated with AbA (20 µg/mL) for 15 min before PHS. Immunoblots for Com2, Ypk1, and p-Ypk1^T662^; G6PDH, loading control. (**G**, **H**) Quantification of (F). One-way ANOVA with Tukey’s test. (**I**) Proteasome inhibition stabilizes ubiquitinated Com2. Wild-type cells depleted of sphingolipids by Myr (3 hrs) were incubated ±MG132 (50 µM) and bortezomib (100 µM) for 30 min, followed by PHS (10 µM). Samples were collected at 0-60 min and immunoblotted with anti-Myc (Com2-13Myc) and anti-G6PDH. (**J**) Quantification of (I) (normalized to G6PDH). Two-way ANOVA with Sidak’s test. (**K**) Model: Myr stabilizes Com2 to promote *YPK1* expression and sphingolipid biosynthesis, whereas under +PHS conditions, Com2 is rapidly degraded via the ubiquitin-proteasome system. Quantified data (C-E, G-H, J) are mean ± SD (n=3). \**p*<0.1, \*\**p*<0.01, \*\*\**p*<0.001, \*\*\*\**p*<0.0001; n.s., not significant.

To pinpoint the biosynthetic step responsible for the PHS-dependent reduction in Com2, we used loss-of-function mutants and step-specific inhibitors targeting distinct stages of sphingolipid synthesis (Figure 7A). Blocking mannnosyldiinositol phosphorylceramide, M(IP) _2_C synthesis (*ipt1*Δ) did not impair the PHS-dependent decrease in Com2, which remained comparable to that in wild-type cells (Figure 7F, G). By contrast, inhibiting ceramide synthesis (*lip1-1* +Dox), IPC synthesis (*aur1-19* +AbA), or mannnosylinositol phosphrylceramide, MIPC synthesis (*csg2*Δ) abolished the PHS-dependent reduction in Com2 (Figure 7F, G). Thus, Com2 downregulation requires flux through the pathway up to MIPC. The rapid PHS-induced change in Com2 expression correlated with changes in p-Ypk1^T662^ levels in wild-type cells and in mutants impaired at different steps of sphingolipid metabolism, except for those in *csg2*Δ cells (Figure 7F, H).

We next considered that the rapid loss of Com2 following PHS addition might reflect regulated protein degradation, such as proteasomal degradation or autophagy. Co-treatment with the proteasome inhibitors MG132 and bortezomib markedly suppressed the PHS-dependent decrease in Com2 and led to the appearance of high-molecular-weight species consistent with polyubiquitinated Com2 (Figure 7I, J). In contrast, deletion of *ATG5*, which is essential for autophagosome formation, did not prevent the PHS-dependent decrease in Com2 (Figure 7—figure supplement). Together, these data indicate that Com2 is downregulated primarily through ubiquitin-proteasome system (UPS)-dependent degradation.

In summary, our results support a model in which an increase in the complex sphingolipids triggers UPS-dependent degradation of Com2, thereby providing negative feedback to restrain downstream sphingolipid synthesis (Figure 7K).

### Identification of Com2 ubiquitination sites and the role of phosphorylation in sphingolipid-dependent Com2 degradation

Because deletion of amino acids 2-40 markedly increased Com2 expression levels (Figure 6D), we hypothesized that this region is involved in PHS-dependent degradation. Consistent with this idea, PHS-dependent degradation of Com2 was completely abolished in the mutant lacking amino acids 2-40 (Figure 8A).

**Figure 8.**
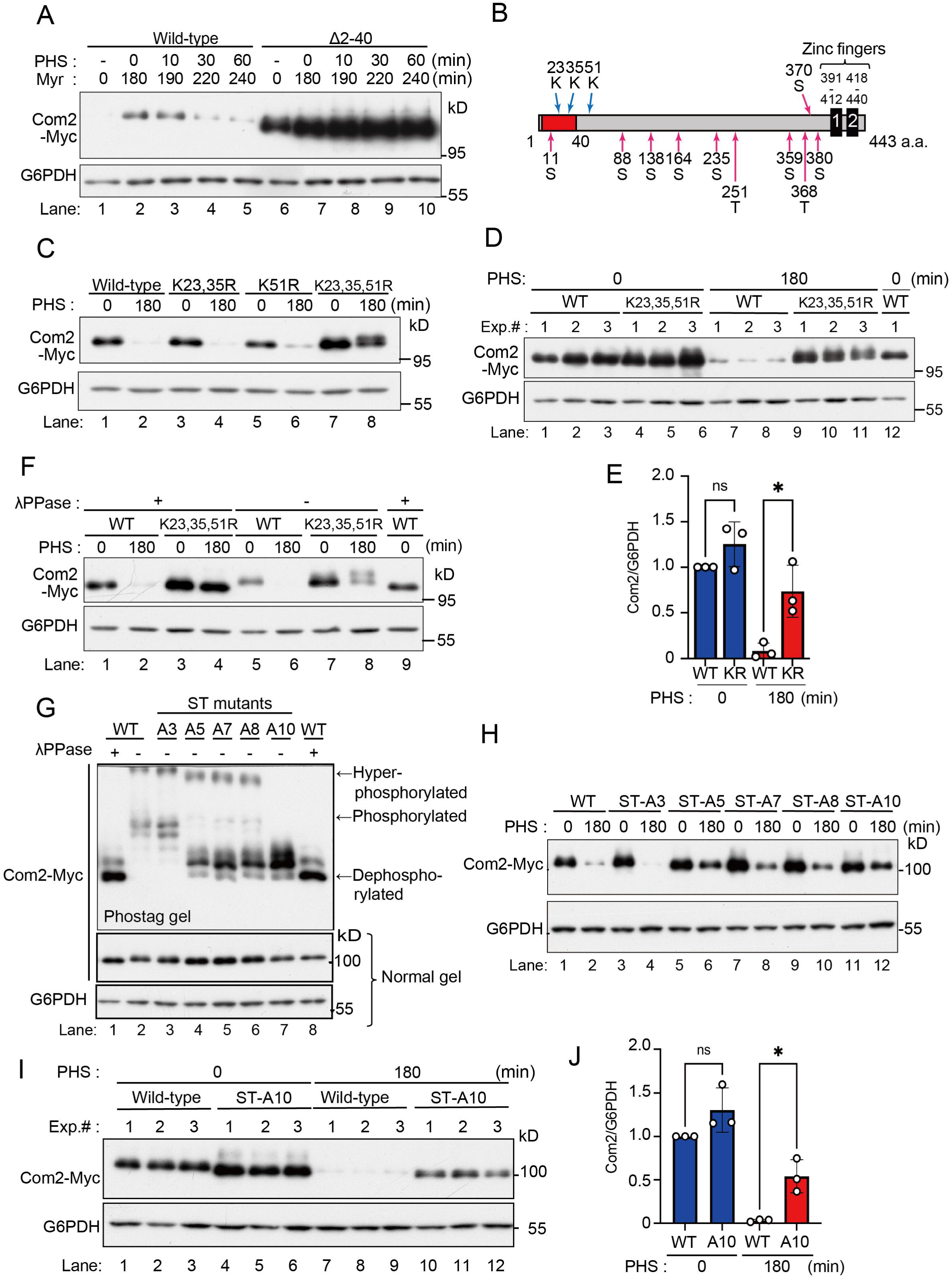
Ubiquitination sites required for sphingolipid-dependent proteasomal degradation of Com2 and the role of Com2 phosphorylation. (**A**) *com2*Δ cells expressing wild-type Com2 or Com2 (Δ2-40) were treated with 1.0 µM Myr for 3 hrs, then with PHS for 3 hrs; Com2 levels were analyzed by immunoblotting (G6PDH, loading control). (**B**) Schematic of predicted ubiquitination and phosphorylation sites in Com2. (**C**) *com2*Δ cells expressing wild-type Com2 or the indicated lysine mutants (K23,35R, K51R, K23,35,51R) were treated with 1.0 µM Myr for 3 hrs, then 10 µM PHS for 3 hrs; Com2 levels after Myr and after PHS were analyzed by immunoblotting (G6PDH, loading control). (**D**, **E**) Immunoblot (D) and quantification (E) comparing wild-type and K23,35,51R Com2 before and after PHS (n=3; one-way ANOVA with Tukey’s test; \**p*<0.1; n.s.). (**F**) Com2 phosphorylation was assessed ±λPPase in cells expressing wild-type Com2 or K23,35,51R after Myr→PHS treatment; immunoblotting for Com2 (G6PDH, loading control). (**G**) Phos-tag SDS-PAGE of Com2 phosphorylation-site mutants (ST-A3, ST-A5, ST-A7, ST-A8, ST-A10) based on mobility shifts. (**H**) Immunoblot analysis of Com2 levels in the indicated phosphorylation-site mutants before and after Myr (3 hrs) → PHS (3 hrs) treatment (G6PDH, loading control). (**I**, **J**) Immunoblot (I) and quantification (J) comparing wild-type and ST-A10 Com2 before and after PHS (n=3; one-way ANOVA with Tukey’s test; \**p*<0.1; n.s.).

To identify ubiquitination sites, lysine residues located within or near this region were substituted with arginine (Figure 8B). Mutation of K23 and K35, either individually or together, or mutation of K51 alone, did not affect PHS-dependent degradation. In contrast, simultaneous mutation of K23, K35, and K51 (KR mutant) significantly suppressed PHS-induced degradation (Figure 8C-E). The KR mutant fully complemented the Myr sensitivity of the *com2*Δ strain, indicating that these substitutions do not impair Com2 function (Figure 8—figure supplement 1A). Notably, the KR mutant exhibited a PHS-induced mobility shift that was reversed by λphosphatase (λPPase) treatment, indicating phosphorylation-dependent modification (Figure 8F). These results suggest that ubiquitination at K23, K35, and K51 promotes proteasome-dependent degradation of Com2.

Because degradation was suppressed in the KR mutant despite increased phosphorylation, we next examined the role of phosphorylation in Com2 stability. Phos-tag SDS-PAGE resolved several phosphorylated Com2 species that shifted downward following λPPase treatment, demonstrating that Com2 is highly phosphorylated (Figure 8G, lanes 1, 2).

In addition to the five phosphorylation sites annotated in the Saccharomyces Genome Database, we identified five predicted AGC kinase target sites, including putative Ypk1 phosphorylation sites, yielding a total of ten candidate phosphorylation sites (Fig. 8—figure supplement 1B). Progressive substitution of these serine/threonine residues with alanine (ST-A3 to ST-A10) caused stepwise reduction of phosphorylation-dependent band shifts, with ST-A10 showing a marked loss of phosphorylation (Figure 8G, lanes 2-7).

Consistent with this result, PHS-dependent degradation was unchanged in ST-A3 but progressively suppressed in mutants containing five or more substitutions and was significantly reduced in ST-A10 (Figure 8H-J). Together, these results indicate that phosphorylation of Com2 is required for its PHS-dependent degradation.

## Discussion

Bilayer assembly during membrane biogenesis requires the coordinated supply and organization of multiple lipid species, necessitating integrated control of lipid synthesis, uptake, metabolism, and subcellular distribution (Nohturfft and Zhang, 2009). Global regulation of lipid metabolism is achieved not only through signaling pathways that modulate the activity of lipid metabolic enzymes, but also through master transcriptional regulators that coordinate expression of genes encoding these enzymes and their modulators. Master transcriptional regulators have been identified for glycerophospholipids, lipid-saturation control, and sterol metabolism in both yeast and mammals (Covino et al., 2018; Nohturfft and Zhang, 2009). In contrast, although sphingolipid metabolic flux is well understood to be regulated through the TORC2-Ypk1 pathway via control of biosynthetic enzyme activity (Berchtold et al., 2012), transcriptional mechanisms governing sphingolipid metabolism remain far less defined (Breslow and Weissman, 2010).

In this study, we show that Com2 functions as a master transcriptional regulator of sphingolipid metabolism in budding yeast by responding to reduced sphingolipid levels, upregulating its own expression, and promoting Ypk1 expression to stimulate sphingolipid synthesis.

Previous work reported that *com2*Δ cells are hypersensitive to sulfur dioxide (SO_2_) and that Com2 acts as a major transcription factor controlling expression of genes required for SO_2_ tolerance (Lage et al., 2019). Com2 has also been identified as a regulator of the UASru element within the promoter of *IME1*, the master regulator of meiosis in budding yeast, where deletion of this element markedly reduces *IME1* transcription (Kahana-Edwin et al., 2013). How the Com2-dependent program uncovered here intersects with these pathways remains unclear; however, Com2 may employ shared regulatory logic across distinct physiological responses, an idea that will be important to address in future studies.

### *YPK1* is a major Com2 target, but additional targets likely contribute to sphingolipid homeostasis

Our data show that *YPK1* is one of the major transcriptional targets of Com2 in sphingolipid metabolism. A genome-wide search based on the Com2-binding site (CBS) consensus sequence also identified a CBS-like element in the promoter of *LCB1*, which encodes a catalytic subunit of SPT, and a *LacZ* reporter assay further suggested that the *LCB1* promoter can respond to Com2. Notably, deletion of the CBS from the *YPK1* promoter (*P_YPK1_*-ΔCBS) reduced Myr-induced Ypk1 upregulation to a level comparable to that seen in *com2*Δ cells (Figure 5D). However, this promoter mutant did not phenocopy the strong Myr sensitivity of *com2*Δ. Likewise, a double mutant lacking CBS elements in both the *YPK1* and *LCB1* promoters failed to recapitulate the *com2*Δ phenotype (Figure 5—figure supplement). Together, these results suggest that, in addition to *YPK1* and *LCB1*, Com2 likely controls other target genes that collectively contribute to sphingolipid homeostasis.

### Com2 degradation is triggered by a cellular readout of complex sphingolipids

Com2 levels increased when sphingolipid synthesis was reduced by Myr treatment, whereas exogenous addition of phytosphingosine (PHS), a ceramide precursor, rapidly promoted Com2 degradation via the ubiquitin-proteasome system (UPS) (Figure 7B, I). Systematic inhibition of individual steps in the sphingolipid pathway further suggested that the PHS-dependent degradation signal reflects the status of complex sphingolipids at or downstream of MIPC (Figure 7F). In glycerophospholipid and sterol metabolism, several lipid-sensing transcriptional systems are well established—for example, Opi1 sensing phosphatidic acid (Loewen et al., 2004), Mga2/Spt23 sensing membrane lipid saturation to control *OLE1* expression (Ballweg et al., 2020; Romanauska and Köhler, 2021), and Upc2 sensing sterol depletion to induce sterol biosynthetic genes (Yang et al., 2015). In contrast, the molecular mechanisms by which cells sense sphingolipid levels, particularly complex sphingolipids, remain poorly understood (Breslow and Weissman, 2010). TORC2-Ypk1 signaling is also thought to be influenced by complex sphingolipid status (Figure 7F, H), yet the identity of the upstream sensor is still unclear. Thus, dissecting the mechanism of Com2 degradation may provide a tractable entry point to uncover the elusive sphingolipid sensor operating at the top of this pathway.

### How are Com2 ubiquitination and degradation controlled?

Many transcription factors are regulated through proteolysis by the UPS in response to environmental and metabolic cues, including oxidative stress, heat shock, nutrient limitation, changes in cellular redox status, and shifts in metabolic state (e.g., glucose availability, AMP/ATP, NAD^+^/NADH, and TCA-cycle metabolites) (Geng et al., 2012). Within the UPS, E3 ubiquitin ligases act as central decision-makers that determine whether a substrate is degraded. In many cases, this decision is implemented through the recognition of degron motifs and their modulation by post-translational modifications, thereby converting stress/metabolic signals into changes in protein half-life (Finley et al., 2012).

Here, we found that PHS-induced degradation of Com2 critically depends on its N-terminal region (notably the segment around residues 2-40) (Figure 8A). Moreover, substitution of lysine residues near the N terminus with arginine (K23R, K35R, and K51R; hereafter the KR mutant) markedly attenuated PHS-dependent Com2 degradation. These observations suggest that the N-terminal region may provide ubiquitin-acceptor lysines and/or serve as a degron that is recognized by an E3 ubiquitin ligase.

Notably, the KR mutant displayed an increased phosphorylation signal after PHS treatment (Figure 8F). In budding yeast, the transcription factor Gcn4 is phosphorylated by Pho85 under non-starvation conditions, which generates a phosphodegron that is recognized by SCF^Cdc4^, leading to ubiquitination and proteasomal degradation (Meimoun et al., 2000). By analogy, Com2 phosphorylation may similarly act as a trigger for UPS-dependent degradation, potentially by creating or exposing a phosphodegron. Consistent with this idea, a mutant in which multiple predicted phosphorylation sites were collectively replaced (ST-A10) exhibited reduced phosphorylation and a significant suppression of PHS-dependent degradation (Figure 8G-J). Together, these results support a model in which Com2 phosphorylation is an important upstream step in its degradation. Nevertheless, the increased phosphorylation observed in the KR mutant could also reflect accumulation of phosphorylated Com2 due to impaired turnover, and future work will be required to establish the causal order of phosphorylation and ubiquitination.

Based on these findings, we propose the model shown in Figure 9. When intracellular sphingolipid levels are low, Com2 is stabilized and promotes sphingolipid synthesis by upregulating genes that positively regulate the pathway and contain Com2-binding sites (CBS), such as *YPK1* and *LCB1*. Conversely, when sphingolipid levels are high, an as-yet-unidentified complex sphingolipid sensor and downstream signaling pathway may promote Com2 phosphorylation, recruit an E3 ubiquitin ligase, and drive Com2 ubiquitination followed by rapid proteasomal degradation.

**Figure 9.**
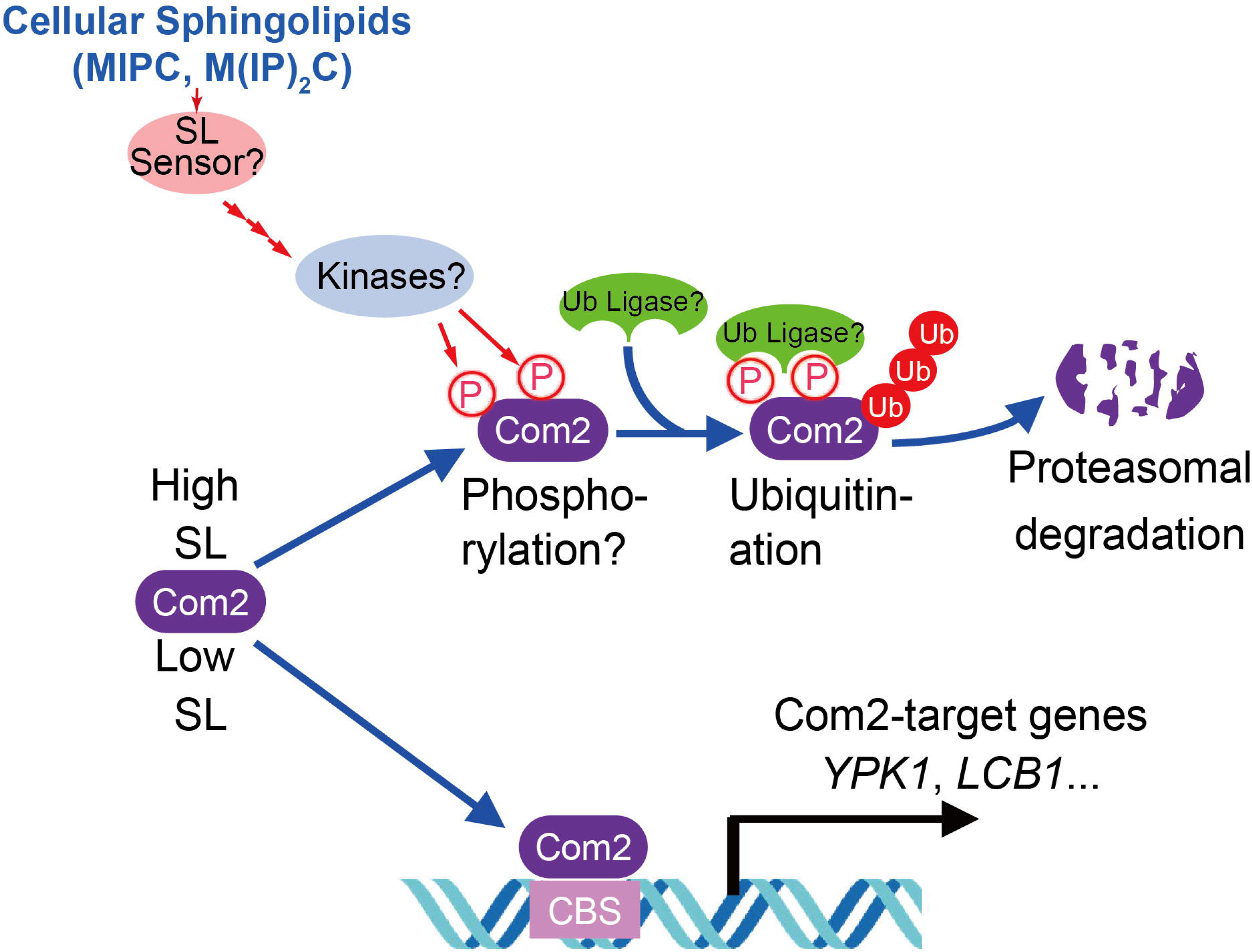
Model for sphingolipid (SL) level-dependent regulation of Com2 degradation via the ubiquitin-proteasome system. When intracellular SL levels are low, degradation of Com2 is suppressed, allowing Com2 to induce the expression of its target genes, such as *YPK1* and *LCB1*, which harbor Com2-binding sites (CBSs) in their promoter regions, thereby promoting sphingolipid biosynthesis. In contrast, when intracellular SL levels are sufficient, complex sphingolipids such as MIPC, M(IP)_2_C are sensed by an as-yet-unidentified sphingolipid sensor, leading to phosphorylation of Com2, presumably by an unidentified kinase. This phosphorylation promotes ubiquitination of Com2 and its subsequent degradation by the proteasome.

### Is transcriptional control of sphingolipid metabolism evolutionarily conserved?

Whether a Com2-like mechanism of sphingolipid metabolic control is evolutionarily conserved in mammals remains an open question. In mice, ZFP750 (ZNF750 in humans) has been reported to function as a master regulator of ceramide synthase genes, particularly in the skin, where it is essential for establishing epidermal barrier function by maintaining proper lipid composition (Butera et al., 2023). ZFP750 directly binds to the promoters of lipid metabolic enzyme genes, including Elovl6 and Elovl7, which are involved in fatty acid elongation, thereby regulating lipid metabolism at the transcriptional level.

Although Com2 cannot be considered a clear ortholog of ZFP750, it is tempting to speculate that ZFP750 may also be subject to functional regulation linked to sphingolipid levels, analogous to that observed for Com2. Notably, more than 700 C2H2-type zinc finger proteins are annotated in the human genome (Nabeel-Shah et al., 2024), raising the possibility that additional transcription factors may regulate sphingolipid metabolism in a manner similar to Com2.

Our findings therefore provide a conceptual framework for understanding sphingolipid homeostasis as a transcriptionally controlled process that may be conserved beyond yeast.

## Materials and Methods

### Strains, plasmids, and media

Descriptions of the strains and plasmids used in this study are presented in Tables 2 and Table 3. *Escherichia coli* DH5α or JM109 was used as the bacterial host for plasmid construction. Coding sequences were amplified by means of PCR, using KOD-Plus-Neo polymerase (Toyobo) or PrimeSTAR^®^ GXL polymerase (TaKaRa Bio Inc.). The plasmids were sequenced to ensure that no mutations were introduced owing to the manipulation. Mutant constructs were generated using site-directed mutagenesis and confirmed using sequencing. The media used for yeast culture were YPD (1% yeast extract, 2% peptone, and 2% glucose) or synthetic dextrose (SD) medium (2% glucose and 0.67% yeast nitrogen base without amino acids). Appropriate amino acids and bases were added to the SD medium, as necessary. Yeast cells were cultured at 26°C, unless otherwise stated. All deletion and tagged strains were constructed using homologous recombination with PCR-generated DNA fragments, as described previously (Longtine et al., 1998). Multiple-mutant strains of *S. cerevisiae* were generated by genetic crossing of the respective single-mutant strains followed by tetrad dissection analysis.

**Table 2.**
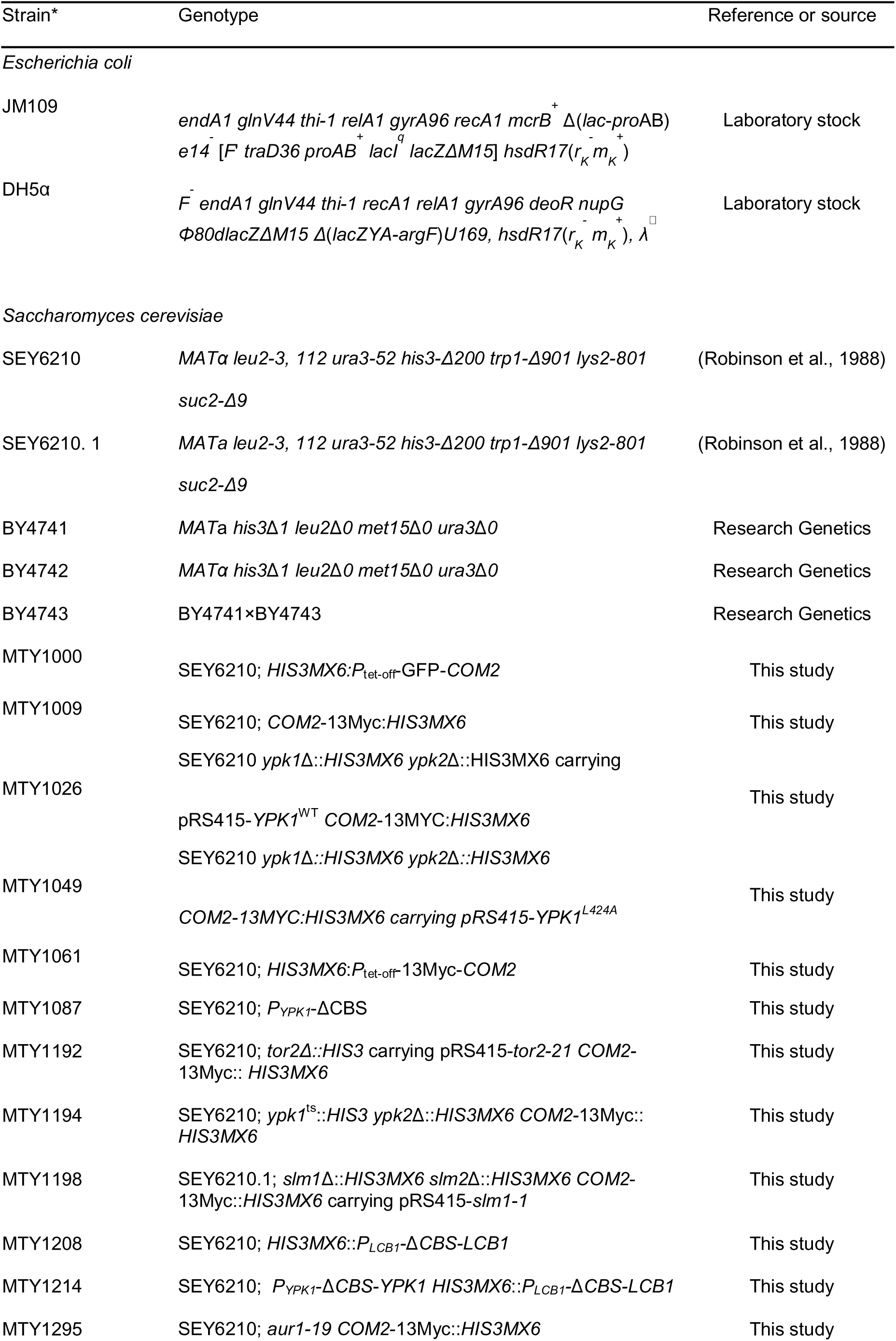
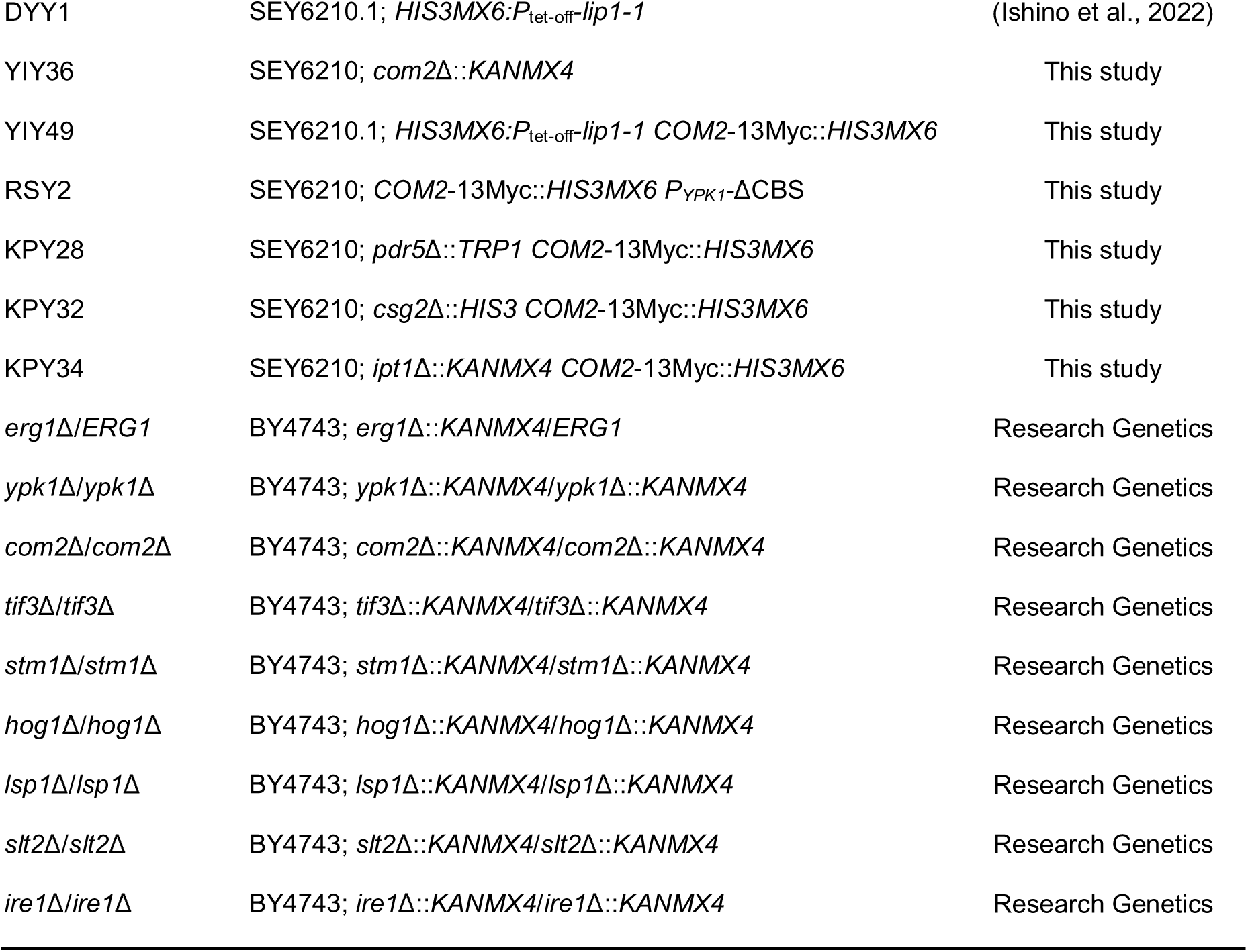
Strains used in this study.

**Table 3.** Plasmids used in this study.

| Plasmids | Construct | Reference or source |
| --- | --- | --- |
| pRS415 | *Amp, LEU2, CEN6* | (Sikorski and Hieter, 1989) |
| pRS416 | *Amp, URA3, CEN6* | (Sikorski and Hieter, 1989) |
| pRS426 | *Amp, URA3, 2 μm* | (Sikorski and Hieter, 1989) |
| pRT3 | pRS416; *URA3, CEN, HIS3MX6-P<sub>ADH1</sub>-tTA2s-P<sub>tetoff</sub>-GFP* | (Ishino et al., 2022) |
| pMT1447 | pRS416; *URA3, CEN, P<sub>SNR52</sub>-YPK1-2-gRNA-T<sub>SUP</sub>-Z4EV-P<sub>Z4EV</sub>-CAS9-CYC1pA* | This study |
| pMT1536 | pRS416; *URA3, CEN, P<sub>COM2</sub>-COM2<sup>T251A+S370, 380A</sup>-13Myc (ST3A)* | This study |
| pMT1547 | pRS416; *URA3, CEN, P<sub>COM2</sub>-COM2<sup>S88, 164, 370, 380A+T251A</sup>-13Myc (ST4A)* | This study |
| pMT1576 | pRS416; *URA3, CEN, P<sub>COM2</sub>-COM2<sup>S88, 164, 235, 370, 380A+T251A</sup>-13Myc (ST6A)* | This study |
| pMT1577 | pRS416; *URA3, CEN, P<sub>COM2</sub>-COM2<sup>S88, 164, 235, 359, 370, 380A+T251A</sup>-13Myc (ST7A)* | This study |
| pMT1578 | pRS416; *URA3, CEN, P<sub>COM2</sub>-COM2<sup>S88, 164, 235, 359, 370, 380A+T251, 368A</sup>-13Myc (ST8A)* | This study |
| pMT1790 | pRS416; *URA3, CEN, P<sub>COM2</sub>-COM2<sup>S11, 88, 138, 164, 235, 359, 370, 380A+T251, 368A</sup>-13Myc (ST10A)* | This study |
| pYI41 | yEP352 Suppressor #6 (*YPK1*) | This study |
| pYI42 | yEP352 Suppressor #13 (*TIF3*) | This study |
| pYI44 | yEP352 Suppressor #26 (*RIM20* (partial), *CAF20, HEM4, RFM1*) | This study |
| pYI46 | yEP352 Suppressor #34 (*COM2*) | This study |
| pYI47 | yEP352 Suppressor #36 (*SPO77* (partial), *PKC1* (partial)) | This study |
| pYI48 | yEP352 Suppressor #21 (*COM2*) | This study |
| pYI51 | yEP352 Suppressor #9 (*LSP1*) | This study |
| pYI53 | yEP352 Suppressor #176 (*BMH2*) | This study |
| pYI56 | yEP352 Suppressor #56 (*PIN4*) | This study |
| pYI58 | yEP352 Suppressor #66 (*SUT2*) | This study |
| pYI62 | pRS416; *P<sub>COM2</sub>-COM2-13Myc (COM2 WT)* | This study |
| pYI68 | p*P<sub>YPK1</sub>-LacZ* | This study |
| pAMS366 | *Amp, URA3, 4×CDRE-LacZ* | (Stathopoulos and Cyert, 1997) |
| pRS16 | p$P_{YPK1-\Delta CBS}$-LacZ | This study |
| pRS18 | p$P_{LCB1}$-LacZ | This study |
| pRS28 | p$P_{LCB1-\Delta CBS}$-LacZ | This study |
| pNK18 | pRS416; *URA3*, *CEN*, $P_{YPK1}$-3HA-YPK1::HIS3MX6 | This study |
| pNK19 | pRS416; *URA3*, *CEN*, $P_{YPK1}$-ΔCom2 binding site (ΔCBS)-3HA-YPK1::HIS3MX6 | This study |
| pTU3 | pRS416; $P_{COM2}$ (-500 to -252)-COM2-13Myc | This study |
| pTU4 | pRS416; $P_{COM2}$ (-500 to -115)-COM2-13Myc | This study |
| pTU6 | pRS416; $P_{COM2}$ (-500 to -70)-COM2-13Myc | This study |
| pTU14 | pRS416; $P_{COM2}$-COM2 (Δ2-89)-13Myc | This study |
| pTU15 | pRS416; $P_{COM2}$-COM2 (Δ2-190)-13Myc | This study |
| pTU17 | pRS416; $P_{COM2}$ (-500 to -180)-COM2-13Myc | This study |
| pTU28 | pRS416; $P_{COM2}$-COM2 (Δ380-443)-13Myc | This study |
| pTU29 | pRS416; $P_{COM2}$-COM2 (Δ231-443)-13Myc | This study |
| pTU30 | pRS416; $P_{COM2}$-COM2 (Δ88-443)-13Myc | This study |
| pTU33 | pRS416; $P_{COM2}$-COM2 (Δ2-40)-13Myc | This study |
| pTU34 | pRS416; $P_{COM2}$-COM2 (Δ144-443)-13Myc | This study |
| pKP6 | pRS416; $P_{COM2}$-COM2 (Δ2-10)-13Myc | This study |
| pKP7 | pRS416; $P_{COM2}$-COM2 (Δ2-20)-13Myc | This study |
| pKP8 | pRS416; $P_{COM2}$-COM2 (Δ2-30)-13Myc | This study |
| pKP13 | pRS416; $P_{COM2}$-COM2 (Δ2-114)-13Myc | This study |
| pKP14 | pRS416; $P_{COM2}$-COM2 (Δ2-139)-13Myc | This study |
| pKP15 | pRS416; $P_{COM2}$-COM2 (Δ2-164)-13Myc | This study |
| pKP20 | pRS416; $P_{COM2}$-COM2 (Δ240-443)-13Myc | This study |
| pKP21 | pRS416; $P_{COM2}$-COM2 (Δ290-443)-13Myc | This study |
| pKP22 | pRS416; $P_{COM2}$-COM2 (Δ340-443)-13Myc | This study |
| pKP23 | pRS416; $P_{COM2}$-COM2 (Δ390-443)-13Myc | This study |
| pAN28 | pRS416; $P_{COM2}$-COM2<sup>K23,35R</sup>-13Myc | This study |
| pAN30 | pRS416; $P_{COM2}$-COM2<sup>K23,35,51R</sup>-13Myc | This study |
| pAN36 | pRS416; $P_{COM2}$-COM2<sup>K51R</sup>-13Myc | This study |

### Yeast growth assays

For the yeast plate growth assay, cells were adjusted to an OD_600_ of 1.0, serially diluted in fourfold steps, and 5 μL of each dilution was spotted onto assay plates. Plates were incubated at the indicated temperature for 3-4 days and then photographed.

For the liquid growth assay, as described previously (Ishino et al., 2022), cells were inoculated into YPD medium or YPD supplemented with Myr at an initial OD_600_ of 0.1. Growth was monitored every 30 min at 26°C using a microplate reader (Tecan Sunrise, Tecan Japan, Co., Ltd, Kanagawa, Japan) with three biological replicates.

### Antibody production

The rabbit polyclonal antibody that recognizes Com2 was produced as follows. A His_6_-fused Com2 protein was expressed in *E. coli* Rosetta-gami B (DE3) pLysS cells and the total cell lysate was separated on an SDS-PAGE gel. The band corresponding to the His_6_-Com2 fusion protein was excised from the SDS-PAGE gel and electroeluted from the gel slices using an AE-3590 electrochamber (Atto). An aliquot of approximately 300 μg of the fusion protein was emulsified with Freund’s complete adjuvant and injected intramuscularly or subcutaneously into young female Japanese white rabbits. Antisera were purified using an affinity column immobilized with His_6_-ProS2-Com2 (382-443 a.a.) fusion protein.

### Western blot

Protein extracts were prepared using the TCA precipitation method, loaded onto normal or Phos-tag SDS-PAGE gels, and transferred to PVDF membranes (Fujifilm Wako). λPPase treatment was performed as described previously (Ishino et al., 2022). The antibodies and dilutions used in this study were listed in Table 4.

**Table 4.** Key resources table.

| Reagent type (species) or resource | Designations | Source or reference | Identifiers | Additional information |
| --- | --- | --- | --- | --- |
| Antibody | Anti-Com2 (Rabbit polyclonal) | This study | Home made | WB (1:2,000) |
| Antibody | Anti-Myc (Mouse monoclonal) | Medical & Biological Laboratories Co., Ltd., Nagoya, Japan | Cat # M192-3 | WB (1:5,000) |
| Antibody | Anti-FLAG (Mouse monoclonal) | Sigma-Aldrich, Merck KGaA, Darmstadt, Germany | Cat # F1804 | WB (1:2,000) |
| Antibody | Anti-Ypk1 (Rabbit polyclonal) | (Ishino et al., 2022) | Home made | WB (1:2,000) |
| Antibody | Anti-phospho-Ypk2<sup>T659</sup> (Rabbit polyclonal) for detection of both p-Ypk1<sup>T662</sup> and p-Ypk2 | (Ishino et al., 2022) | MBL, Nagoya, Japan; custom-made | WB (1:50,000) |
| Antibody | Anti-G6PDH (Rabbit polyclonal) | Sigma-Aldrich, Merck KGaA, Darmstadt, Germany | Cat # SAB2100871 | WB (1:50,000) |
| Antibody | Anti-mouse IgG (HRP-conjugated) | Cell Signaling Technology, Danvers, MA, USA | Cat # 7076 | WB (1:10,000) |
| Antibody | Anti-rabbit IgG (HRP-conjugated) | Capel, MP Biomedicals, Solon, OH, USA | Cat # 55611 | WB (1:10,000) |
| Enzyme | KOD -Plus- Neo polymerase | TOYOBO, Co., Ltd., Osaka, Japan | Cat # KOD-401 |  |
| Enzyme | PrimeSTAR<sup>®</sup> GXL polymerase | TaKaRa Bio, Shiga, Japan | Cat # R050A |  |
| Enzyme | Taq DNA polymerase | BioAcademia, Osaka, Japan | Cat # 02-012 |  |
| Chemical compound | Myriocin | Sigma-Aldrich, Merck KGaA, Darmstadt, Germany | Cat # M1177 |  |
| Chemical compound | Aureobasidin A | TaKaRa Bio, Shiga, Japan | Cat # 630499 |  |
| Chemical compound | Doxycycline Hyclate | Tokyo Chemical Industry Co., Ltd., Tokyo, Japan | Cat # D4116 |  |
| Chemical compound | Miconazol | FUJIFILM Wako Pure Chemical Corporation, Osaka, Japan | Cat # 510-22771 |  |
| Chemical compound | Telbinafine | FUJIFILM Wako Pure Chemical Corporation, Osaka, Japan | Cat # 516-40951 |  |
| Chemical compound | Tunicamycin | FUJIFILM Wako Pure Chemical Corporation, Osaka, Japan | Cat # 202-08241 |  |
| Chemical compound | Fluorescent Brightener 28 | FUJIFILM Wako Pure Chemical | Cat # 593-28141 |  |
|  | (Calcofluor White) | Corporation, Osaka, Japan |  |  |
| Chemical compound | 1-(tert-butyl)-3-(3-methylbenzyl)-1H-pyrazole[3,4-d]pyrimidin-4-amine, 3-MB-PP1 | Cyman Chemical, Ann Arbor, MI, USA | Cat # 17860 |  |
| Software | GraphPad Prism 10 | Graphpad Software, San Diego, CA, USA |  | 10.5.0 |
| Software | Adobe Photoshop 2024 | Adobe Inc., San Jose, CA, USA |  | 25.3.1 |
| Software | Adobe Illustrator 2024 | Adobe Inc., San Jose, CA, USA |  | 28.7.10 |
| Software | Microsoft Excel 365 | Microsoft corporation, Redmond, WA, USA |  |  |

### Sphingolipid analysis

Sphingolipids were extracted from *S. cerevisiae* and analyzed using TLC, as described previously (Yamaguchi et al., 2018).

### Construction of *LacZ* reporter plasmids

The fragments containing the promoter regions of *YPK1* and *LCB1* were amplified by means of PCR and cloned into the XhoI and BamHI sites of the 4×*CDRE-LacZ* fusion plasmid, pAMS366 (Stathopoulos and Cyert, 1997) (gifted by Dr. Martha Cyert, Stanford University) using the In-Fusion^®^ Cloning Kit (Clontech), resulting in p*P_YPK1_-LacZ* and p*P_LCB1_-LacZ* reporter plasmids, respectively.

### Genome editing for generation of the *P_YPK1_-*Δ*CBS* strain

The pMT1447 plasmid, constructed from the pSpCas9(BB)-2A-GFP plasmid (Ran et al., 2013) combined with the Z_4_EV expression system (McIsaac et al., 2013), co-expresses the nuclease protein Cas9 and gRNA, which guides the Cas9 protein with the target promoter sequence of *YPK1*. The repair DNA fragment was amplified from pNK19 carrying the *P_YPK1_-*Δ*CBS* fragment. The yeast wild-type strain was co-transformed with the pMT1603 plasmid and the repair DNA fragment using the lithium acetate transformation method. After selecting the uracil prototrophic transformants, the genomic DNA purified from the yeast transformants was used for genotyping by means of PCR using specific primers, to confirm the deletion of the CBS in the chromosomal *YPK1* promoter. The pMT1603 plasmid was lost by means of 5-FOA counter selection.

### Statistical analysis

Statistical analysis was performed using Prism 10 (GraphPad) or Excel (Microsoft) software, and the specific tests carried out have been mentioned in the text and figure legends.

## Acknowledgments

We express our deep appreciation to Prof. Scott D. Emr for generously providing the plasmids and strains. We thank Profs. Isamu Kameshita and Noriyuki Sueyoshi for providing the λPPase-expressing plasmid. We also thank Prof. Martha Cyert for providing the *LacZ* reporter plasmid, pAMS366. We also appreciate Prof. Takayuki Sekito for his technical advice on the experimental methods. We acknowledge the technical expertise of the DNA Core Facility of the Gene Research Center, Kagawa University. This work was supported by JSPS KAKENHI grant numbers JP19K05828, 23K26835 (to M.Tabuchi).

## Author contributions

Mitsuaki Tabuchi designed the research; Kosei Matsumoto, Ayane Nagai, Nao Komatsu, Yuko Ishino, Rina Shirai, Toshiya Ueno, Mio Masaki, and Mitsuaki Tabuchi performed the research; Kosei Matsumoto, Ayane Nagai, Nao Komatsu, Yuko Ishino, Rina Shirai, and Mitsuaki Tabuchi analyzed the data; Kosei Matsumoto, Ayane Nagai, and Mitsuaki Tabuchi wrote the manuscript; Motohiro Tani, Tatsuya Maeda, Naotaka Tanaka, and Ken-taro Sakata reviewed and edited the manuscript; Tatsuya Maeda provided resources; and Mitsuaki Tabuchi supervised the study and was responsible for funding acquisition and project administration.

## Notes

### Competing Interest Statement

The authors have declared no competing interest.

### Summary of Updates

We corrected the asterisk notation indicating the p-value in Figure 7J after re-checking the statistical analysis results and finding an error. In addition, for Figures 7G and 7H, we revised the corresponding figure legends because the statistical analysis had been performed using one-way ANOVA with Tukey's test, rather than two-way ANOVA with Sidak's test. We also re-uploaded the files in EPS format. Furthermore, we added Source Data #1 as a PDF file and Source Data #2 as a ZIP file containing compressed CSV files of the statistical analysis data for each figure.

## References

Ambroziak J, Henry SA. 1994. *INO2* and *INO*4 gene products, positive regulators of phospholipid biosynthesis in *Saccharomyces cerevisiae*, form a complex that binds to the *INO1* promoter. Journal of Biological Chemistry 269:15344–15349. DOI: 10.1016/S0021-9258(17)36612-7

Audhya A, Loewith R, Parsons AB, Gao L, Tabuchi M, Zhou H, Boone C, Hall MN, Emr SD. 2004. Genome-wide lethality screen identifies new PI4,5P2 effectors that regulate the actin cytoskeleton. The EMBO Journal 23:3747–3757. DOI: 10.1038/sj.emboj.7600384

Ballweg S, Sezgin E, Doktorova M, Covino R, Reinhard J, Wunnicke D, Hänelt I, Levental I, Hummer G, Ernst R. 2020. Regulation of lipid saturation without sensing membrane fluidity. Nature Communications 11:756. DOI: 10.1038/s41467-020-14528-1

Berchtold D, Piccolis M, Chiaruttini N, Riezman I, Riezman H, Roux A, Walther TC, Loewith R. 2012. Plasma membrane stress induces relocalization of Slm proteins and activation of TORC2 to promote sphingolipid synthesis. Nature Cell Biology 14:542–547. DOI: 10.1038/ncb2480

Breslow DK, Collins SR, Bodenmiller B, Aebersold R, Simons K, Shevchenko A, Ejsing CS, Weissman JS. 2010. Orm family proteins mediate sphingolipid homeostasis. Nature 463:1048–1053. DOI: 10.1038/nature08787

Breslow DK, Weissman JS. 2010. Membranes in Balance: Mechanisms of Sphingolipid Homeostasis. Molecular Cell 40:267–279. DOI: 10.1016/j.molcel.2010.10.005

Butera A, Agostini M, Cassandri M, De Nicola F, Fanciulli M, D’Ambrosio L, Falasca L, Nardacci R, Wang L, Piacentini M, Knight RA, Jia W, Sun Q, Shi Y, Wang Y, Candi E, Melino G. 2023. ZFP750 affects the cutaneous barrier through regulating lipid metabolism. Science Advances 9:eadg5423. DOI: 10.1126/sciadv.adg5423

Covino R, Hummer G, Ernst R. 2018. Integrated Functions of Membrane Property Sensors and a Hidden Side of the Unfolded Protein Response. Molecular Cell 71:458–467. DOI: 10.1016/j.molcel.2018.07.019

Eltschinger S, Loewith R. 2016. TOR Complexes and the Maintenance of Cellular Homeostasis. Trends in Cell Biology 26:148–159. DOI: 10.1016/j.tcb.2015.10.003

Finley D, Ulrich HD, Sommer T, Kaiser P. 2012. The Ubiquitin–Proteasome System of *Saccharomyces cerevisiae*. Genetics 192:319–360. DOI: 10.1534/genetics.112.140467

Geng F, Wenzel S, Tansey WP. 2012. Ubiquitin and Proteasomes in Transcription. Annual Review of Biochemistry 81:177–201. DOI: 10.1146/annurev-biochem-052110-120012

Goldstein JL, DeBose-Boyd RA, Brown MS. 2006. Protein Sensors for Membrane Sterols. Cell 124:35–46. DOI: 10.1016/j.cell.2005.12.022

Gururaj C, Federman R, Chang A. 2013. Orm Proteins Integrate Multiple Signals to Maintain Sphingolipid Homeostasis. Journal of Biological Chemistry 288:20453–20463. DOI: 10.1074/jbc.M113.472860

Harayama T, Riezman H. 2018. Understanding the diversity of membrane lipid composition. Nature Reviews Molecular Cell Biology 19:281–296. DOI: 10.1038/nrm.2017.138

Hua X, Sakai J, Brown MS, Goldstein JL. 1996. Regulated Cleavage of Sterol Regulatory Element Binding Proteins Requires Sequences on Both Sides of the Endoplasmic Reticulum Membrane. Journal of Biological Chemistry 271:10379–10384. DOI: 10.1074/jbc.271.17.10379

Ishino Y, Komatsu N, Sakata K, Yoshikawa D, Tani M, Maeda T, Morishige K, Yoshizawa K, Tanaka N, Tabuchi M. 2022. Regulation of sphingolipid biosynthesis in the endoplasmic reticulum via signals from the plasma membrane in budding yeast. The FEBS Journal 289:457–472. DOI: 10.1111/febs.16189

Kahana-Edwin S, Stark M, Kassir Y. 2013. Multiple MAPK Cascades Regulate the Transcription of *IME1*, the Master Transcriptional Activator of Meiosis in Saccharomyces cerevisiae. PLoS ONE 8:e78920. DOI: 10.1371/journal.pone.0078920

Kwiatek JM, Han G-S, Carman GM. 2020. Phosphatidate-mediated regulation of lipid synthesis at the nuclear/endoplasmic reticulum membrane. Biochimica et Biophysica Acta (BBA) - Molecular and Cell Biology of Lipids 1865:158434. DOI: 10.1016/j.bbalip.2019.03.006

Lage P, Sampaio-Marques B, Ludovico P, Mira NP, Mendes-Ferreira A. 2019. Transcriptomic and chemogenomic analyses unveil the essential role of Com2-regulon in response and tolerance of *Saccharomyces cerevisiae* to stress induced by sulfur dioxide. Microbial Cell 6:509–523. DOI: 10.15698/mic2019.11.697

Loewen CJR. 2003. A conserved ER targeting motif in three families of lipid binding proteins and in Opi1p binds VAP. The EMBO Journal 22:2025–2035. DOI: 10.1093/emboj/cdg201

Loewen CJR, Gaspar ML, Jesch SA, Delon C, Ktistakis NT, Henry SA, Levine TP. 2004. Phospholipid Metabolism Regulated by a Transcription Factor Sensing Phosphatidic Acid. Science 304:1644–1647. DOI: 10.1126/science.1096083

Longtine MS, Mckenzie Iii A, Demarini DJ, Shah NG, Wach A, Brachat A, Philippsen P, Pringle JR. 1998. Additional modules for versatile and economical PCR-based gene deletion and modification in *Saccharomyces cerevisiae*. Yeast 14:953–961. DOI: 10.1002/(SICI)1097-0061(199807)14:10%3C953::AID-YEA293%3E3.0.CO;2-U

McIsaac RS, Oakes BL, Wang X, Dummit KA, Botstein D, Noyes MB. 2013. Synthetic gene expression perturbation systems with rapid, tunable, single-gene specificity in yeast. Nucleic Acids Research 41:e57–e57. DOI: 10.1093/nar/gks1313

Meimoun A, Holtzman T, Weissman Z, McBride HJ, Stillman DJ, Fink GR, Kornitzer D. 2000. Degradation of the Transcription Factor Gcn4 Requires the Kinase Pho85 and the SCFCDC4 Ubiquitin–Ligase Complex. Molecular Biology of the Cell 11:915–927. DOI: 10.1091/mbc.11.3.915, PMID: 10712509

Muir A, Ramachandran S, Roelants FM, Timmons G, Thorner J. 2014. TORC2-dependent protein kinase Ypk1 phosphorylates ceramide synthase to stimulate synthesis of complex sphingolipids. eLife 3:e03779. DOI: 10.7554/eLife.03779

Nabeel-Shah S, Pu S, Burns JD, Braunschweig U, Ahmed N, Burke GL, Lee H, Radovani E, Zhong G, Tang H, Marcon E, Zhang Z, Hughes TR, Blencowe BJ, Greenblatt JF. 2024. C2H2-zinc-finger transcription factors bind RNA and function in diverse post-transcriptional regulatory processes. Molecular Cell 84:3810–3825.e10. DOI: 10.1016/j.molcel.2024.08.037

Niles BJ, Mogri H, Hill A, Vlahakis A, Powers T. 2012. Plasma membrane recruitment and activation of the AGC kinase Ypk1 is mediated by target of rapamycin complex 2 (TORC2) and its effector proteins Slm1 and Slm2. Proceedings of the National Academy of Sciences 109:1536–1541. DOI: 10.1073/pnas.1117563109

Nohturfft A, Brown MS, Goldstein JL. 1998. Sterols regulate processing of carbohydrate chains of wild-type SREBP cleavage-activating protein (SCAP), but not sterol-resistant mutants Y298C or D443N. Proceedings of the National Academy of Sciences 95:12848–12853. DOI: 10.1073/pnas.95.22.12848

Nohturfft A, DeBose-Boyd RA, Scheek S, Goldstein JL, Brown MS. 1999. Sterols regulate cycling of SREBP cleavage-activating protein (SCAP) between endoplasmic reticulum and Golgi. Proceedings of the National Academy of Sciences 96:11235–11240. DOI: 10.1073/pnas.96.20.11235

Nohturfft A, Yabe D, Goldstein JL, Brown MS, Espenshade PJ. 2000. Regulated Step in Cholesterol Feedback Localized to Budding of SCAP from ER Membranes. Cell 102:315–323. DOI: 10.1016/S0092-8674(00)00037-4

Nohturfft A, Zhang SC. 2009. Coordination of Lipid Metabolism in Membrane Biogenesis. Annual Review of Cell and Developmental Biology 25:539–566. DOI: 10.1146/annurev.cellbio.24.110707.175344

Ran FA, Hsu PD, Wright J, Agarwala V, Scott DA, Zhang F. 2013. Genome engineering using the CRISPR-Cas9 system. Nature Protocols 8:2281–2308. DOI: 10.1038/nprot.2013.143

Rawson RB, Zelenski NG, Nijhawan D, Ye J, Sakai J, Hasan MT, Chang TY, Brown MS, Goldstein JL. 1997. Complementation Cloning of, a Gene Encoding a Putative Metalloprotease Required for Intramembrane Cleavage of SREBPs. Molecular Cell 1:47–57. DOI: 10.1016/S1097-2765(00)80006-4

Robinson JS, Klionsky DJ, Banta LM, Emr SD. 1988. Protein sorting in Saccharomyces cerevisiae: isolation of mutants defective in the delivery and processing of multiple vacuolar hydrolases. Molecular and Cellular Biology 8:4936–4948. DOI: 10.1128/mcb.8.11.4936-4948.1988, PMID: 3062374

Roelants FM, Breslow DK, Muir A, Weissman JS, Thorner J. 2011. Protein kinase Ypk1 phosphorylates regulatory proteins Orm1 and Orm2 to control sphingolipid homeostasis in *Saccharomyces cerevisiae*. Proceedings of the National Academy of Sciences 108:19222–19227. DOI: 10.1073/pnas.1116948108

Romanauska A, Köhler A. 2021. Reprogrammed lipid metabolism protects inner nuclear membrane against unsaturated fat. Developmental Cell 56:2562–2578.e3. DOI: 10.1016/j.devcel.2021.07.018

Sakai J, Duncan EA, Rawson RB, Hua X, Brown MS, Goldstein JL. 1996. Sterol-Regulated Release of SREBP-2 from Cell Membranes Requires Two Sequential Cleavages, One Within a Transmembrane Segment. Cell 85:1037–1046. DOI: 10.1016/S0092-8674(00)81304-5

Sakai J, Rawson RB, Espenshade PJ, Cheng D, Seegmiller AC, Goldstein JL, Brown MS. 1998. Molecular Identification of the Sterol-Regulated Luminal Protease that Cleaves SREBPs and Controls Lipid Composition of Animal Cells. Molecular Cell 2:505–514. DOI: 10.1016/S1097-2765(00)80150-1

Siggers T, Reddy J, Barron B, Bulyk ML. 2014. Diversification of Transcription Factor Paralogs via Noncanonical Modularity in C2H2 Zinc Finger DNA Binding. Molecular Cell 55:640–648. DOI: 10.1016/j.molcel.2014.06.019

Sikorski RS, Hieter P. 1989. A System of Shuttle Vectors and Yeast Host Strains Designed for Efficient Manipulation of DNA in Saccharomyces ceratisiae. Genetics 122:19–27.

Stathopoulos AM, Cyert MS. 1997. Calcineurin acts through the *CRZ1/TCN1* - encoded transcription factor to regulate gene expression in yeast. Genes & Development 11:3432–3444. DOI: 10.1101/gad.11.24.3432

Tabuchi M, Audhya A, Parsons AB, Boone C, Emr SD. 2006. The Phosphatidylinositol 4,5-Biphosphate and TORC2 Binding Proteins Slm1 and Slm2 Function in Sphingolipid Regulation. Molecular and Cellular Biology 26:5861–5875. DOI: 10.1128/MCB.02403-05

van Meer G, Voelker DR, Feigenson GW. 2008. Membrane lipids: where they are and how they behave. Nature reviews. Molecular cell biology 9:112–124. DOI: 10.1038/nrm2330, PMID: 18216768

Wang X, Sato R, Brown MS, Hua X, Goldstein JL. 1994. SREBP-1, a membrane-bound transcription factor released by sterol-regulated proteolysis. Cell 77:53–62. DOI: 10.1016/0092-8674(94)90234-8

White MJ, Hirsch JP, Henry SA. 1991. The *OPI1* gene of Saccharomyces cerevisiae, a negative regulator of phospholipid biosynthesis, encodes a protein containing polyglutamine tracts and a leucine zipper. Journal of Biological Chemistry 266:863–872. DOI: 10.1016/S0021-9258(17)35253-5

Yamaguchi Y, Katsuki Y, Tanaka S, Kawaguchi R, Denda H, Ikeda T, Funato K, Tani M. 2018. Protective role of the HOG pathway against the growth defect caused by impaired biosynthesis of complex sphingolipids in yeast *Saccharomyces cerevisiae*. Molecular Microbiology 107:363–386. DOI: 10.1111/mmi.13886

Yang H, Tong J, Lee CW, Ha S, Eom SH, Im YJ. 2015. Structural mechanism of ergosterol regulation by fungal sterol transcription factor Upc2. Nature Communications 6:6129. DOI: 10.1038/ncomms7129

