## Supplemental Data 1 for "Regulation of sphingolipid synthesis by the C2H2 zinc finger transcription factor Com2 through ubiquitin-proteasome mediated degradation pathway"

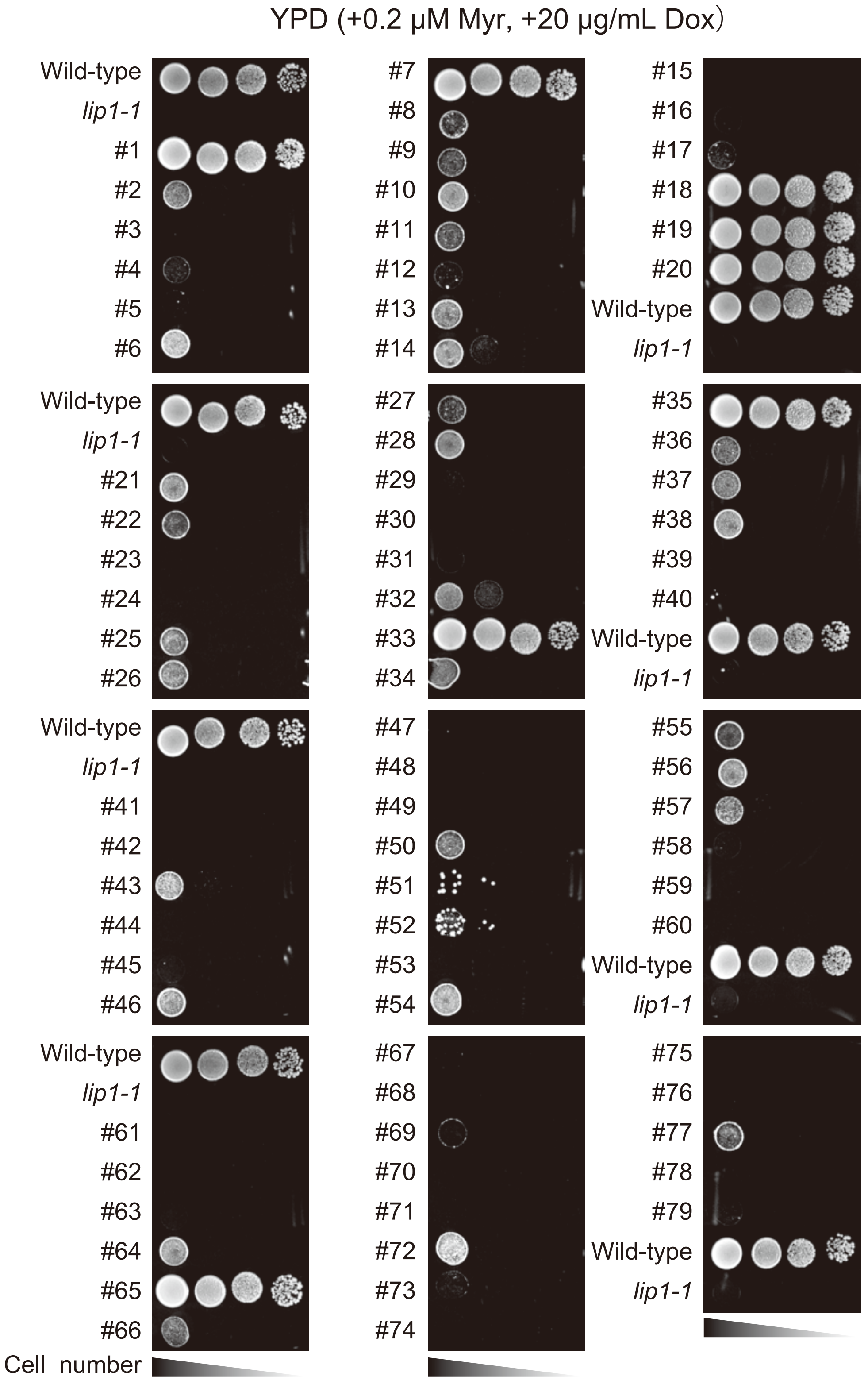


*MLM1* (*LIP1*) : #1, #7, #18, #19, #20, #33, #35, #65
*MLM2* (*COM2*) : #21, #34, #38, #46, #55, #64, #77
*MLM3* (*TIF3*) : #11, #13
*MLM4* (*STM1*) : #50, #54
*MLM5* (*YPK1*) : #6
*MLM6* (*LSP1*) : #9
*MLM7* (*BMH2*) : #17
*MLM8* (*RIM20* (partial), *CAF20*, *HEM4*, *RFM1*) : #26
*MLM9* (*TVP18*): #28
*MLM10* (*SRO77* (partial), *PKC1* (partial)): #36
*MLM11* (*PIN4*): #56
*MLM12* (*SUT2*): #66
