## Supplementary figures and images for "Regulation of sphingolipid synthesis by the C2H2 zinc finger transcription factor Com2 through ubiquitin-proteasome mediated degradation pathway"

### Supplemental Data 2

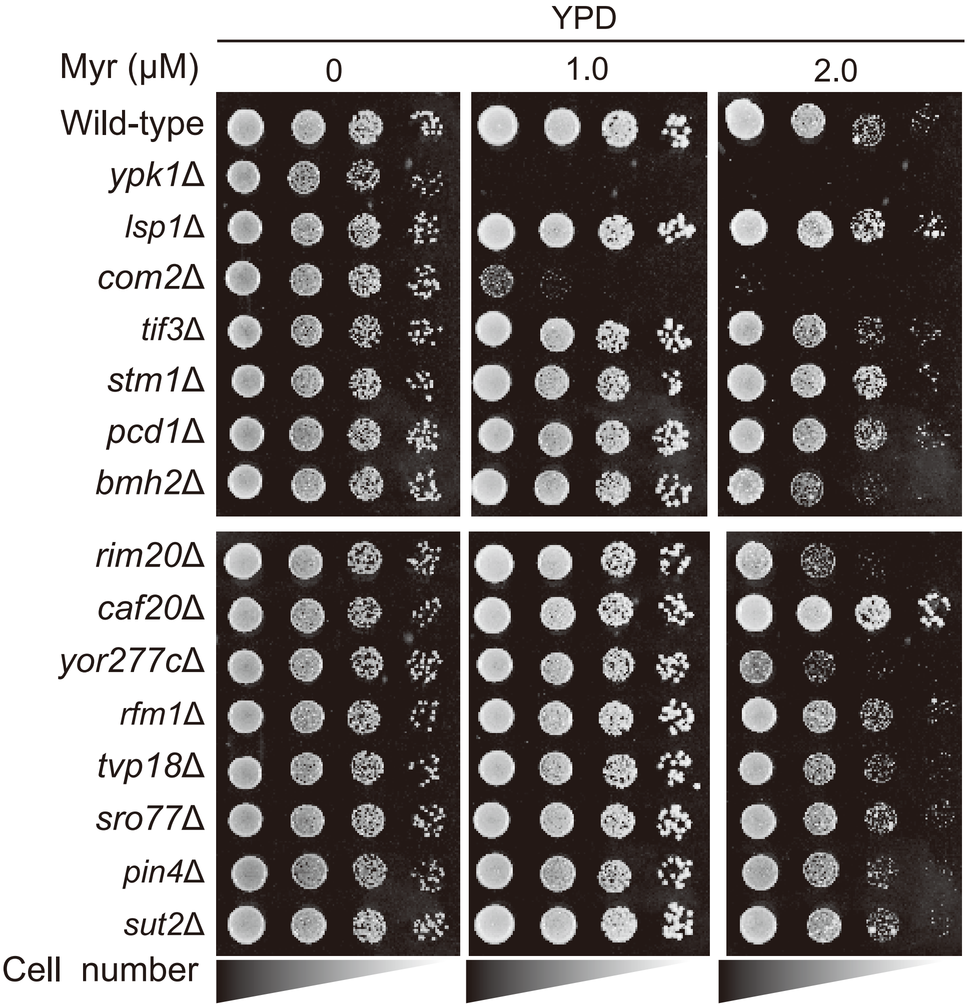

### Supplemental Data 3

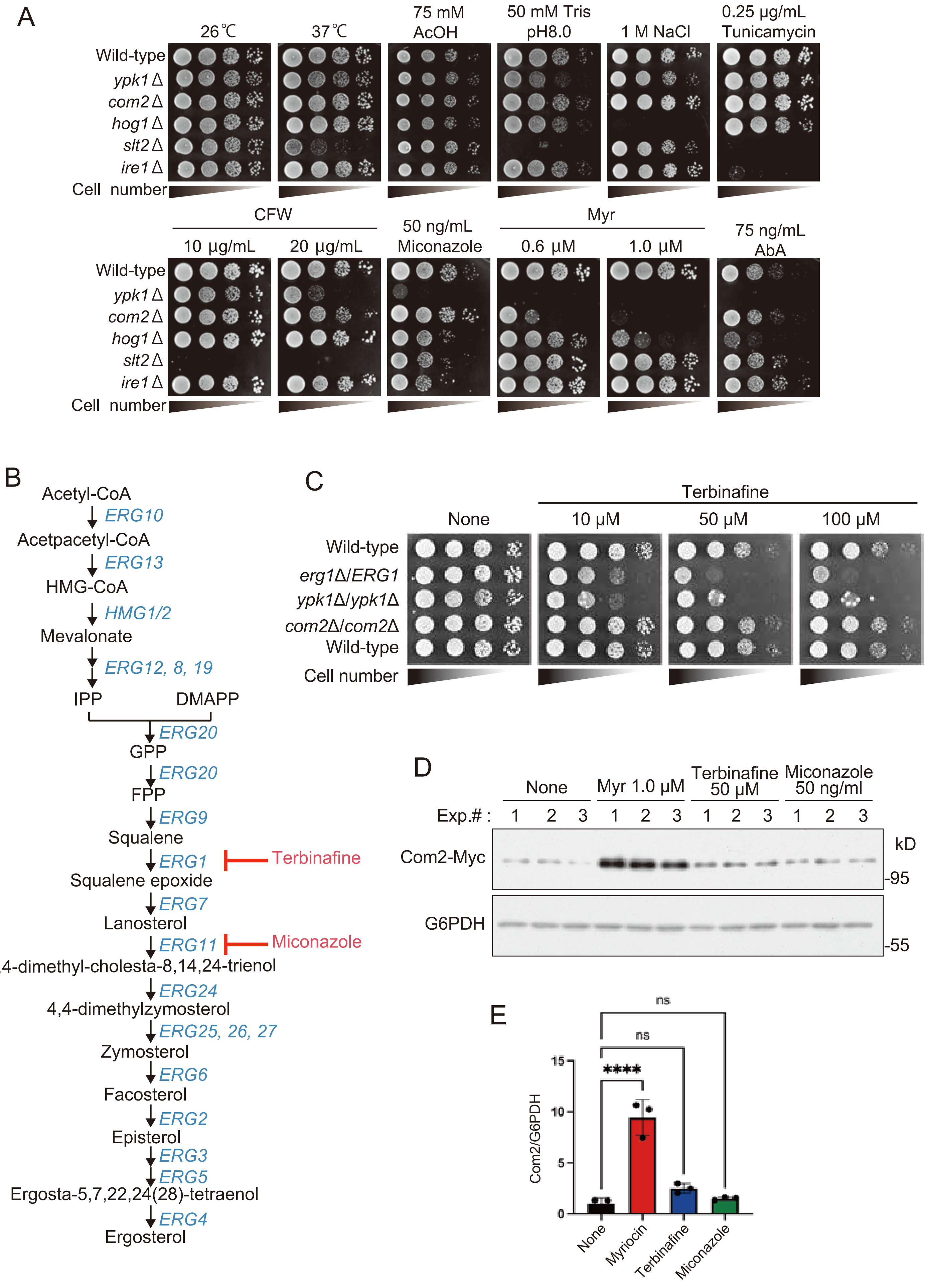

### Supplemental Data 4

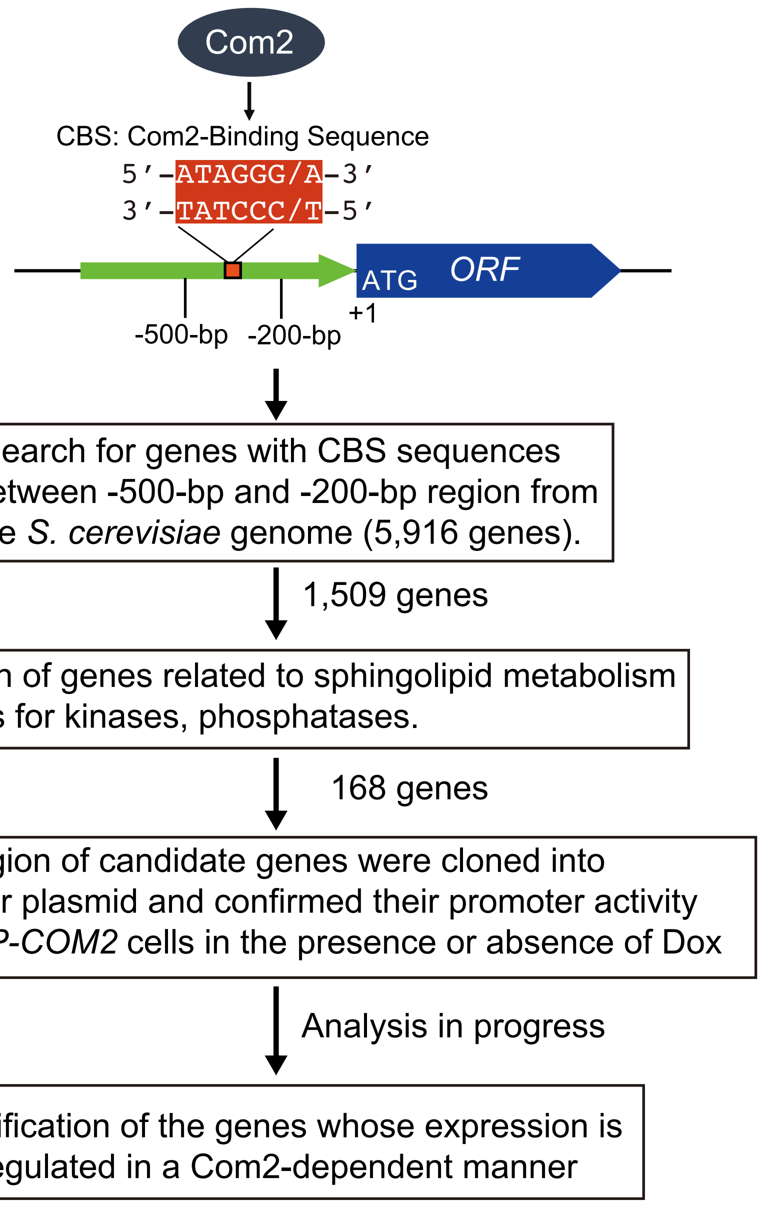

### Supplemental Data 5

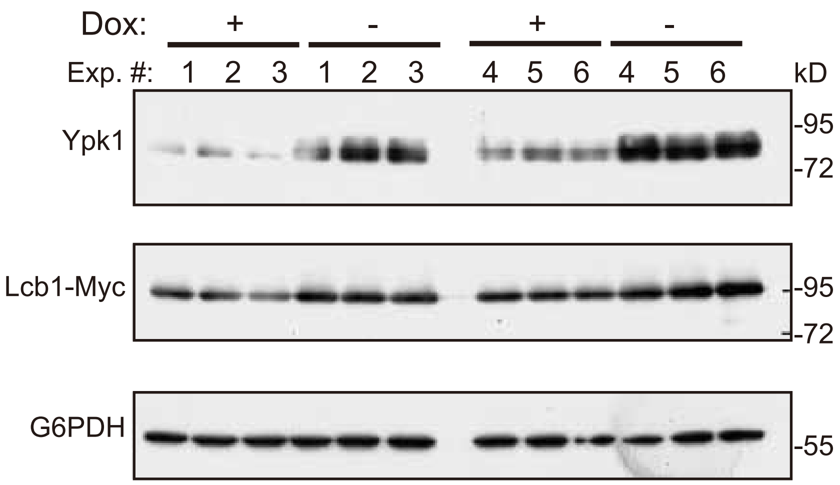

### Supplemental Data 6

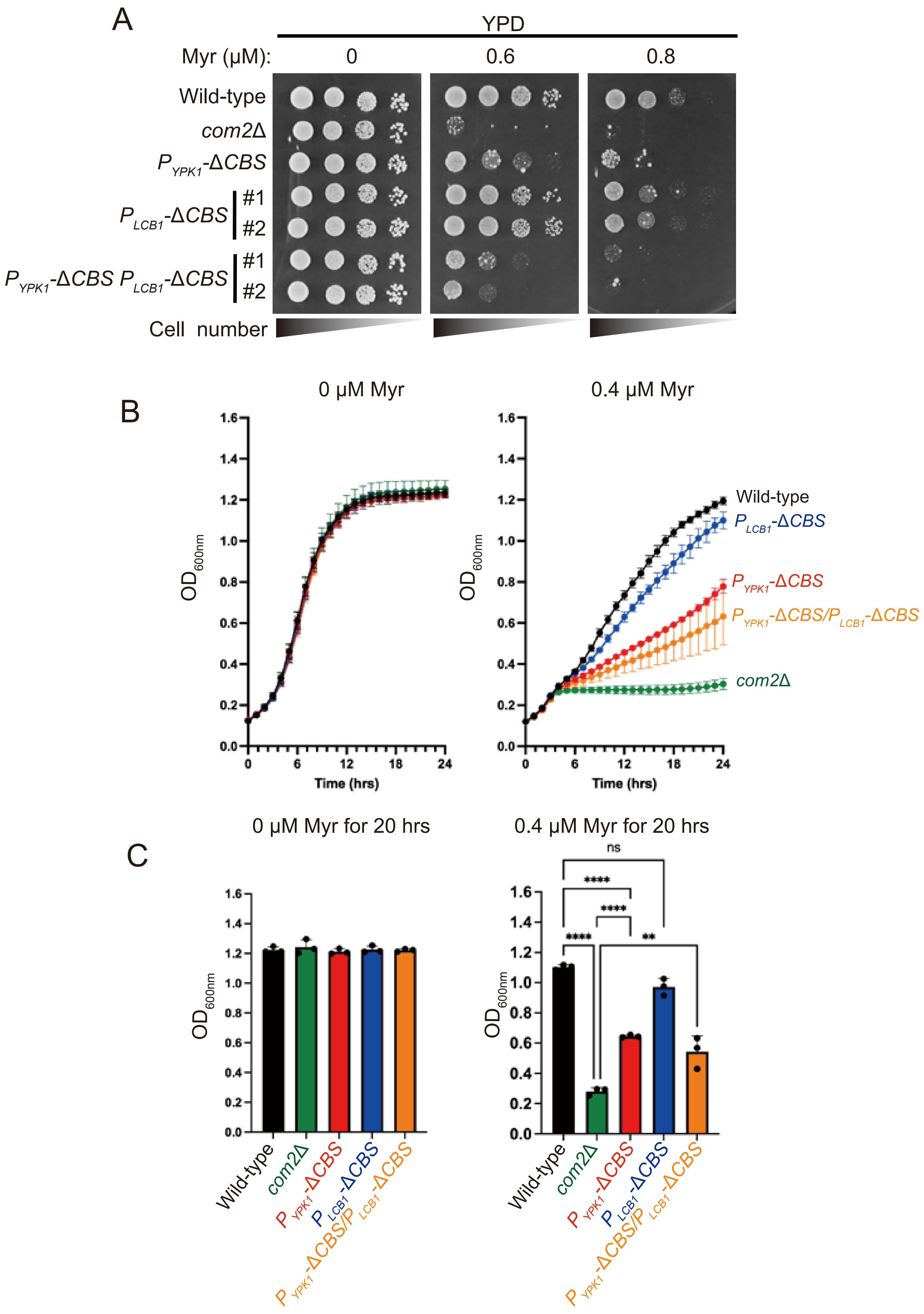

### Supplemental Data 7

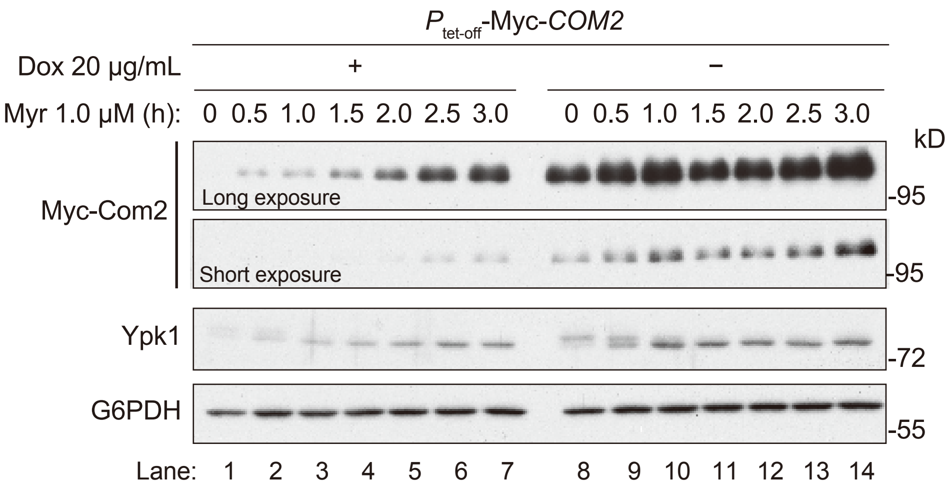

### Supplemental Data 8

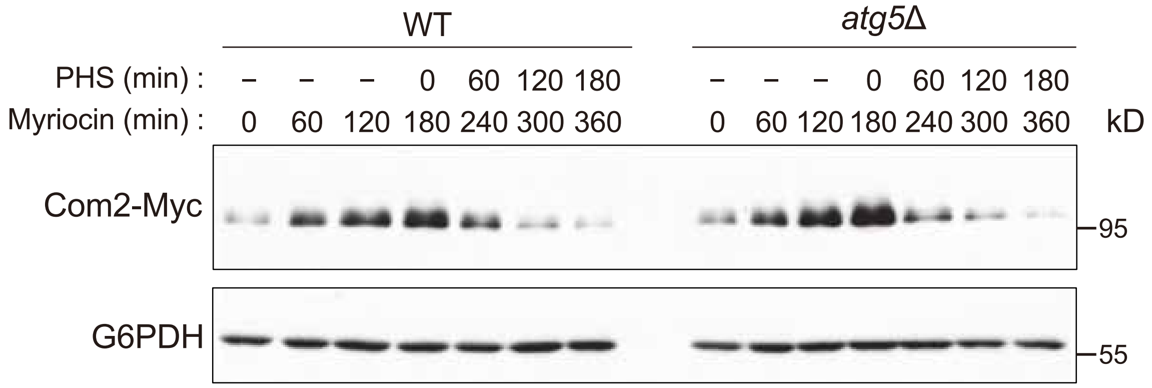
