## Supplemental Data 9 for "Regulation of sphingolipid synthesis by the C2H2 zinc finger transcription factor Com2 through ubiquitin-proteasome mediated degradation pathway"

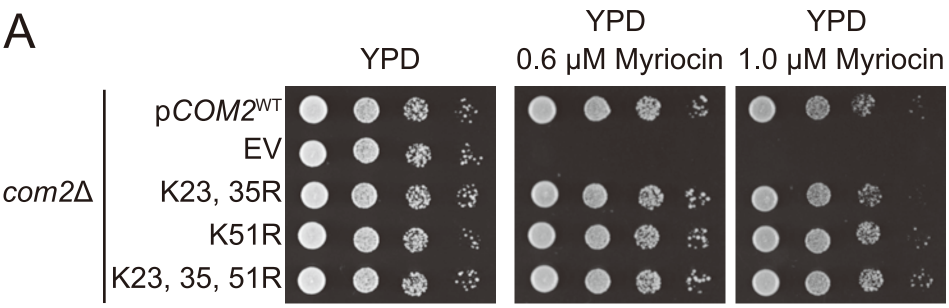


B

| position | Sequence | SGD* | Ypk1** | SGK1*** | SGK3*** | SGK*** | AGC*** | ST-A3 | ST-A5 | ST-A7 | ST-A8 | ST-A10 |
|---|---|---|---|---|---|---|---|---|---|---|---|---|
| 11S | YPLQRFE**S**NDTVFSY | + | | | | | | | | | | + |
| 88S | NQLERRL**S**ISDYRTE | + | | | + | | + | | + | + | + | + |
| 138S | ESDVLMV**S**DDELEVN | + | | | | | | | | | | + |
| 164S | DGLNRIS**S**TNNLKNL | + | | | + | | | | + | + | + | + |
| 235S | PIQTENS**S**SQKMFKN | | | + | | + | | | | + | + | + |
| 251T | FFRSRKS**T**LIKSLPL | | + | | + | + | + | + | + | + | + | + |
| 359S | SVQSSSS**S**HGLVVRK | | | | | + | | | | + | + | + |
| 368T | GLVVRKK**T**GSMQKTR | | | + | | | + | | | | + | + |
| 370S | VVRKKTG**S**MQKTRGR | | + | + | + | + | + | + | + | + | + | + |
| 380S | KTRGRKP**S**LIPDASK | + | + | | | + | + | + | + | + | + | + |

*SGD: https://www.yeastgenome.org/locus/S000000932/protein

**Muri et al., eLife: https://elifesciences.org/articles/03779

***GPS web server : http://gps.biocuckoo.org/online.php

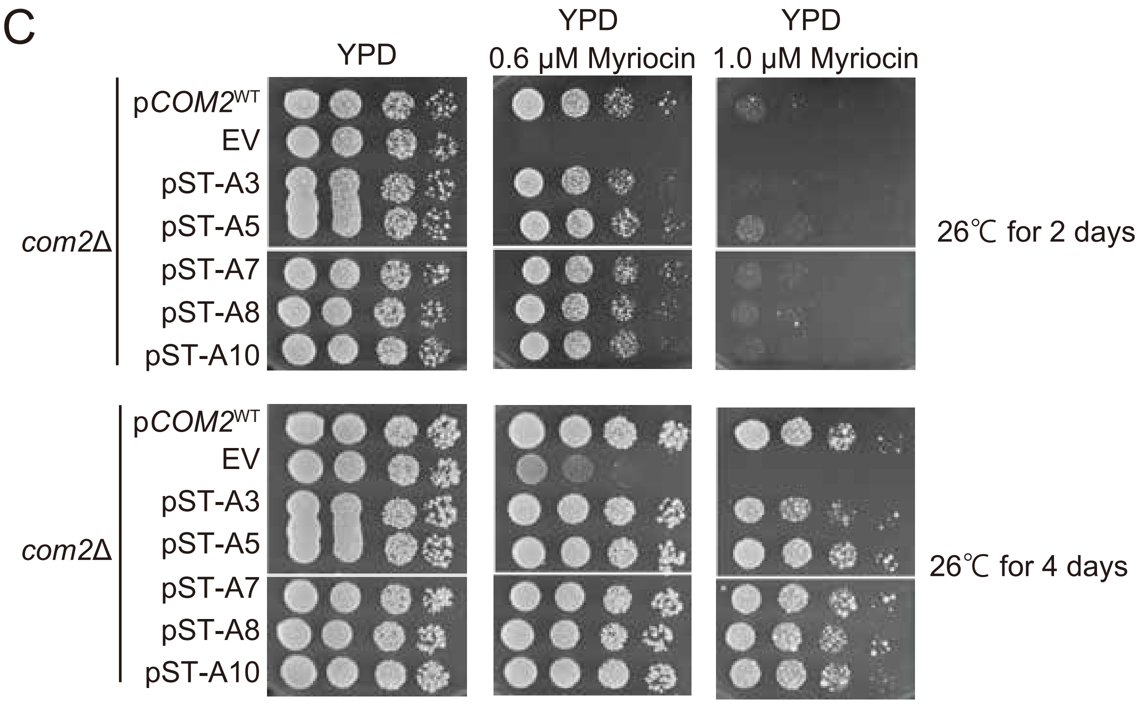
