## Supplementary material for "Regulation of sphingolipid synthesis by the C2H2 zinc finger transcription factor Com2 through ubiquitin-proteasome mediated degradation pathway": Source data-1

### Figure 1D original data

YPD

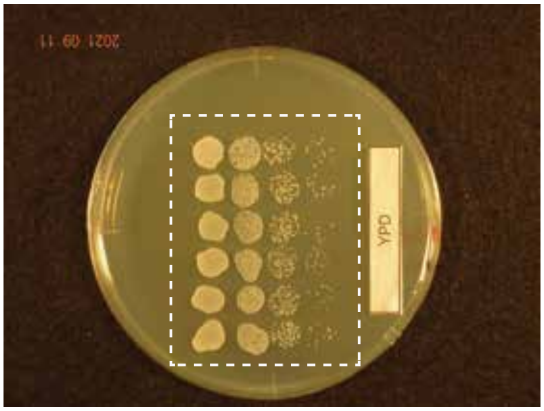


YPD + Dox (20 μg/mL)

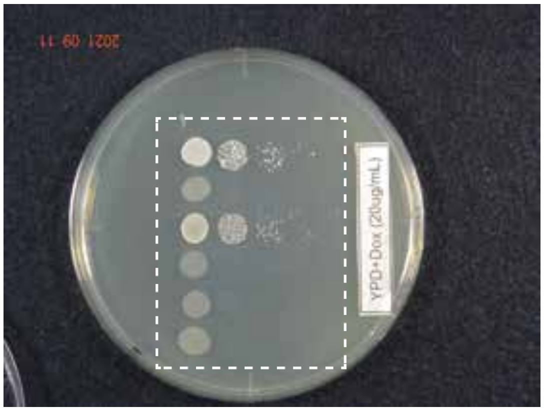


YPD + 0.2 μM Myriocin

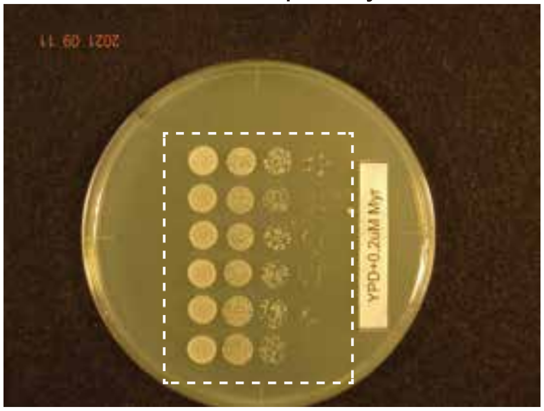


YPD + Dox (20 μg/mL), 0.2 μM Myriocin

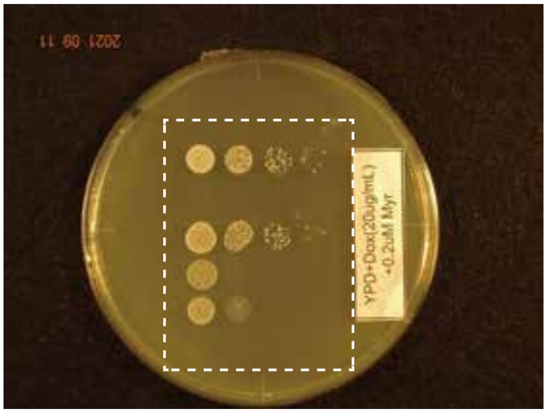

### Figure 1E source data

YPD

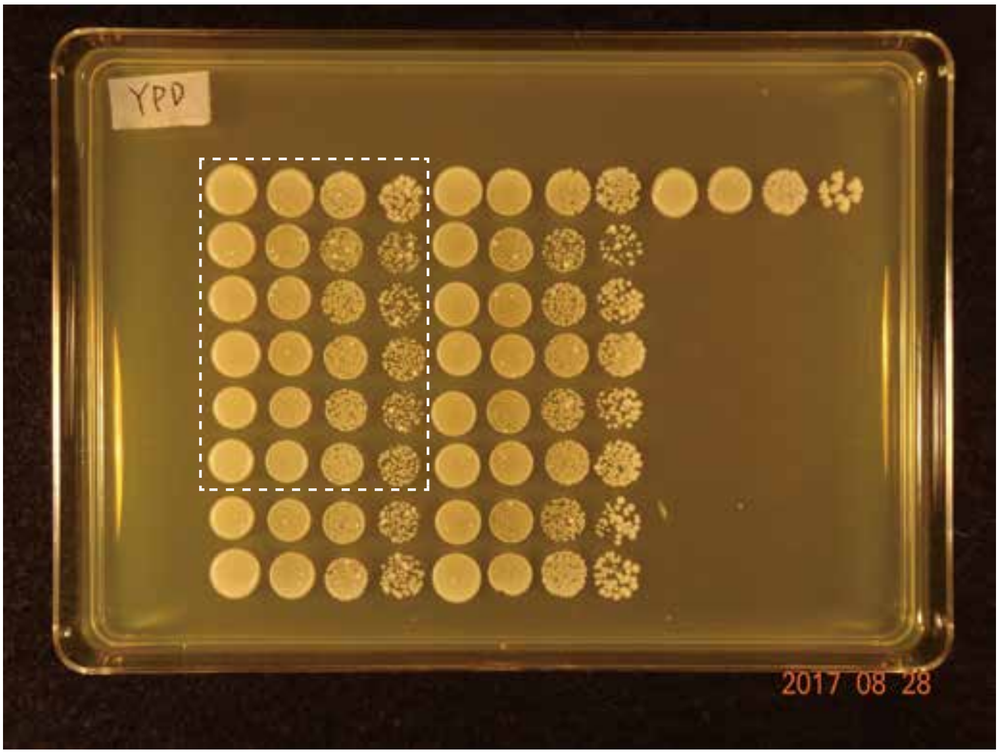


YPD + 1

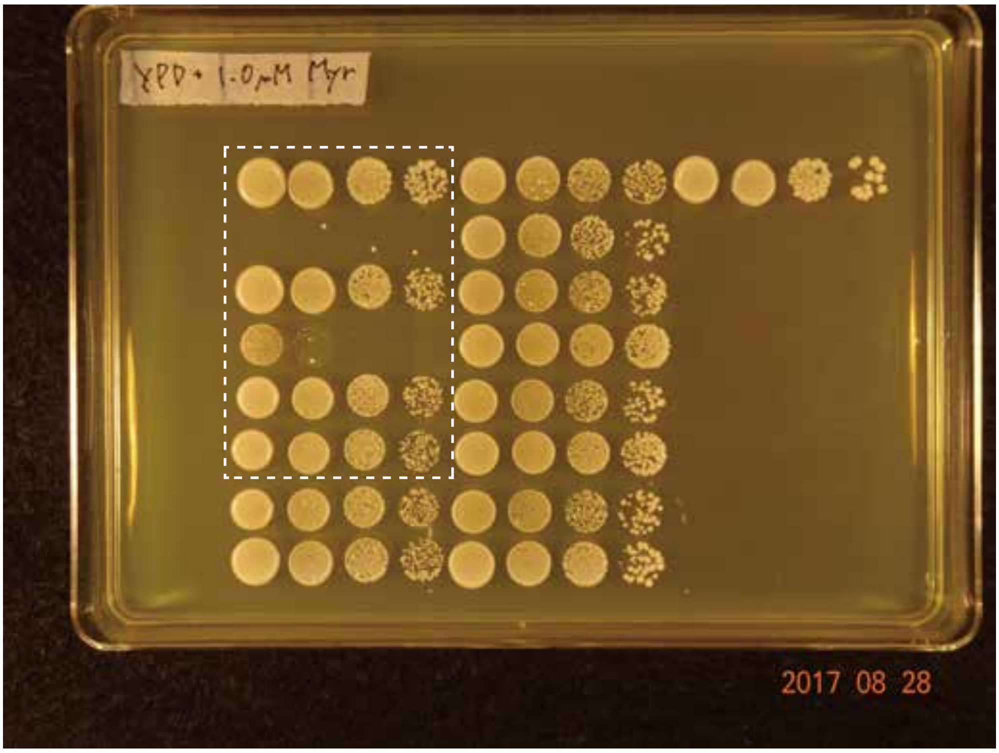

Figure 1F, G source data

n1, processed for Fig. 1F

αMyc

αG6PDH

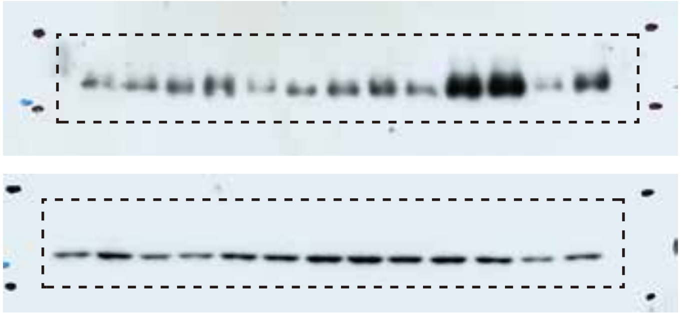

n2

αMyc

αG6PDH

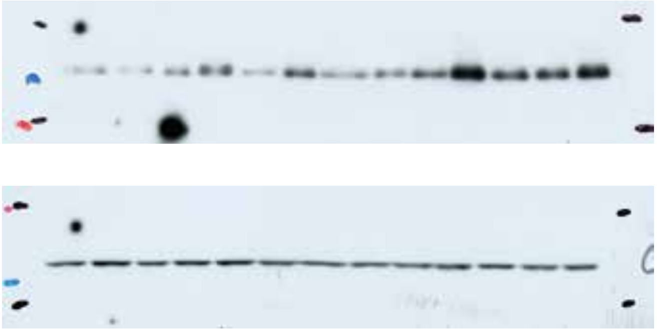

n3

αMyc

αG6PDH

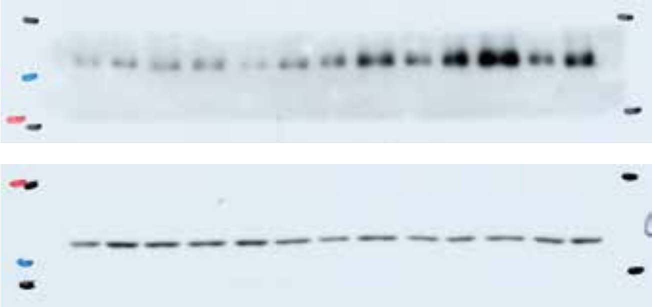

### Figure 1-figure supplement 3A source data

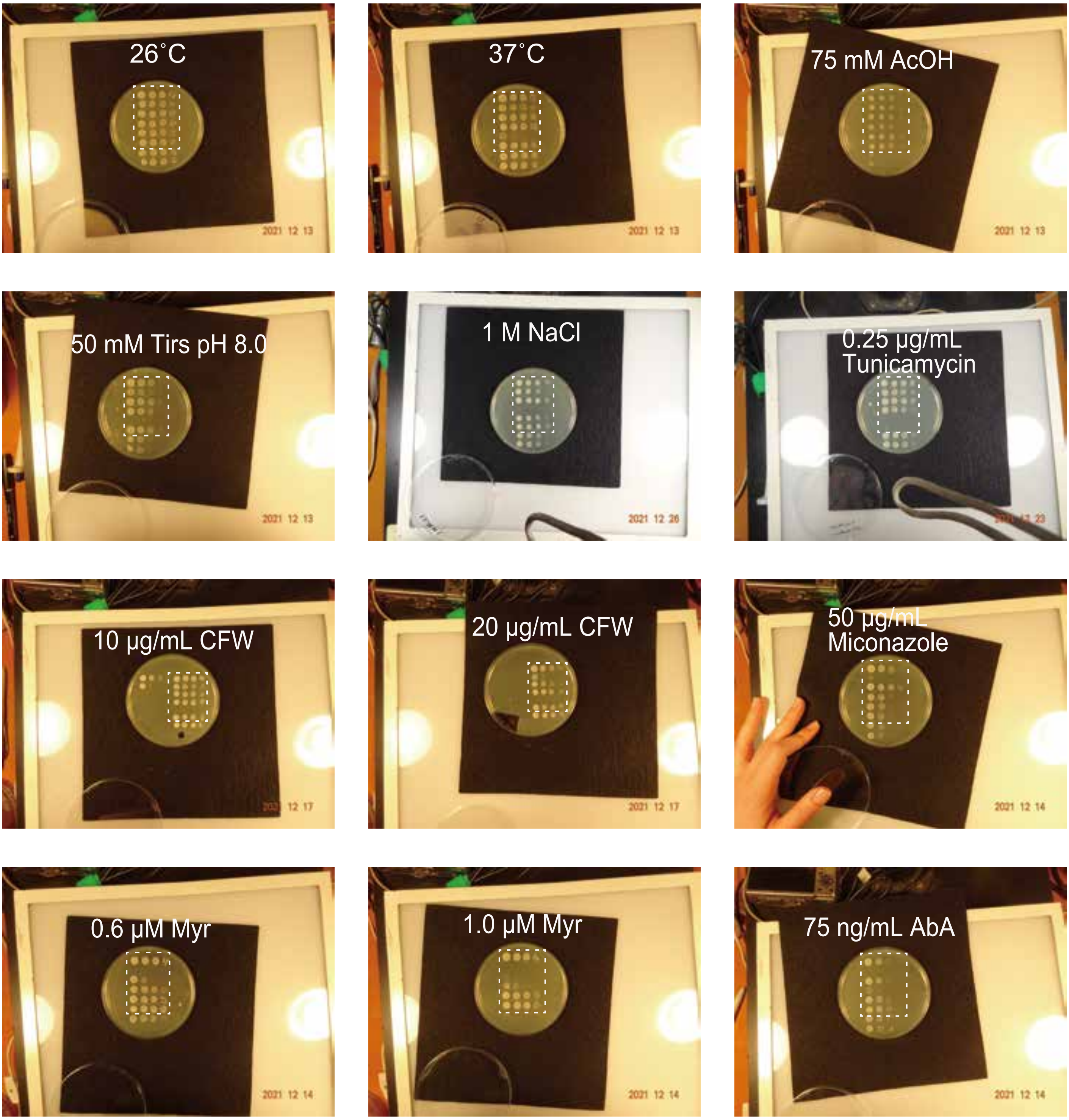

Figure 1-figure supplement 3C source data

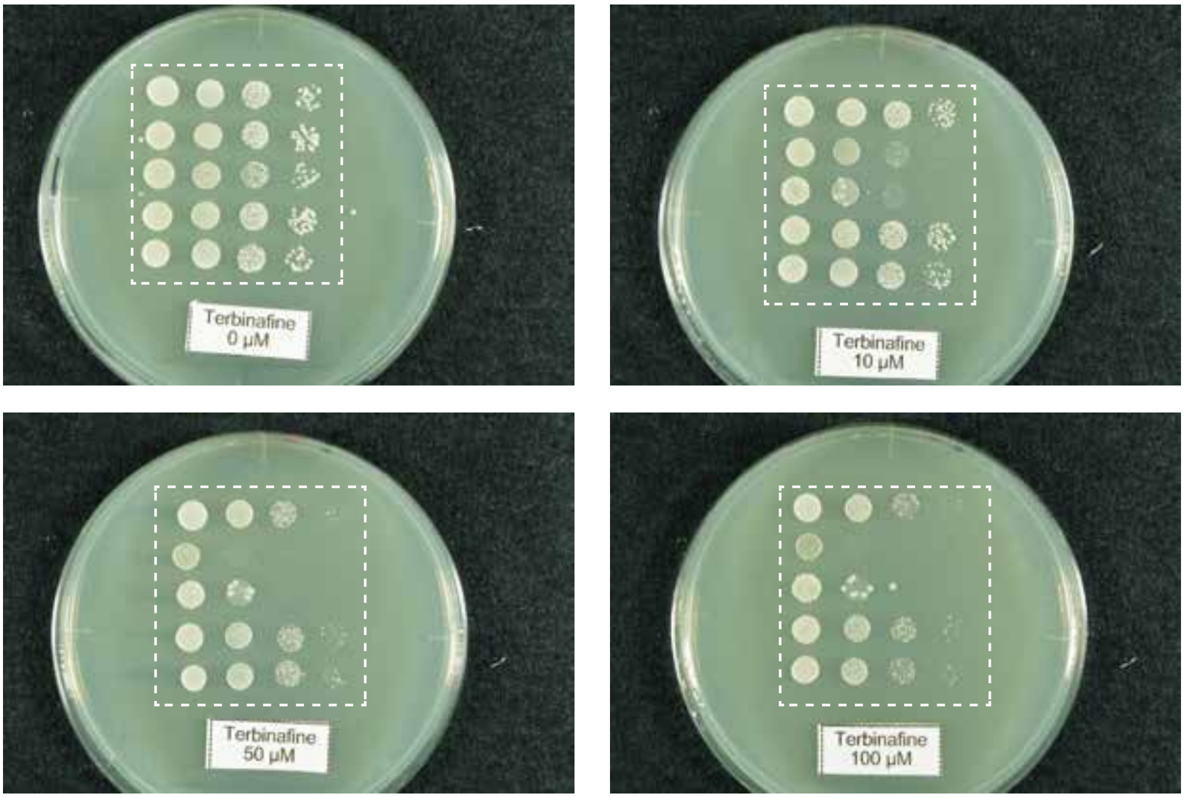

Figure 1-figure supplement 3D source data

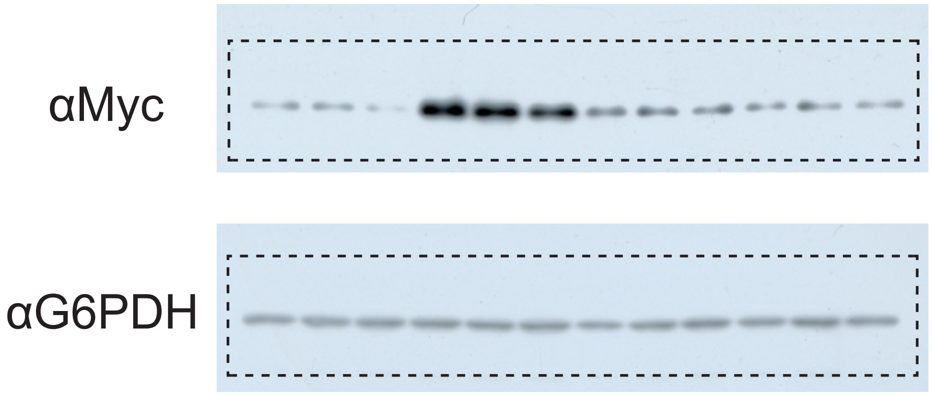

### Figure 2 source data

#### 2B original data

Mock

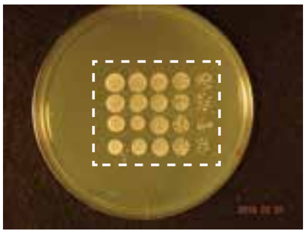

3-MB-PP1

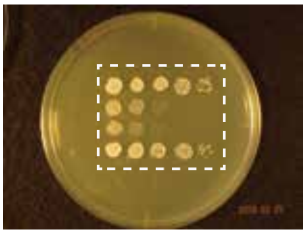

#### 2C original data

Mock

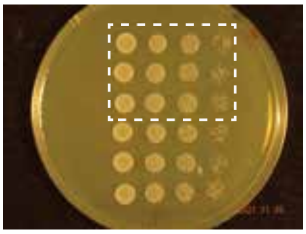

0.7 μM Myr

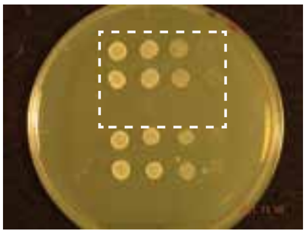

#### 2C original data

αMyc

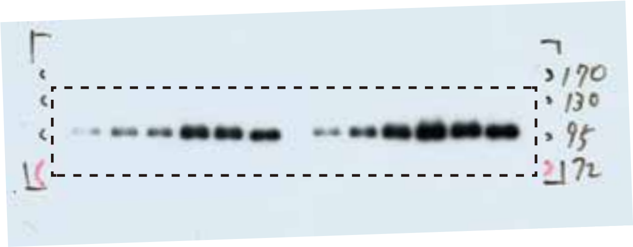


αMyc

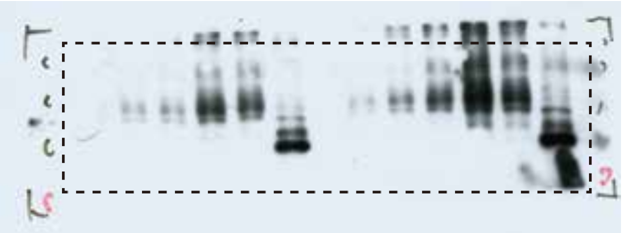

αFLAG

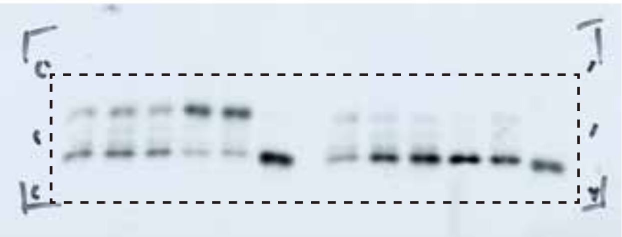

αG6PDH

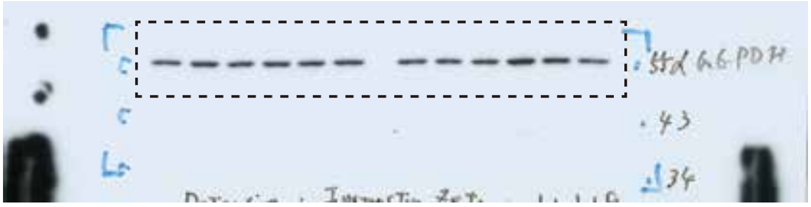


#### 2E original data

None

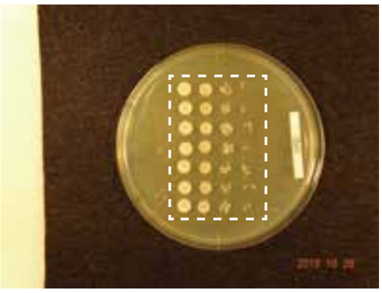

1.0 μM Myr

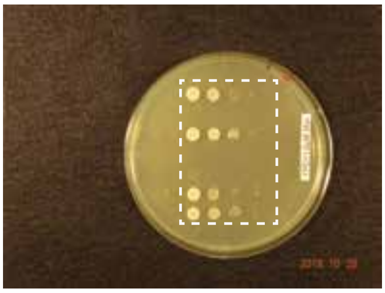

1.0 μM Myr

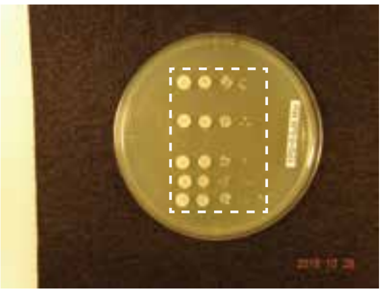

### Figure 3B source data

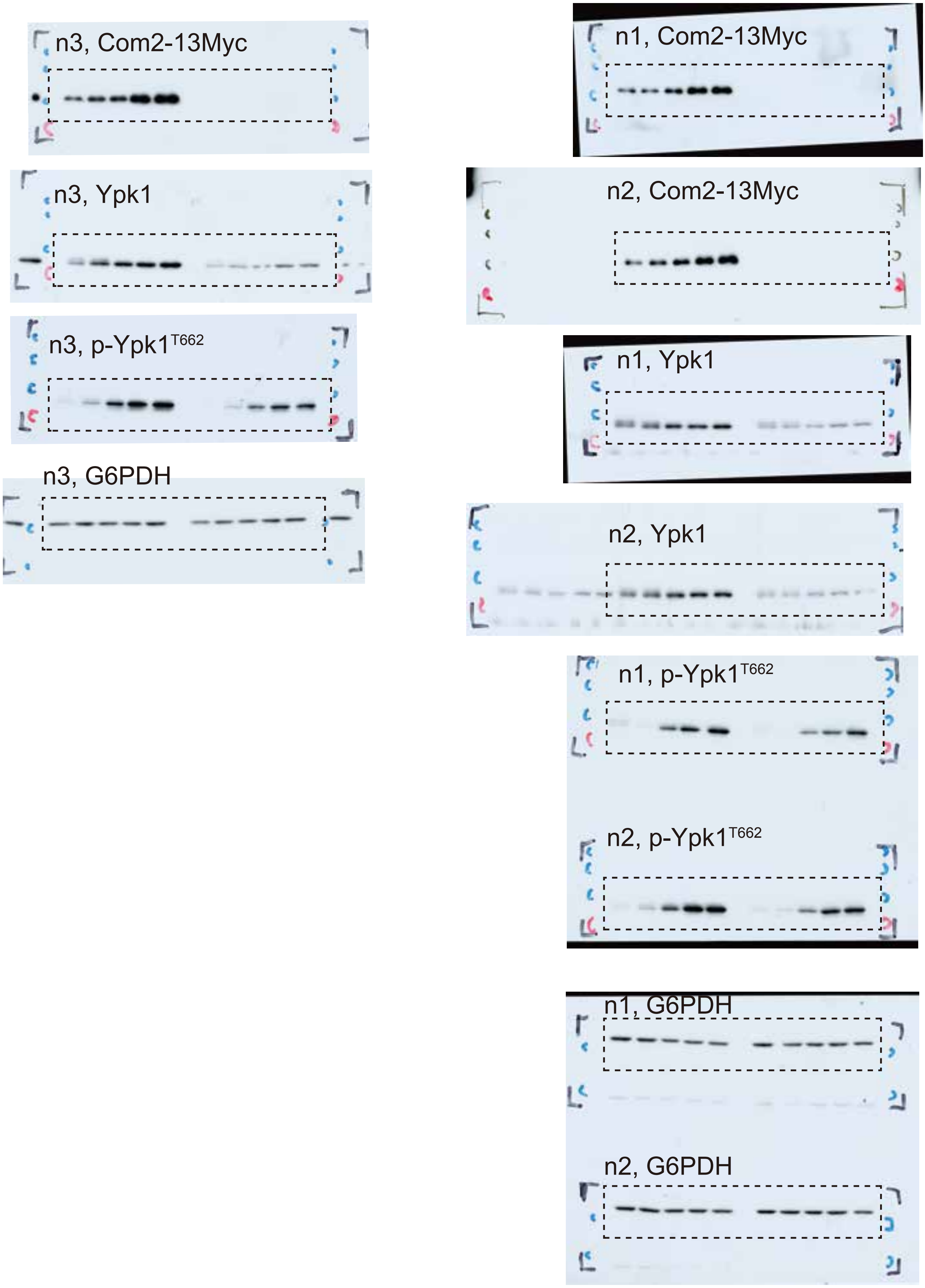

Figure 3G source data

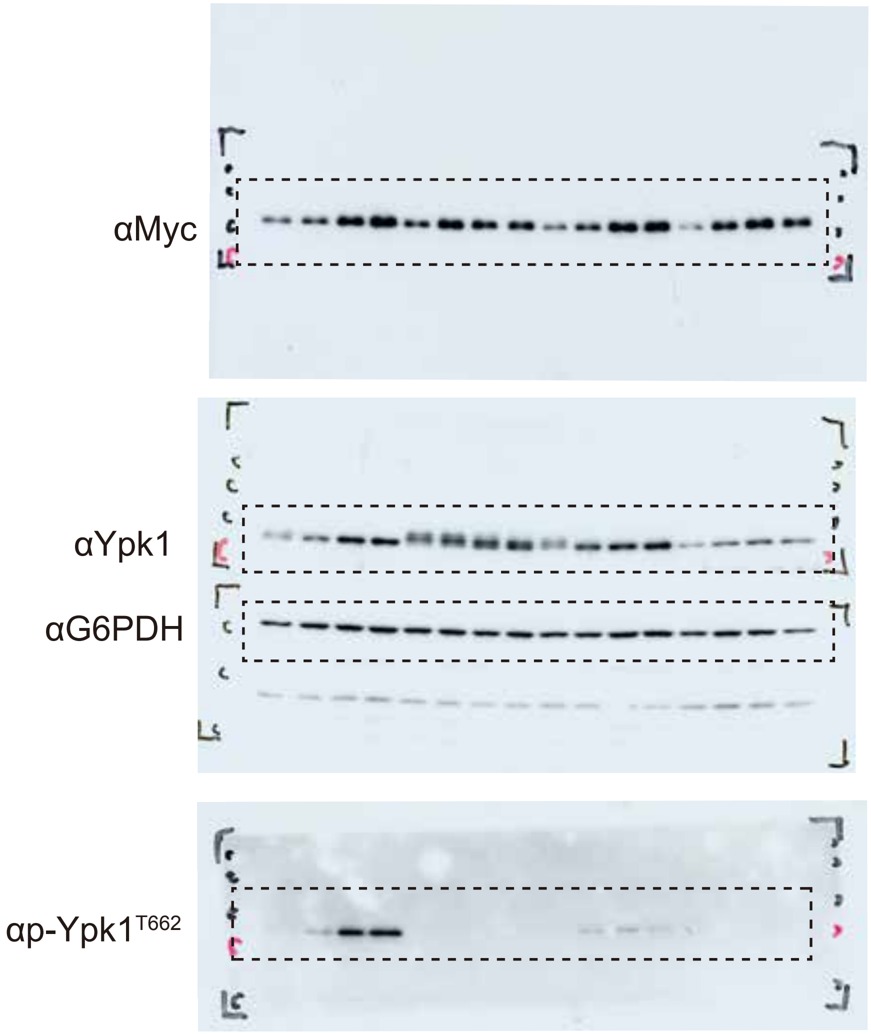

### Figure 4B source data

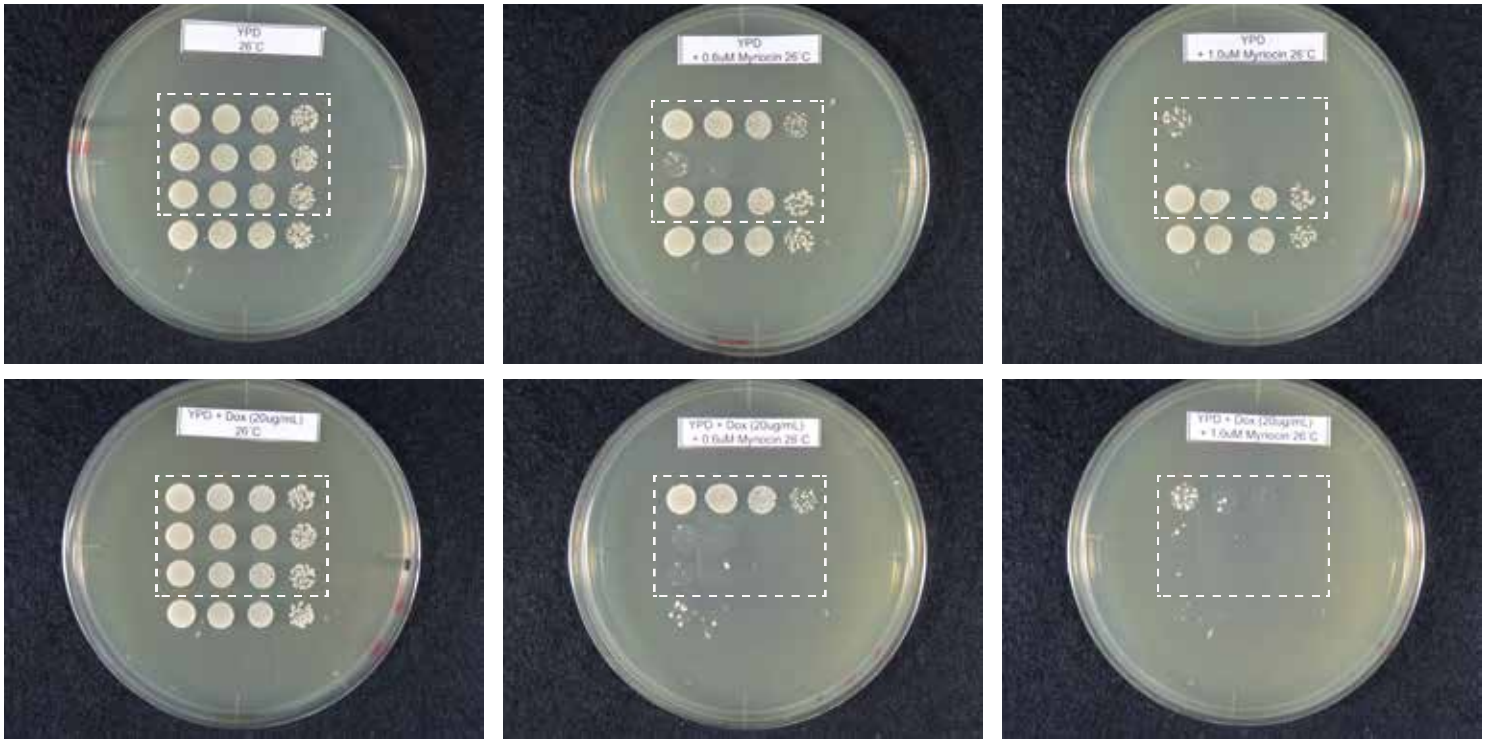

### Figure 4D source data

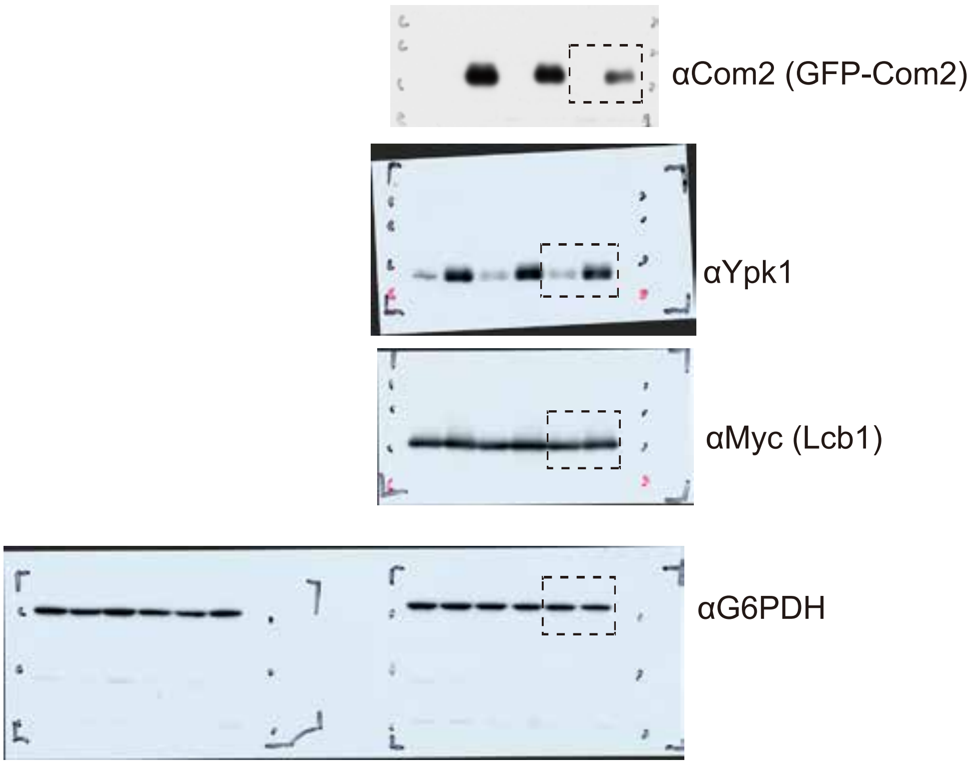

Figure 5B source data

Myr 0 μM

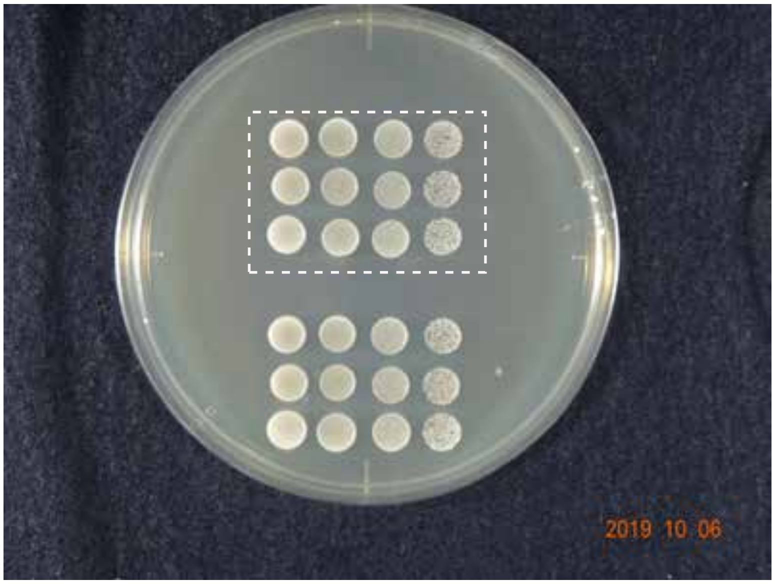


Myr 0.6 μM

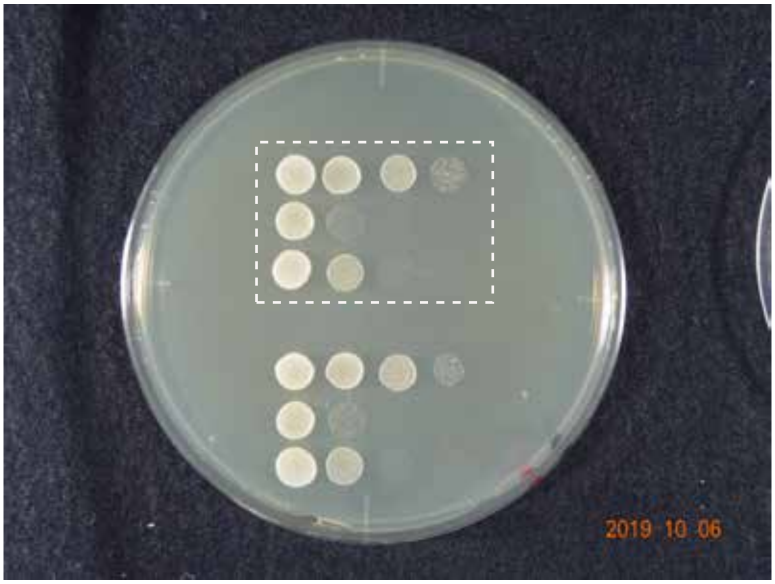

Figure 4-figure supplement 2 source data

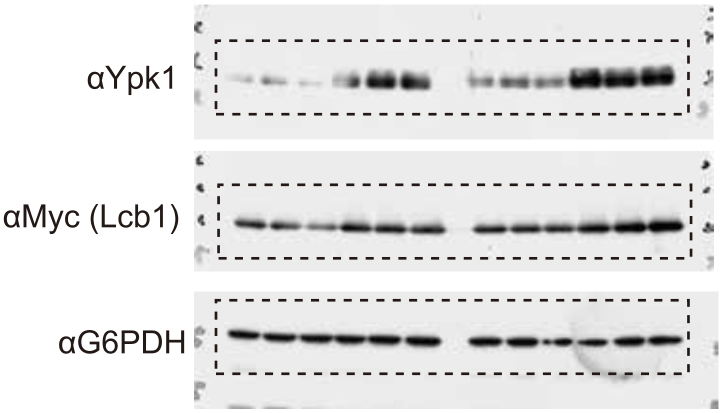

Figure 5D source data

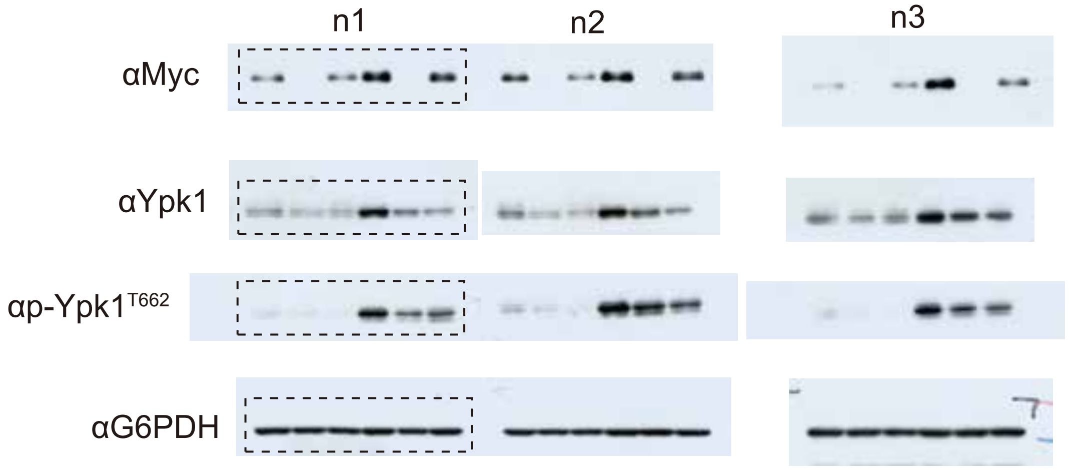

### Figure 5H source data

n1

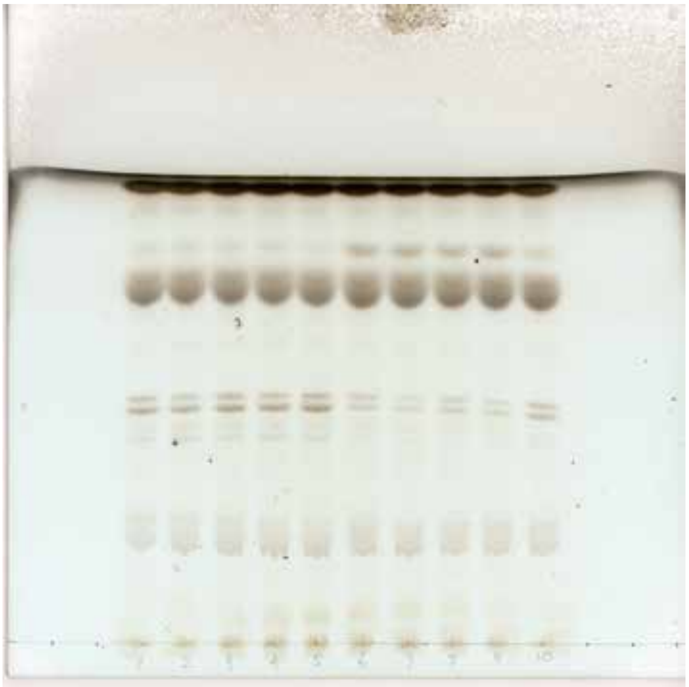

n2

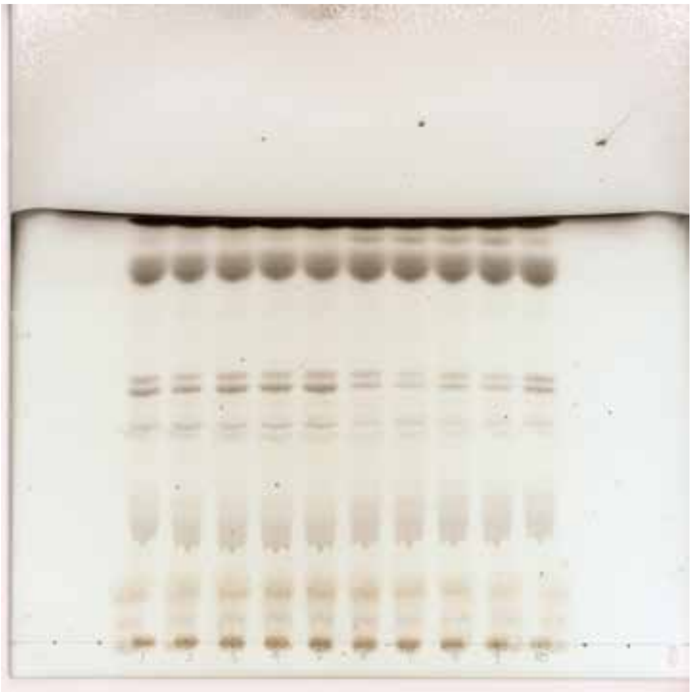

n3

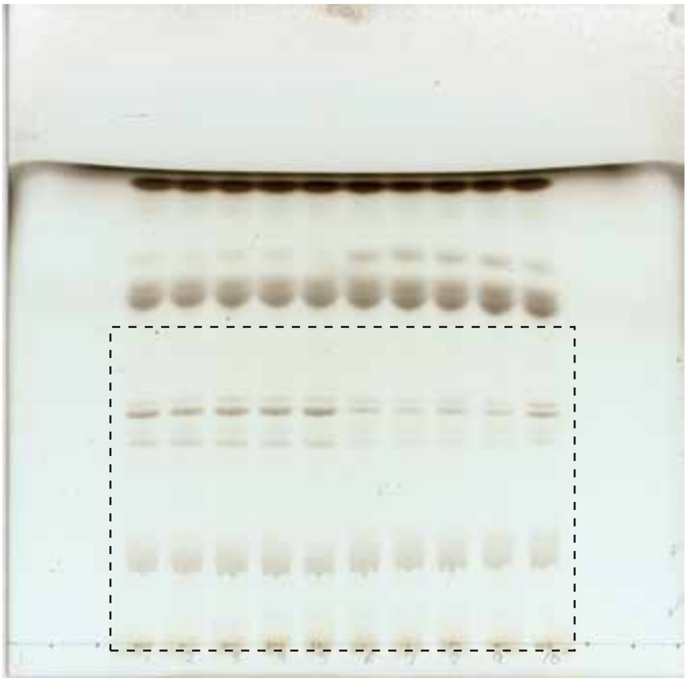

n4

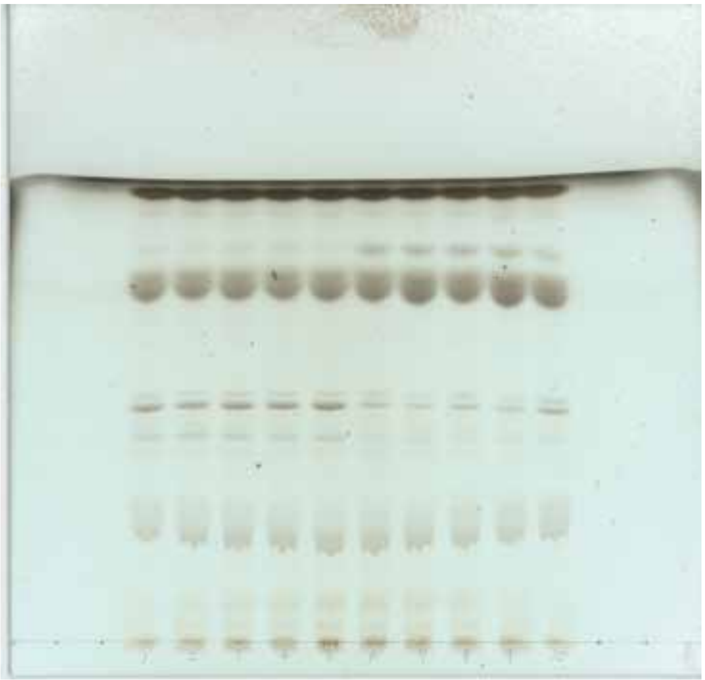

Figure 5-figure supplment source data

None

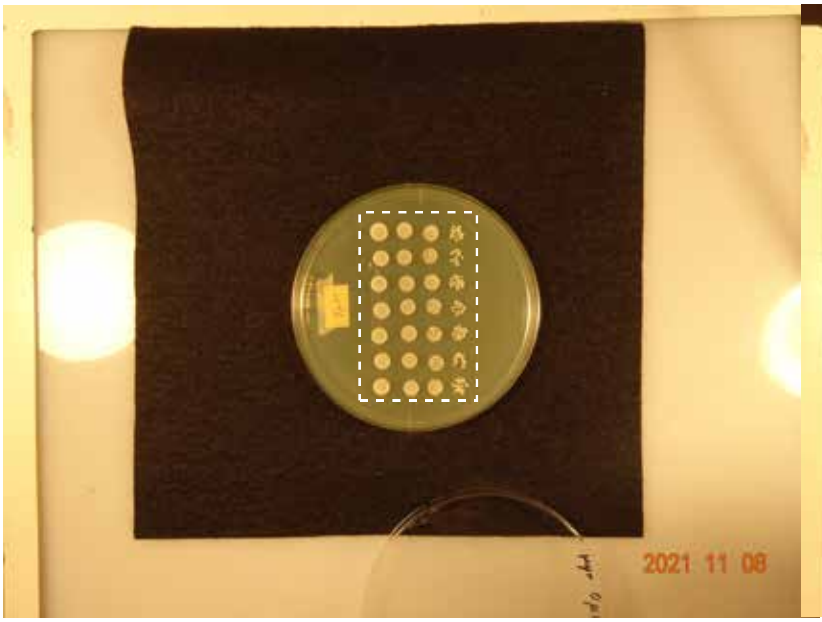


0.8 μM Myr

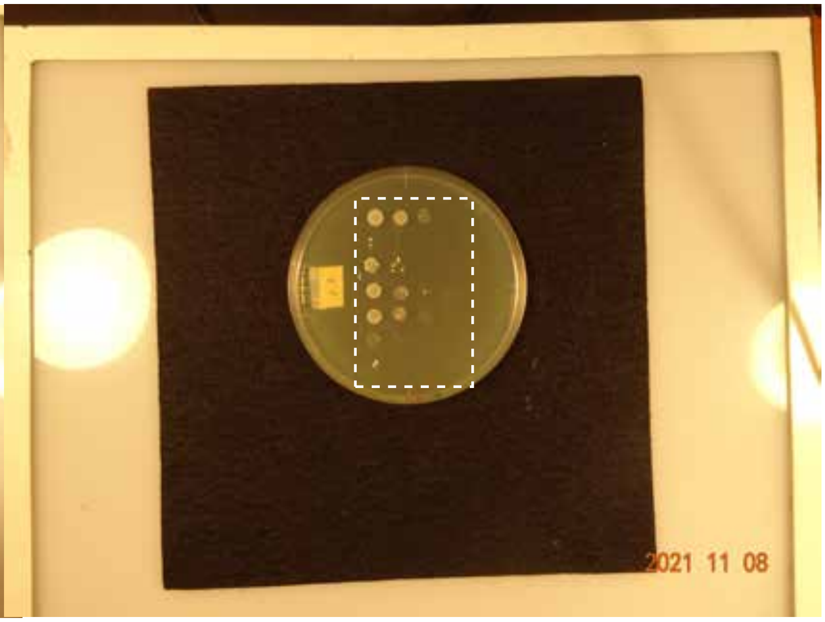


0.6 μM Myr

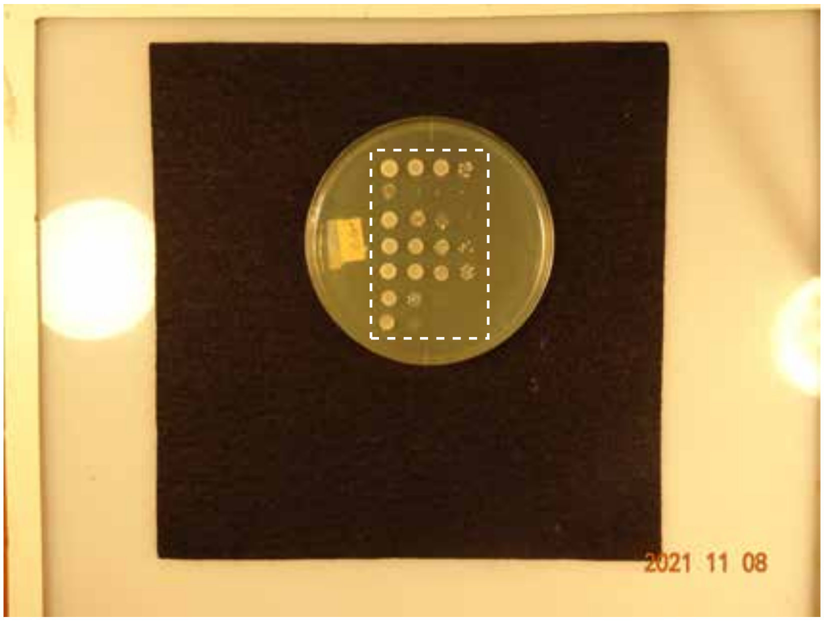

Figure 6-figure supplement source data

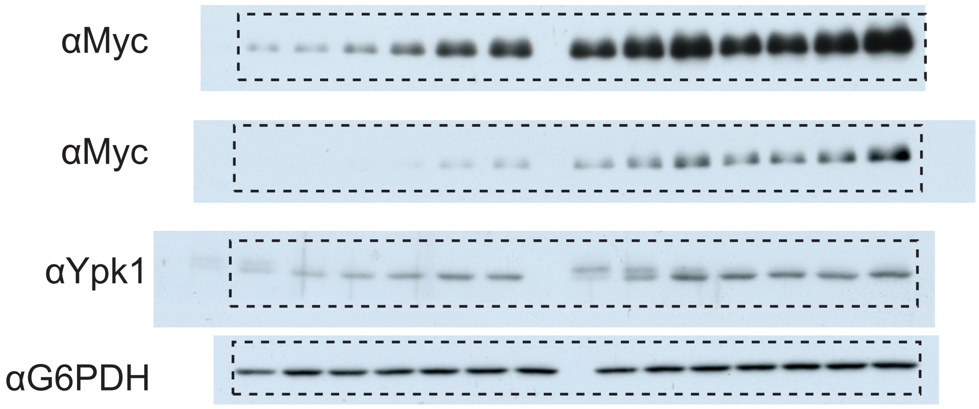

Figure 6B source data

None | 0.6 μM Myr

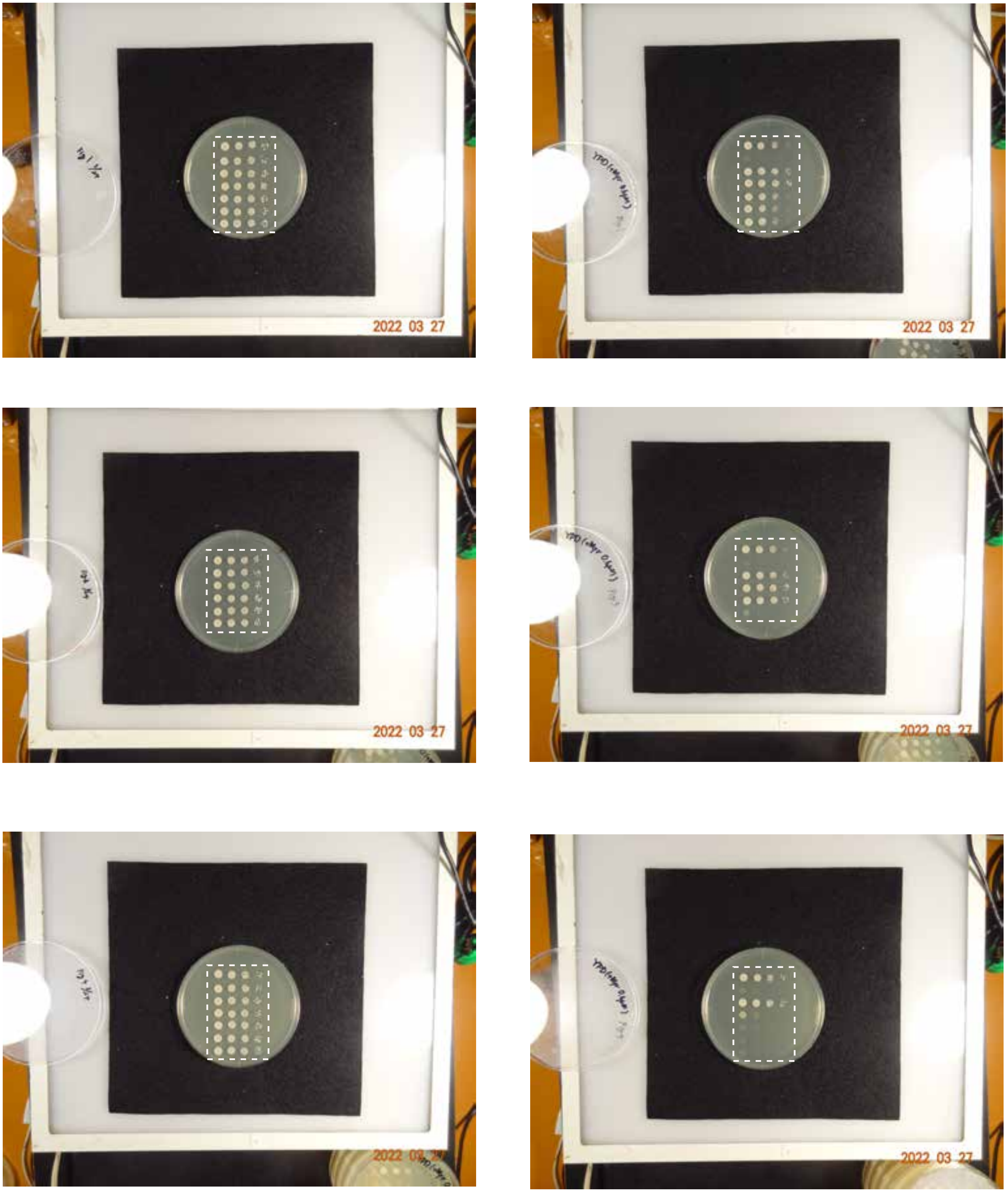

Figure 6C source data

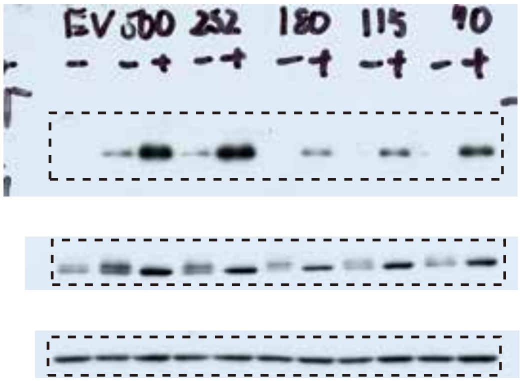


Figure 6D source data

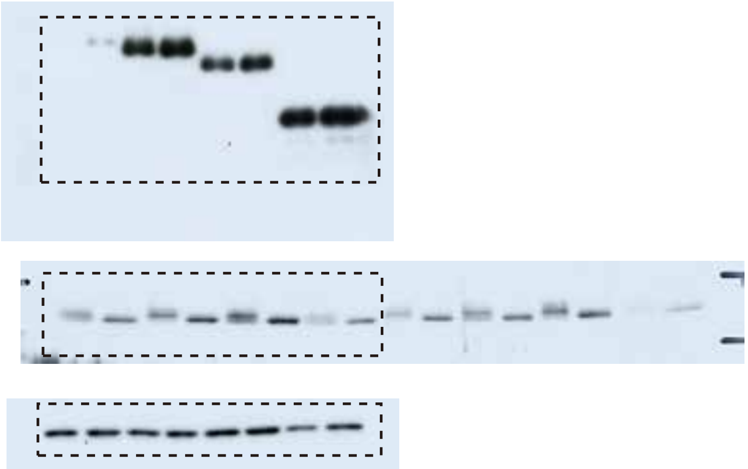

Figure 6E source data

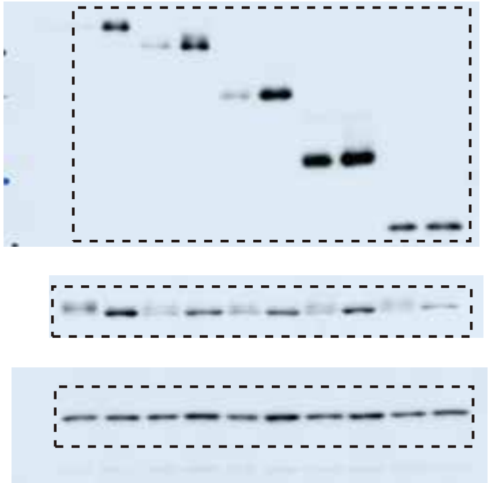

Figure 6-figure supplement source data

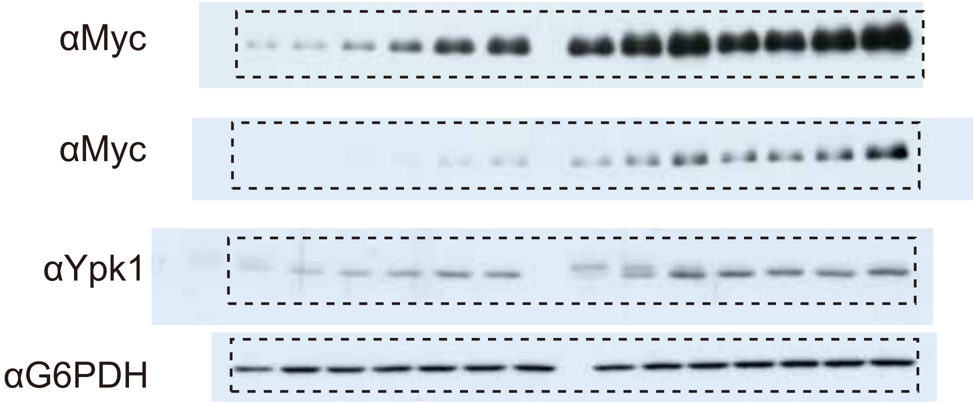

Figure 7B source data

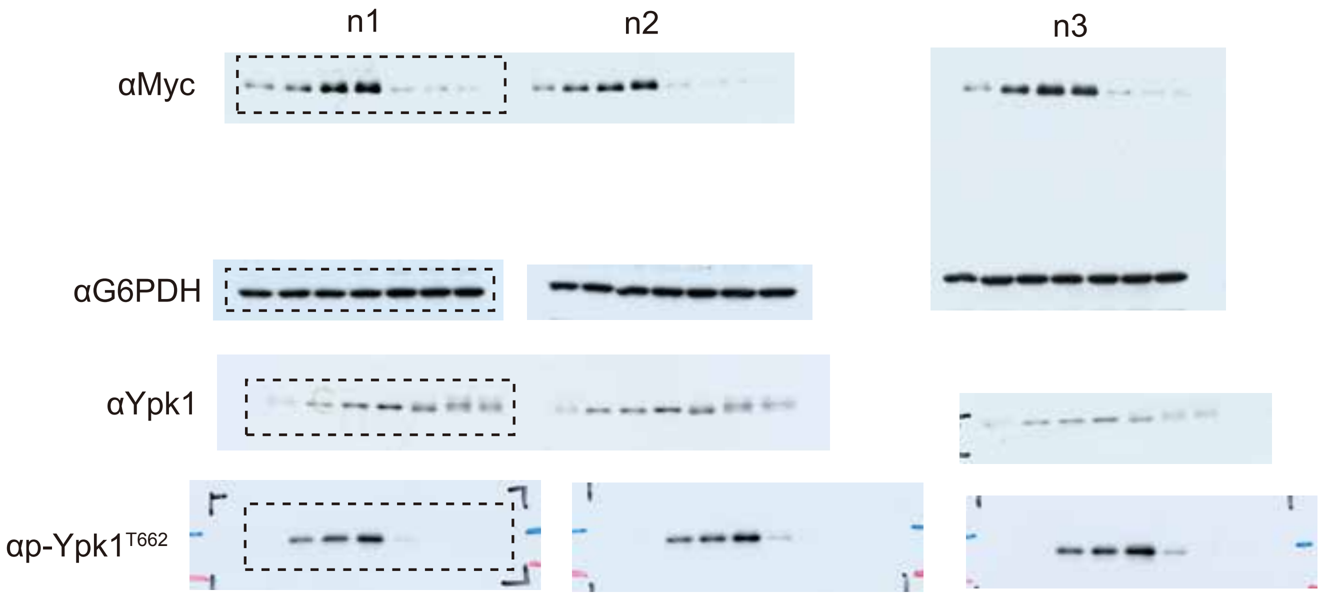


Figure 7F source data

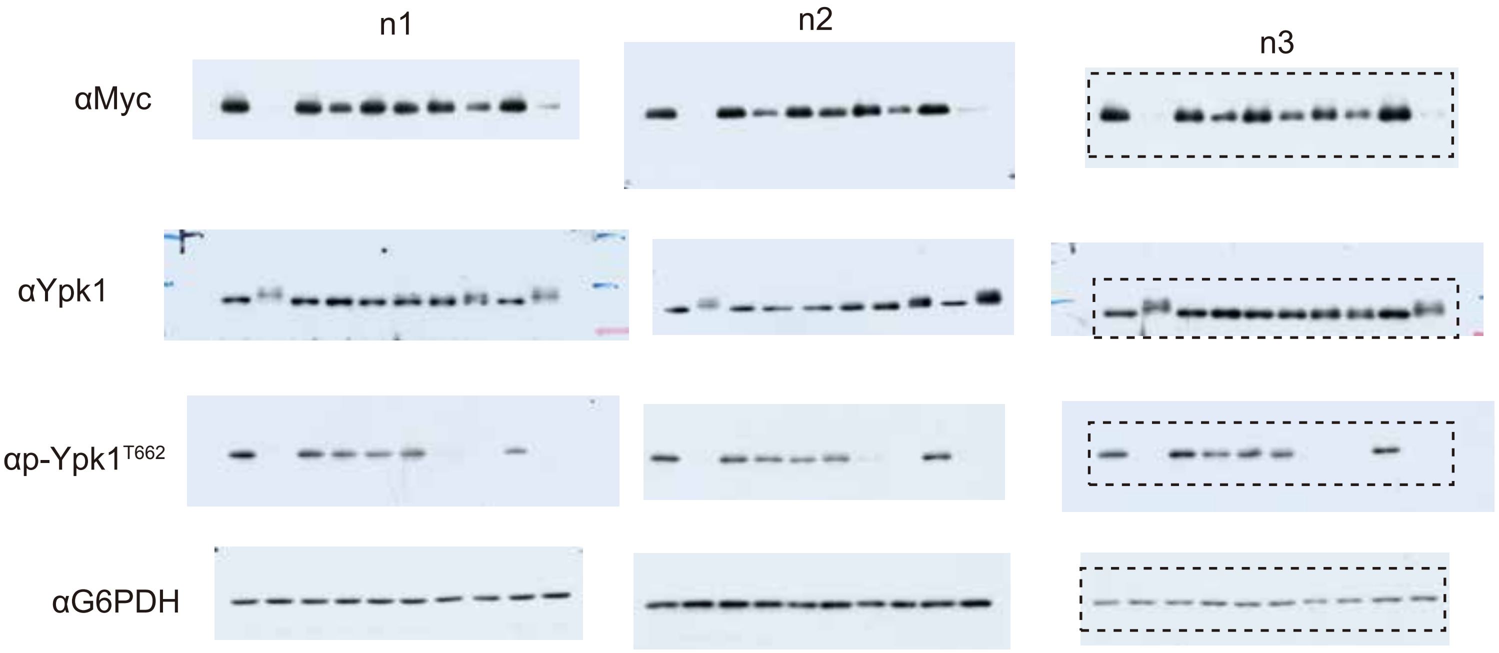

Figure 7I source data

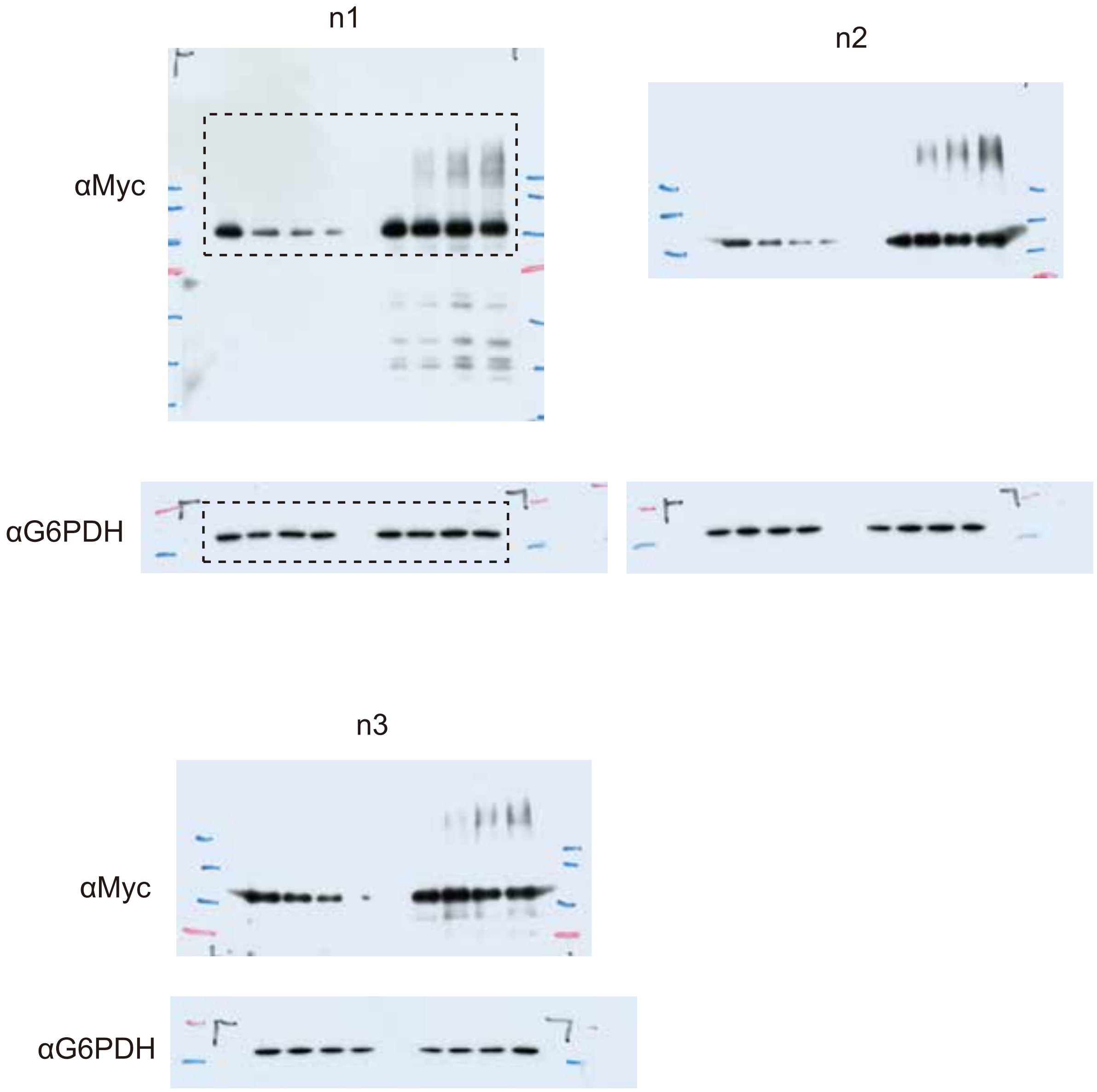

Figure 7-figure supplement source data

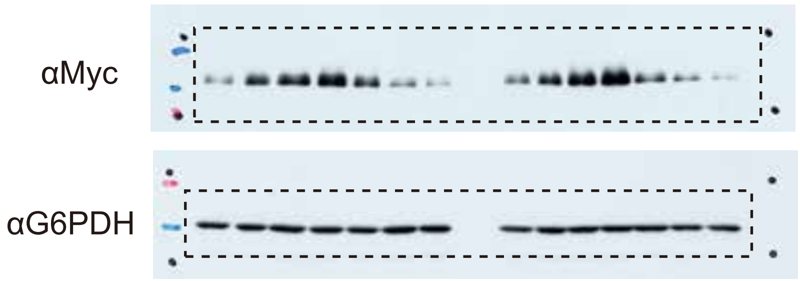

#### Figure 8A source data

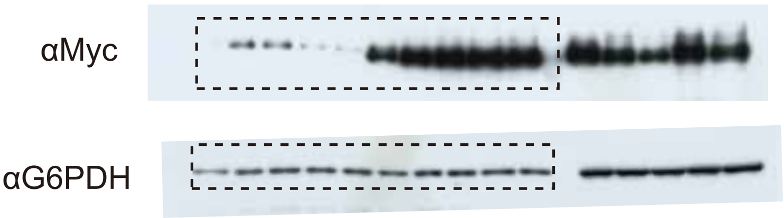


#### Figure 8C source data

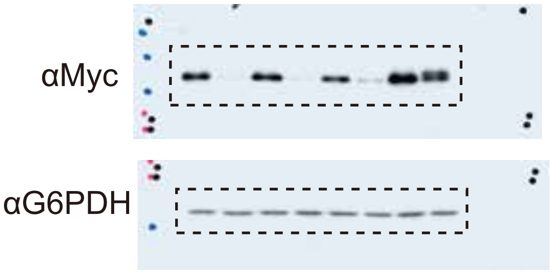


#### Figure 8D source data

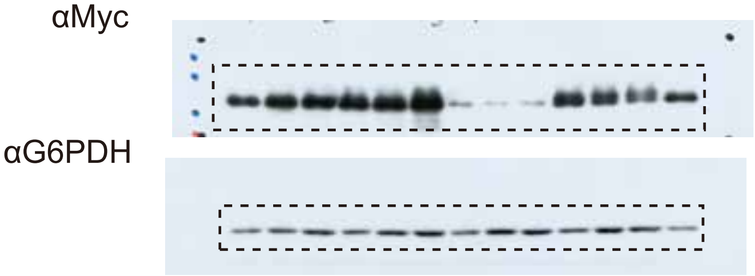


#### Figure 8F source data

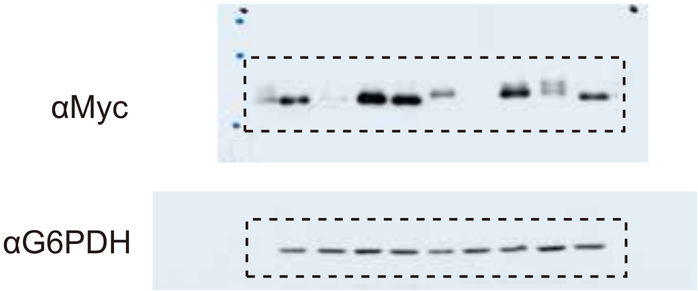


#### Figure 8G source data

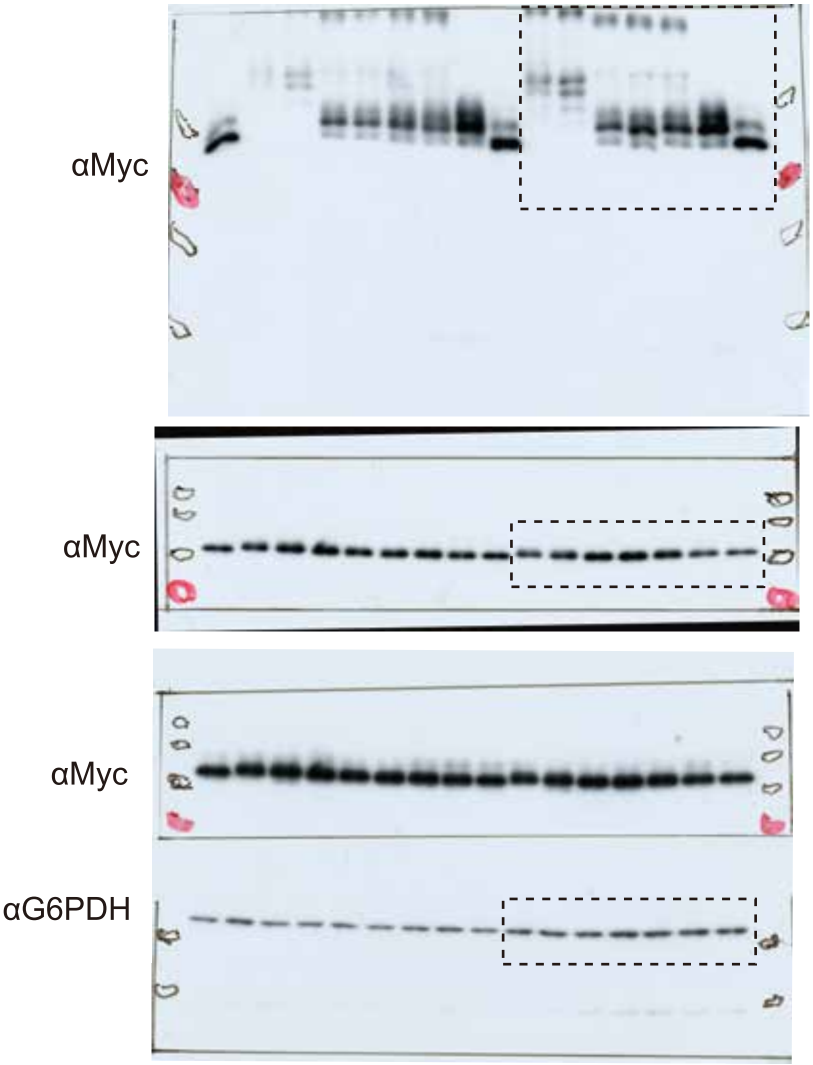


#### Figure 8H source data

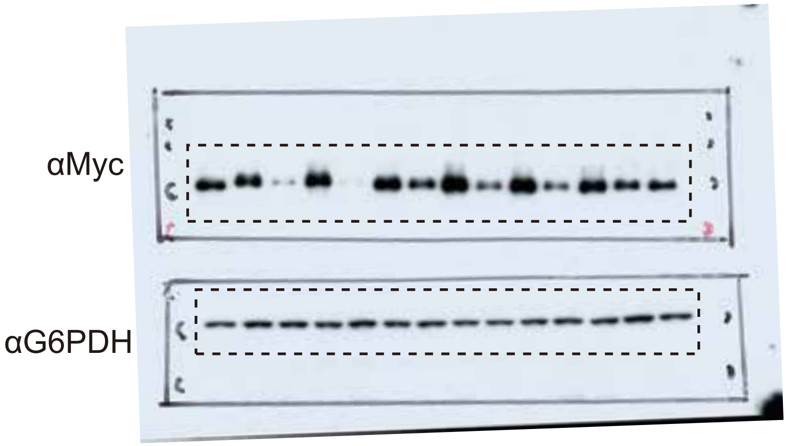


#### Figure 8I source data

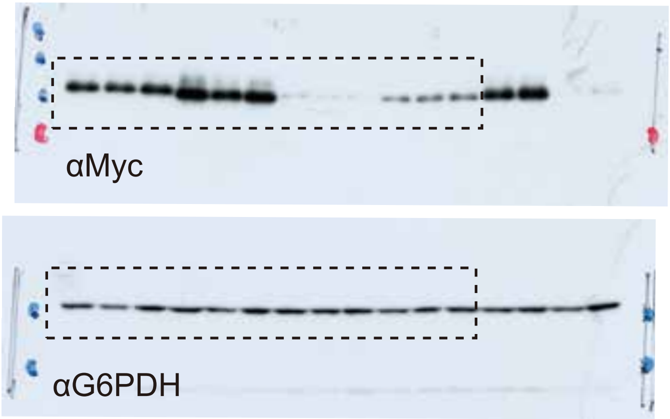

Figure 8-figure supplement A source data

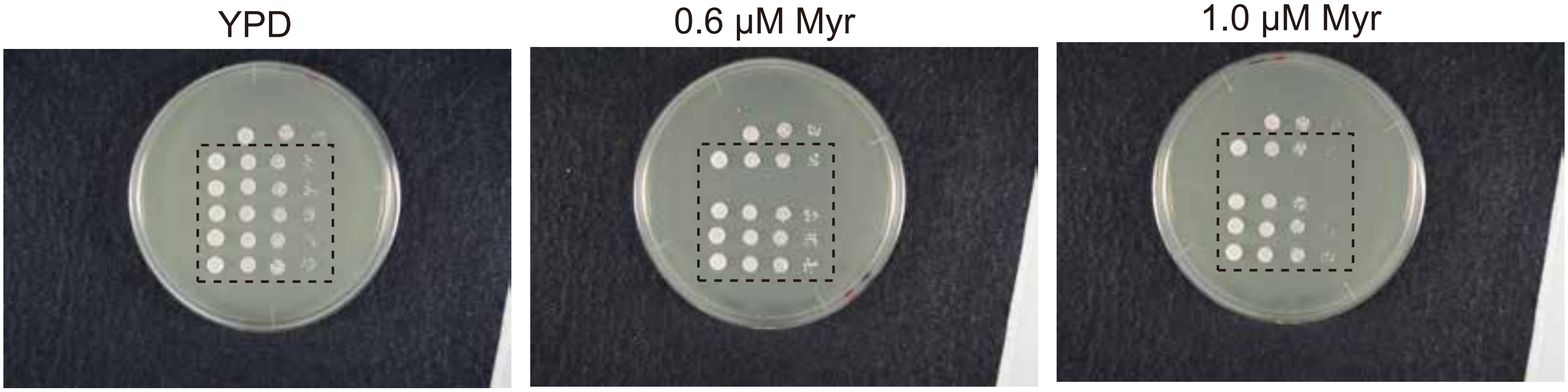


Figure 8-figure supplement C source data

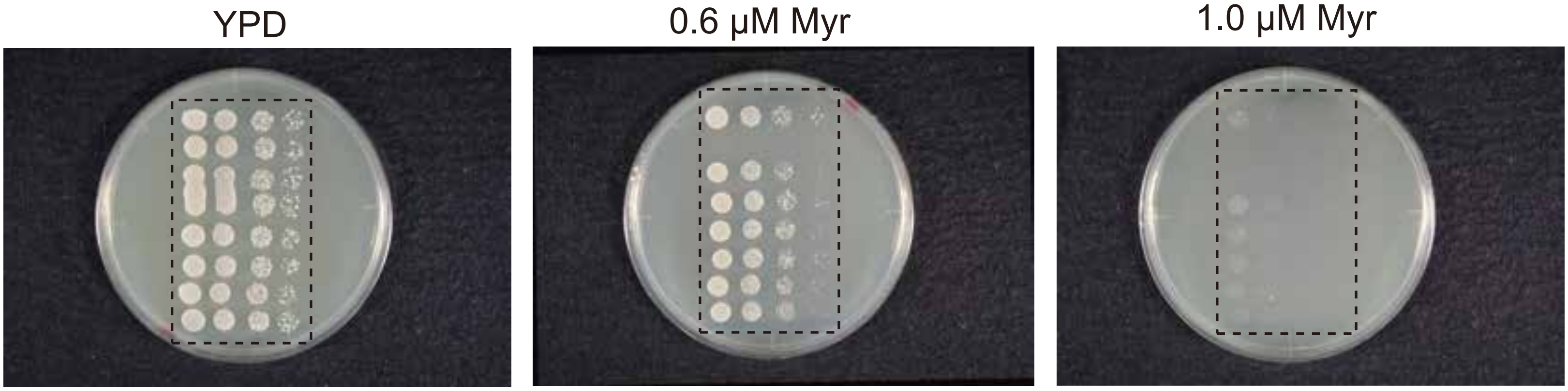
