## Supplementary material for "Regulation of sphingolipid synthesis by the C2H2 zinc finger transcription factor Com2 through ubiquitin-proteasome mediated degradation pathway": Supplmental Figure Legends

### Supplemental Figure legends

**Figure 1—figure supplement 1.** Screening to identify sphingolipid metabolism related factors that suppress Myriocin sensitivity of *lip1-1* cells. Wild-type cells and *lip1-1* cells carrying a plasmid including each suppresser gene were spotted in 10-fold serial dilution on YPD supplemented with 0.2 µM Myriocin (Myr) in the presence of Dox (20 µg/mL).

**Figure 1—figure supplement 2.** The diploid homozygous knockout cells of *MLM* genes were spotted at a 10-fold serial dilution on YPD supplemented with 0, 1.0, 2.0 µM Myr and incubated at 26 ˚C for 2-3 days.

**Figure 1—figure supplement 3.** (**A**) Wild-type, *ypk1*Δ, *com2*Δ, *hog1*Δ, *slt2*Δ and *ire1*Δ cells were spotted in 10-fold serial dilution on YPD for each condition (26 ˚C, 37 ˚C, 75 mM Acetic acid (AcOH), 50 mM Tris-HCl pH 8.0, 1 M NaCl, 0.25 µg/mL Tunicamycin (TN), 20 µg/mL Calcofluor white (CFW), 0.0150% SDS, 50 ng/mL Miconazole, 0.6 µM / 1.0 µM Myr, and 75 ng/mL AbA. (**B**) Detailed description of each step of the *de novo* sterol biosynthetic pathway in the yeast *Saccharomyces cerevisiae* and its experimental mechanism of action. The pathway and proteins responsible for the synthesis of yeast sterols are shown. (**C**) The indicated cells were spotted in 10-fold serial dilutions onto YPD supplemented with 10 µM, 50 µM, and 100 µM Terbinafine. (**E**, **F**) Wild-type cells were grown to mid-log phase in SD liquid medium and separately treated with 1.0 µM Myr, 50 µM Terbinafine, or 50 ng/ml Miconazole for 3 hrs at 26 ˚C, after which lysates were separated by SDS-PAGE and immunoblotted with anti-Myc or anti-G6PDH antibodies to detect Com2-13Myc or G6PDH (loading control), respectively. The amount of Com2-13Myc in wild-type cells (without Myr) was normalized to the amount of G6PDH, and each data was set as 100%. Data represents the mean ± SD of three independent experiments. Statistical significance was determined by one-way ANOVA followed by Tukey's multiple comparison test (****$p$<0.0001, ns: not significant).

**Figure 4—figure supplement 1.** *In silico* analysis to identify the candidate genes of the Com2 dependent expression in response to reduced sphingolipids. Schematic representation of the workflow for the bioinformatic approach towards identifying Com2 targets from the *S. cerevisiae* genome using Yeast Genome Pattern Matching (https://www.yeastgenome.org/nph-patmatch).

**Figure 4—figure supplement 2.** Raw immunoblot data used for the quantification shown in Figure 4E. $P_{\text{tet-off}}$-*GFP-COM2* cells were grown to mid-log phase in SD liquid medium in the presence (+) or absence (−) of doxycycline (Dox) and then treated with 1.0 μM Myr for 2 hrs. Cell lysates from six independent experiments were subjected to SDS-PAGE followed by immunoblotting with anti-Ypk1, anti-Myc, or anti-G6PDH antibodies to detect Ypk1, Lcb1-13Myc, or G6PDH (loading control), respectively.

**Figure 5—figure supplement.** Evaluation of myriocin sensitivity in *YPK1* and *LCB1* promoter CBS-deletion mutants. (**A**) Wild-type, *com2*Δ, $P_{YPK1}$-Δ*CBS*, $P_{LCB1}$-Δ*CBS*, or $P_{YPK1}$-Δ*CBS*/$P_{LCB1}$-Δ*CBS* double-mutant strains were spotted as 10-fold serial dilutions (starting at $OD_{600}$=1) onto YPD agar plates containing myriocin (0, 0.6, or 0.8 μM) and incubated at 26°C for 2-3 days to assess myriocin sensitivity. (**B**) The same strains were cultured in YPD liquid medium containing myriocin (0 or 0.4 μM) in microplates, and growth curves were obtained by measuring $OD_{600}$ every 30 min. (**C**) $OD_{600nm}$ values at 20 hrs of cultivation in the presence of myriocin (0 or 0.4 μM) were quantified (n=3). Statistical significance was determined by one-way ANOVA followed by Tukey's multiple comparison test (***p*<0.01, *****p*<0.0001, ns: no significant difference).

**Figure 6—figure supplement.** Myriocin-dependent upregulation of Com2 expression is independent of the promoter. $P_{\text{tet-off}}$-*13Myc-COM2* cells were grown to logarithmic phase in SD liquid medium in the presence (+Dox) or absence (−Dox) of doxycycline. Myriocin was then added to a final concentration of 1 μM, and protein extracts were prepared at the indicated time points. The expression levels of 13Myc-Com2 and Ypk1 were analyzed by immunoblotting. G6PDH was used as a loading control.

**Figure 7—figure supplement.** Wild-type and *atg5*Δ cells expressing Com2-13Myc were treated with 1.0 μM Myr for 3 hrs., followed by the addition of 10 μM PHS and further incubation for 3 hrs. Samples were collected at the indicated time points, and Com2-13Myc protein levels were analyzed by immunoblotting. G6PDH was used as a loading control.

**Figure 8—figure supplement.** (**A**) *com2*Δ cells expressing wild-type Com2, an empty vector, or Com2 mutants carrying substitutions at the predicted ubiquitination sites were spotted in 10-fold serial dilutions onto YPD medium containing the indicated concentrations of myriocin (Myr). (**B**) Analysis of putative phosphorylation sites in Com2. Phosphorylation sites of Com2 annotated in the *Saccharomyces* Genome Database (SGD), putative Ypk1 phosphorylation sites reported by Muir et al., and AGC kinase–dependent phosphorylation sites in Com2 predicted using the GPS web server (http://gps.biocuckoo.org/online.php) are shown. Serine/threonine residues mutated to alanine in the stepwise phosphorylation-site mutants are indicated. (**C**) Wild-type or *com2*Δ cells transformed with plasmids expressing wild-type Com2 or the indicated phosphorylation-site mutants were spotted in 10-fold serial dilutions onto YPD medium containing the indicated concentrations of Myr.
